# Physiological diversity and the assembly of nitrifying communities in aerobic granular sludge

**DOI:** 10.64898/2026.11.12.766174

**Authors:** Eloi Martinez-Rabert, Rebeca Gonzalez-Cabaleiro

## Abstract

One-stage partial nitritation/anammox systems provide a cost-effective and efficient approach to nitrogen removal, but its wide application for sewage treatment remains a challenge. Four key ecological dynamics determine its success: *i*) the protection of anammox bacteria from oxygen exposure by nitrifiers, *ii*) the inhibition of nitrite-oxidizing bacteria through residual ammonia accumulation, *iii*) the competition for oxygen among nitrifiers, and *iv*) the competition for nitrite between nitrite-oxidizing bacteria and anammox bacteria. Using a comprehensive Individual-based Model, we simulate the ecological interactions among microbial communities under relevant operational and environmental conditions, emphasizing the role of nitrite-oxidizing bacteria diversity by differentiating *Nitrobacter*, *Nitrospira* and *Nitrotoga* genera. Our simulation experiments identify and quantify the fundamental constraints when optimizing one-stage partial nitritation/anammox systems, particularly the challenge of suppressing nitrite-oxidizing bacteria while maintaining a strong anammox activity. Notably, our results show that the previously observed stratification between ammonia- and nitrite-oxidizing communities, does not indicate optimal nitrogen process partitioning. These findings highlight the limitations of relying solely on operational design to control community structure, as this approach may fail to ensure stable and efficient nitrogen removal across diverse conditions in one-stage partial nitritation/anammox systems.

**Author Summary:** Nitrogen pollution by ammonia, nitrite and nitrate, is the biggest driver of biodiversity loss, ecosystem degradation, and main responsible of algal blooms and eutrophication in marine and freshwater ecosystems. Hence, the development of more efficient and cost-effective sewage treatment systems is crucial, being the combination of partial nitritation (partial conversion of ammonia to nitrite) and anammox processes (conversion of ammonia and nitrite to nitrogen gas) in a single reactor the most promising one. However, its application for sewage treatment remains a challenge due to the effective suppression of microbes consuming nitrite, known as nitrite-oxidizing bacteria. Here, we have studied the relevance of the diversity of nitrite-oxidizing bacteria on the performance of partial nitritation/anammox process through mathematical modelling. We have been able to identify and quantify the fundamental constraints when optimizing one-stage partial nitritation/anammox systems due to the challenge of suppressing nitrite-oxidizing bacteria while maintaining a strong anammox activity. These results show that the diversity of nitrite-oxidizing bacteria play a crucial role on the optimization and maintenance of a stable partial nitritation is fundamental to optimize and stabilize the one-stage partial nitritation and anammox process.

## Introduction

Over the recent few decades, our understanding of the ecology of the nitrification process, the biological oxidation of ammonia to nitrate via nitrite, has advanced thanks to three fundamental discoveries: *i*) bacteria able to perform anaerobic ammonia oxidation (with the establishment of the anammox process) [1]; *ii*) the archaeal ammonia oxidation [2, 3] and *iii*) the complete ammonia oxidation by *Nitrospira* (comammox process) under aerobic and hypoxic conditions [4, 5]. As a result of these discoveries, more sustainable and economic technologies for nitrogen removal than classical nitrification-denitrification processes have been developed. Among these, the combination of partial nitritation (PN, aerobic oxidation of ammonia to nitrite, 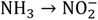) and anammox process (AMX, anaerobic ammonia oxidation with concomitant reduction of nitrite to nitrogen gas, 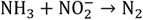), known as partial nitritation/anammox (PN/A) process [6], is the one that has been more successful in its industrial implementation [7]. In the PN/A process, the oxygen demand is reduced up to 60% compared with traditional nitrification-denitrification, additionally eliminating any organic carbon requirement [7]. However, the effective application of PN/A, as of all anammox-based processes, is limited to specifically treat ammonia-rich sewage, like effluents from anaerobic sludge digesters [8].

The PN/A process is applied in two different configurations: one-stage (single aerobic reactor) and two-stage (two reactors, one aerobic for PN and other anaerobic for anammox process). Although the two-stage configuration PN/A process has a higher efficiency and easier operational management, the lower cost of one-stage PN/A has made this technology the preferred solution [7, 9]. The key challenge for the large-scale application of the PN/A process to treat municipal sewage, especially in the one-stage configuration, is on the difficulty of long-term suppression of nitrite-oxidizing bacteria (NOB) while the maintenance of adequate activity of anammox bacteria [10]. Anammox bacteria transform NO_2_^−^ directly to N_2_ gas, meanwhile NOB transforms NO_2_^−^ to NO_3_^−^, which in a PN/A configuration remains accumulated in the water. For an effective suppression of NOB, a combination of low oxygen concentration, accumulation of ammonia, and stratification of AOB and NOB inside a granule is identified as main requirements [11]. However, the maintenance of a stable NOB suppression is challenging, supposedly because of the wide physiological diversity of NOB [12]. Examples are the re-emergence of active *Nitrobacter* after the successful suppression of *Nitrospira* and *Nitrobacter* in Poot *et al.* (2016) [11], the shift of NOB populations from *Nitrospira* to *Nitrotoga* [13], or the persistence of *Nitrobacter* under high concentration of residual ammonia [14].

The success of one-stage PN/A process is ruled by four necessary ecological conditions (Fig 1): *i*) the protection of anammox bacteria against oxygen; *ii*) the inhibition of NOB activity by residual ammonia accumulation; *iii*) the competition for oxygen between AOB and NOB; and *iv*) the competition for nitrite between NOB and anammox bacteria. Not all NOB populations are influenced by these ecological dynamics in the same way. *Nitrobacter* and *Nitrotoga* are the populations of NOB with the highest tolerance to free ammonia (*ii*) [15, 16], although it has been found that some *Nitrospira* species (such as *Nitrospira japonica*) may also have a significant ammonia tolerance [17].

**Fig 1.**
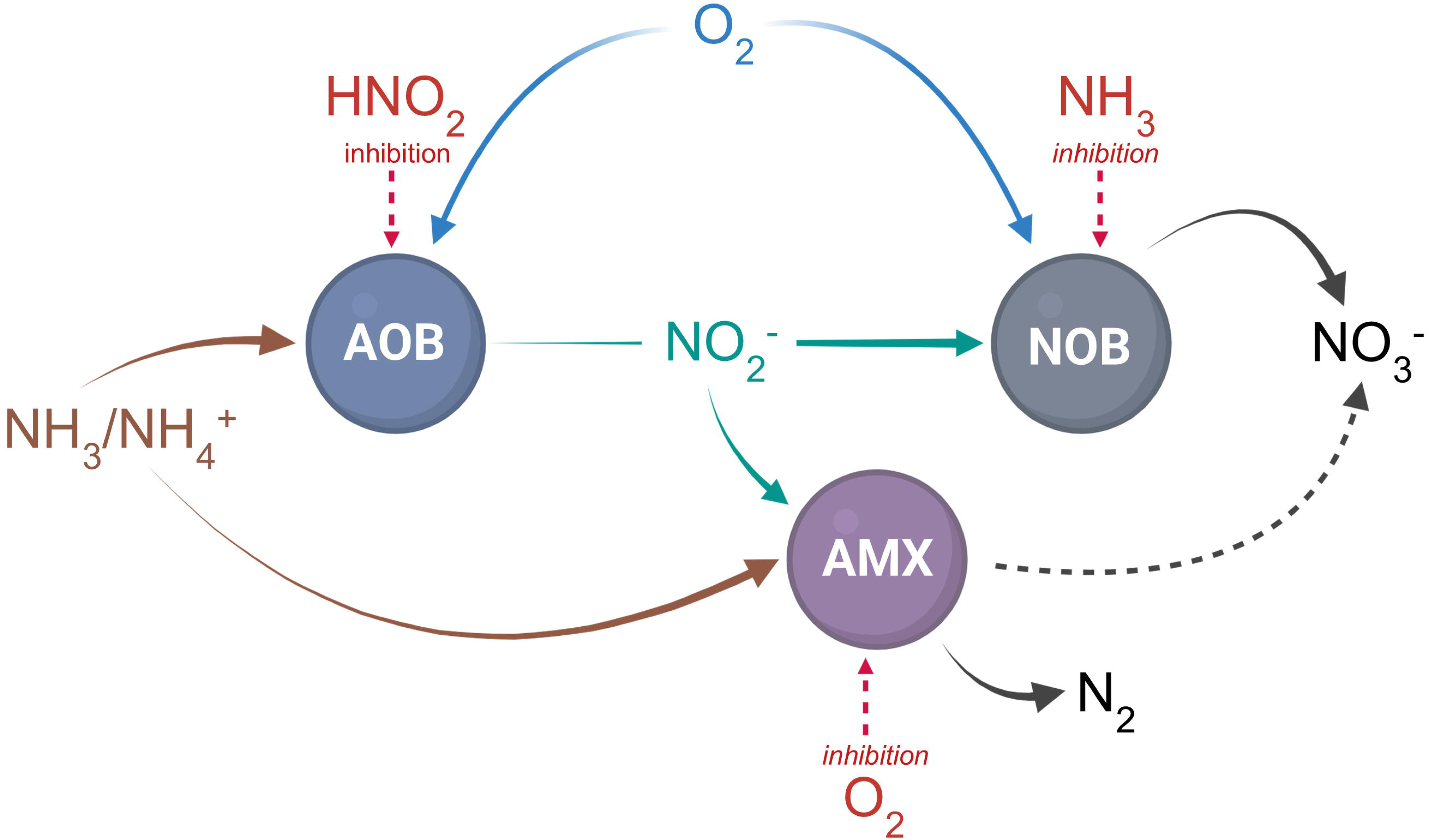
Ecological impact of nitrification on anammox activity. Ecological network of nitrification associated with anammox process. Brown arrows represent the competition for ammonia, blue arrows represent the competition for oxygen, green arrows represent the competition for nitrite, and red lines represent inhibition effects.

The substrate affinity and growth yield seem to be the most important physiological parameters to consider when trying to control the competition for oxygen and nitrite among populations. Considering the NOB diversity and the sensitivity analysis presented in Hao *et al*. 2002 [18], *Nitrospira* would be the stronger competitor to anammox bacteria in one-stage PN/A because its affinity for nitrite is comparable to the one observed for anammox bacteria (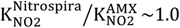; Fig 2A), and its affinity for oxygen tends to be higher than the observed affinity for AOB (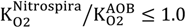; Fig 2B). At the same time, nitrite oxidation has a higher growth yield on oxygen than ammonia oxidation (4.84×10^−2^ C-mol biomass/mol-O_2_ for nitrite oxidation; 2.72×10^−2^ C-mol biomass/mol-O_2_ for ammonia oxidation), hindering the suppression of NOB activity by oxygen competition [19].

**Fig 2.**
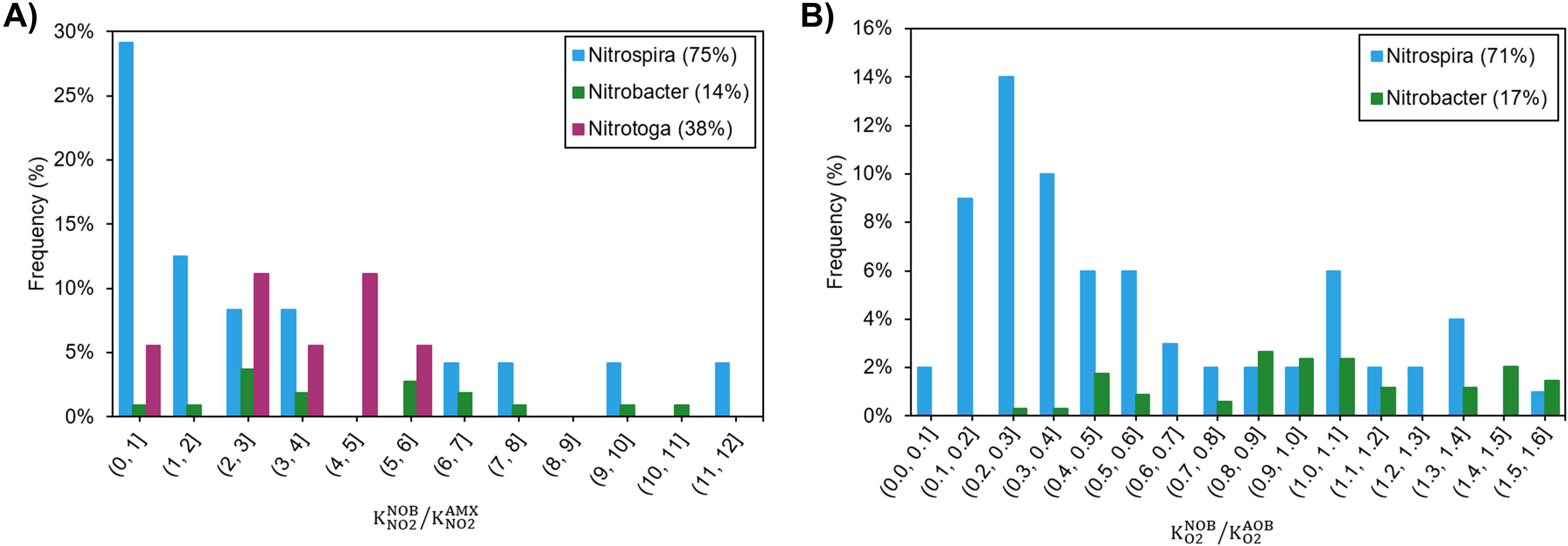
Kinetic analysis of competitions between AOB, NOB and anammox bacteria. Histograms show the frequency of **A)** 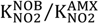 ratio and **B)** 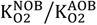 ratio, differentiating NOB population between *Nitrobacter*, *Nitrospira* and *Nitrotoga*. The range of 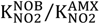 and 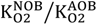 ratios shown here are the same as that presented in Hao, Heijnen [18]. Numbers in parenthesis in the legend show the total percentage of the ratio included in the histogram. To create the histograms, the kinetic database of Martinez-Rabert, E., *et al* 2022 [20] was used.

Several modelling studies have been conducted to figure out the main dynamics and mechanisms that control the suppression of NOB in partial nitritation, contributing to our understanding of nitrification and partial nitritation [18, 21–27]. In this work, we build up on this knowledge to study the coexistence of different NOB populations (*Nitrobacter*, *Nitrospira* and *Nitrotoga* genera), argued as one of the main reasons challenging NOB suppression.

A multiscale model is used as framework to simulate the activity of individual bacteria to investigate the influence of the diversity of NOB populations on a nitrogen removal process in which nitrification, nitritation and PN/A processes take place in the same aerobic granule (representing a one-stage PN/A process) under specific conditions. In this model, we have defined the microbial growth of the three most common populations of NOB in wastewater treatment plants (WWTP) – *Nitrobacter*, *Nitrospira* and *Nitrotoga* – by applying kinetic parameters coming from an extensive database [20]. The rigorous physiological definition of these populations makes the model capable of predicting when the persistence of NOB populations influences the nitrogen conversion process partitioning and how to control it, overall drawing the limits of, for example, the one-stage PN/A process application for municipal sewage treatment.

## Results and Discussion

The Individual-based Model framework is configured to simulate a continuous bioreactor with controlled setpoints of ammonia and oxygen. The concentration of ammonia in the bulk liquid is set to 20.0 mg/L, which ensures the influence of residual ammonia concentration on NOB repression [11]. In the continuous reactor, nitrifying granules evolve. They are formed by the growth of aerobic nitrifiers which use nitrogen forms as electron donors and compete among them for the O_2_ available (as electron acceptor). With their activity, they deplete the available O_2_ generating an anaerobic granule core. In this anaerobic core, when enough nutrients are available, anammox bacteria can thrive. Using this mathematical framework, the effects of oxygen concentration in the liquid bulk (an operational parameter of the system and therefore controllable), and temperature (a non-controllable environmental condition) are analysed predicting the composition of the microbial community within the granules.

With the objective to study to which extent the physiological diversity of NOB controls the nitrifying community assembly in aerobic granules, three specific genera are defined in the model, *Nitrobacter*, *Nitrotoga* and *Nitrospira*, by using specific kinetic parameters for growth associated with each genus. For *Nitrospira*, the most diverse NOB genus [12], a sensitivity analysis of kinetic parameters (oxygen affinity and ammonia tolerance) is presented. S1 Table summarises the operational and environmental parameters of the base case study.

### Influence of dissolved oxygen

To study the influence of O_2_ over the nitrifying community at 20°C, six concentrations are evaluated: 0.1 mg/L (S1 Fig), 0.25 mg/L, 0.5 mg/L, 1.0 mg/L and 3.0 mg/L (Fig 3), and 6.0 mg/L (S2 Fig). At 0.1 mg-O_2_/L (a concentration that is slightly lower than the oxygen affinity of AOB; S2 Table), nitrite oxidation was not successfully supressed observing accumulation of nitrate at steady state. In these conditions, anammox bacteria performed substantially (63.14% ± 0.64%) (S1 Fig). Increasing the oxygen concentration to 0.25 mg/L and 0.5 mg/L (Figs 3A and 3B, respectively), the relative abundance of *Nitrospira* (only aerobic populations considered) was reduced from 27.7% ± 1.5% (0.1 mg-O_2_/L) to 19.4% ± 12.4% and 7.7% ± 3.8% (0.25 mg-O_2_/L and 0.5 mg-O_2_/L, respectively). Only at 1.0 mg-O_2_/L NOB population was successfully supressed as *Nitrospira* was outcompeted by AOB (Fig 3C), which is consistent with the first stage of operation reported by Poot *et al.* (2016) [11]. When oxygen concentration was lower than 1.0 mg-O_2_/L, *Nitrospira* closely competed with AOB, because its higher growth yield and affinity on oxygen (S2 Table), remaining at the outside of the granule (Figs 3A and 3B). When oxygen concentration was higher than 1.0 mg-O_2_/L, *Nitrospira* remained active generating a *layered stratification* [28] of AOB and NOB populations (Fig 3D).

**Fig 3.**
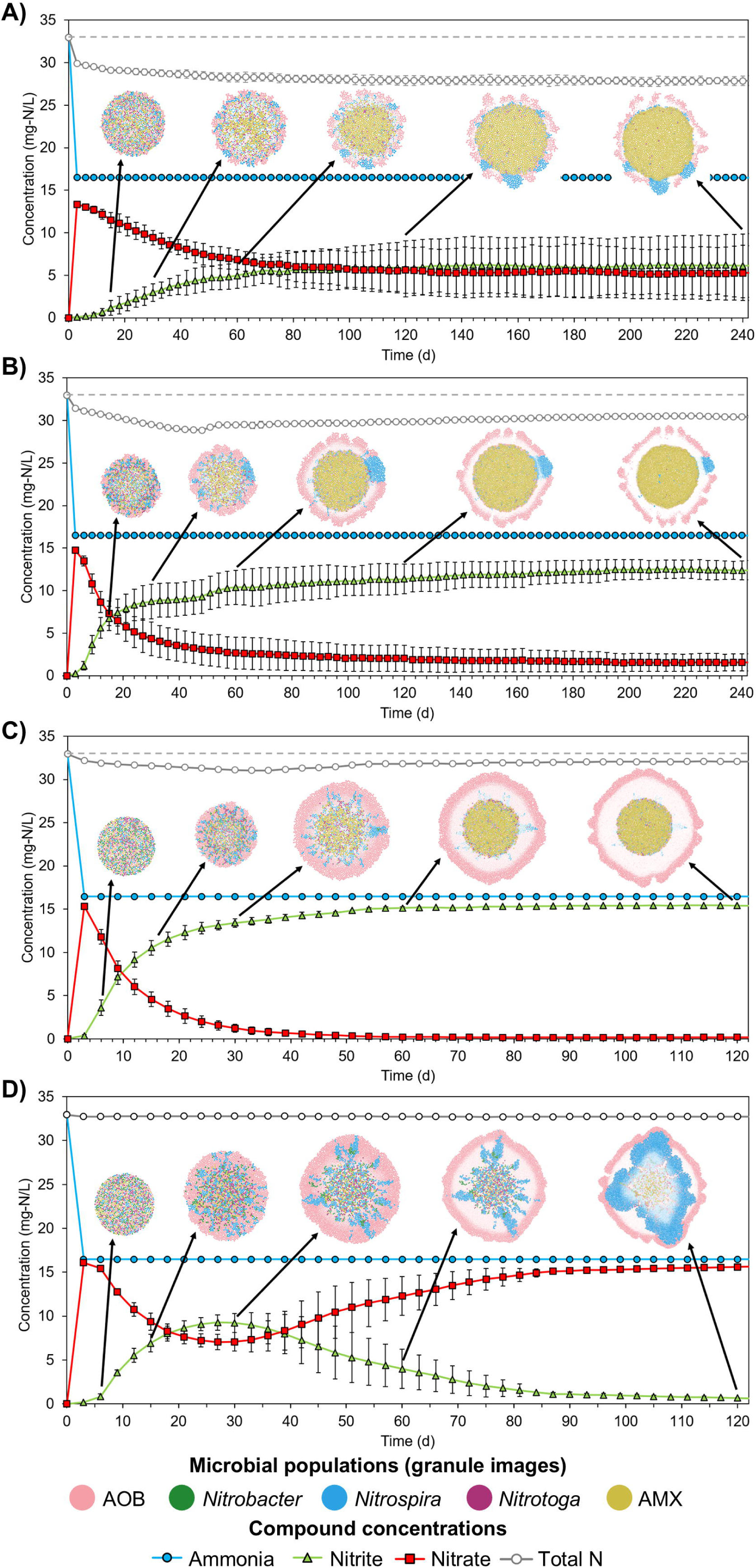
Influence of dissolved oxygen concentration on the community distribution and activity. **A)** [O_2_] = 0.25 mg/L (*n* = 3). **B)** [O_2_] = 0.5 mg/L (*n* = 3). **C)** [O_2_] = 1.0 mg/L (*n* = 3). **D)** [O_2_] = 3.0 mg/L (*n* = 3). Temperature was set to 20°C. Arrows indicate the time at which the images of granules were taken (15d, 30d, 60d, 120d, 240d for <1.0 mg-O_2_/L; 6d, 15d, 30d, 60d, 120d for ≥1.0 mg-O_2_/L). Granule images do not follow any scale. Granule replicates can be found in S3 Fig and S4 Fig.

The *layered stratification* (AOB/NOB stratification) has been identified in literature as a requirement for a successful PN/A operation [11, 21, 23]. However, in the performed simulations, nearly all the nitrite in the bulk liquid was converted to nitrate by NOB, achieving complete nitrification, when AOB/NOB stratification was established. Only during the transitory phases, in which no stratification of AOB and NOB was observed, nitrite was accumulated (3.0 mg-O_2_/L and 6.0 mg-O_2_/L; Fig 3D and S2 Fig, respectively). Considering this, AOB/NOB stratification would not correlate with effective NOB suppression and, in fact, the highest abundance of NOB was observed after the development of the *layered stratification* (S5 Fig). This is consistent with the results presented in a previous work, in which *layered stratification* was identified as the optimal spatial organization of microbial communities in which commensalism was the dominant ecological interaction [28]. Therefore, based on the simulation results, we can hypothesise that AOB/NOB stratification is an indication that NOB suppression is failing.

At 20°C, excluding the case at 0.1 mg-O_2_/L (S1 Fig), the PN/A process cannot be maintained, with total nitrification or just partial nitritation as dominant activities in the bioreactor. Anammox bacteria cannot sustain the necessary activity to transform entirely the nitrite of the system to N_2_ gas. When the tested O_2_ concentration was increased, the PN/A portion was negatively impacted (Fig 4). At 1.0 mg-O_2_/L, anammox bacteria remained active in the granule core (Fig 3C), but their contribution to nitrogen conversion was minimum (<10%, Fig 4), circumstance also observed in Poot *et al.* (2016) [11]. Under these operational conditions, the oxygen consumption by AOB is not enough to ensure an anaerobic environment at the core of the granule and anammox bacteria wash out of the system after 40 days of operation ([O_2_]_core_ > 0.01 mg/L; S6 Fig). At a setpoint of 3.0 mg-O_2_/L or higher, anammox bacteria were completely repressed (Fig 3D and S2 Fig), which can be explained by the higher concentration of O_2_ predicted at the granule core ([O_2_]_core_ > 0.76 mg/L at 3.0 mg-O_2_/L and [O_2_]_core_ > 3.75 mg/L at 6.0 mg-O_2_/L after 120d of operation).

**Fig 4.**
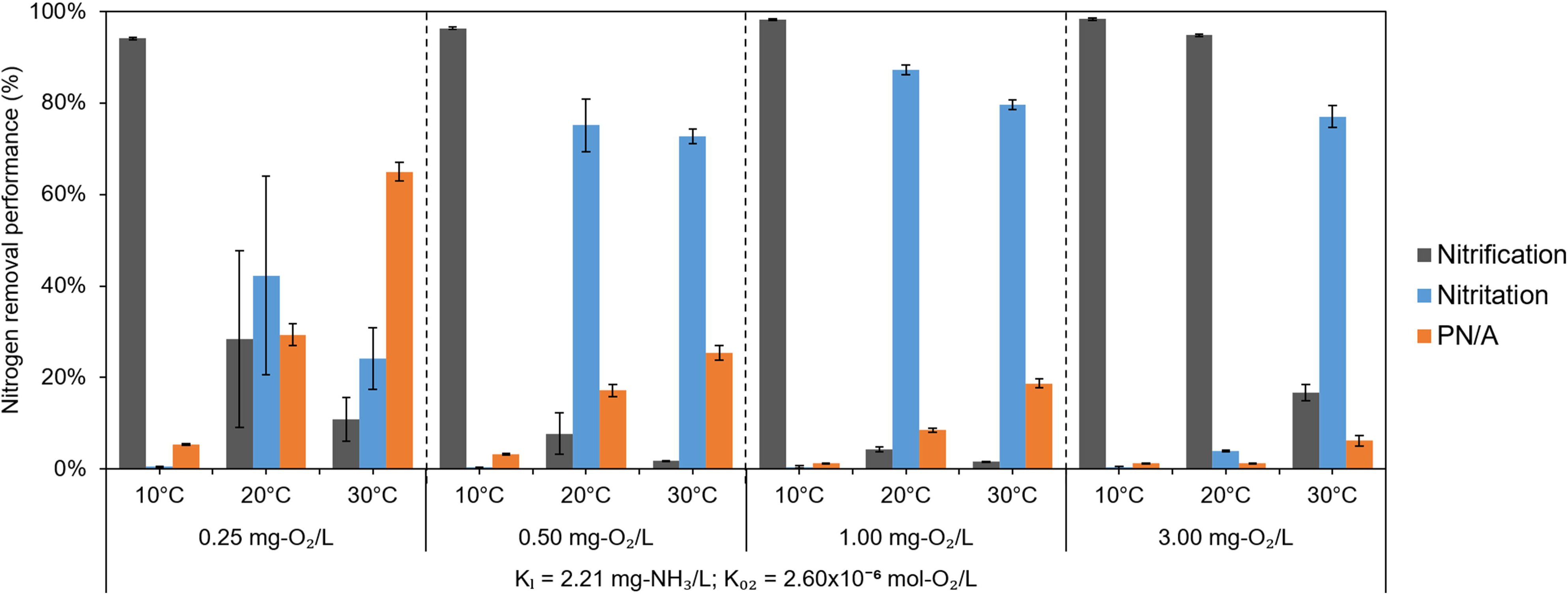
Nitrogen processes partitioning at stationary state (average of the last 10 days of operation). Simulations with the conditions of the base case (S1 Table). Maximum process portion values are included in S3 Table. Errors bars show standard deviation of *n* = 3 simulation replicates. The statistical pairwise comparison is shown in S1 Appendix.

At concentrations ≥3.0 mg-O_2_/L, transitory accumulation of nitrite is predicted. Higher O_2_ concentrations in the liquid bulk shortened the periods of nitrite accumulation. At this oxygen concentration level, nitrification dominated the nitrogen conversion process at the end of operation (>93.5%; Fig 4 and S8 Fig), with NOB populations outcompeting anammox bacteria, reducing nitrite concentrations in the granule core. Fig 3D and S2 Fig show the evolution of the community structure in the granule, where the presence of anammox bacteria slowly decayed at its core.

### Influence of temperature

The influence of temperature is analysed by simulating the base case (S1 Table) at 10°C, 20°C and 30°C. Oxygen concentration was also tested for the new temperatures using the same concentrations as in the previous section: 0.1 mg/L (S7 Fig), 0.25 – 3.0 mg/L (Fig 5) and 6.0 mg/L (S8 Fig).

**Fig 5.**
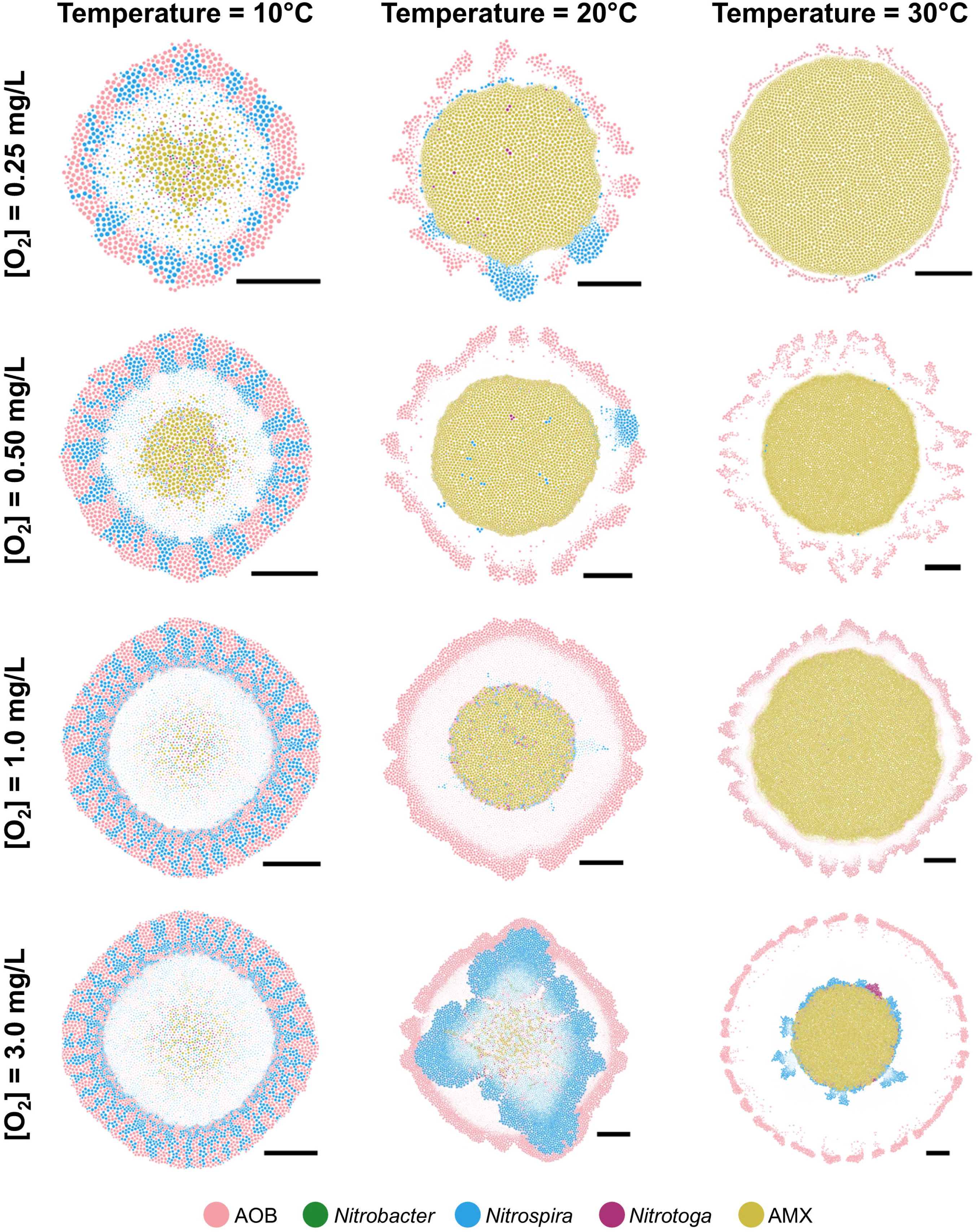
Influence of dissolved oxygen ([O_2_]) and temperature in the community structure. The images show the steady state of the community: 400 days for 0.25 mg-O_2_/L, and 10°C and 30°C; 360 days for 0.25 mg-O_2_/L and 20°C, and 0.5 mg-O_2_/L and 10°C – 30°C; 385 days for 10°C and 1.0 mg-O_2_/L; 120 days for 20°C and 1.0, 3.0 mg-O_2_/L; 85 days for 30°C and 1.0 mg-O_2_/L; 300 days for 10°C and 3.0 mg-O_2_/L; and 315 days for 30°C and 3.0 mg-O_2_/L. Scale bars represents 40 µm. Replicates are included in S3 Fig and S4 Fig. Evolution of granules are shown in S1 Video – S12 Video.

The complete NOB suppression was only achieved at 20°C and 30°C but at different O_2_ concentrations in the bulk liquid for both temperatures. At 30°C, the O_2_ concentration that ensured total NOB suppression was lower than 1.0 mg-O_2_/L (Fig 5 and S7 Fig), with few *Nitrospira* individuals observed at steady state at 0.25 mg-O_2_/L (Fig 5, S3 Fig and S4 Fig). At 20°C, 1.0 mg-O_2_/L was necessary to ensure total NOB suppression (Fig 3C). At 10°C, *Nitrospira* was always present in the aggregate, distributed radially and growing next to AOB in the superficial layer of the granule (*columned stratification* [28], Fig 5 and S8 Fig). This similar distribution of AOB and *Nitrospira* was observed at 20°C and ≤0.5 mg-O_2_/L, but with fewer and more compact clusters of *Nitrospira* (Fig 5). At 30°C and 3.0 mg-O_2_/L NOB (*Nitrospira* and *Nitrotoga*) remained active surrounding the cluster of anammox bacteria in the granule core with an inactive space between them and the AOB layer. The AOB/NOB stratification was observed again at 30°C and 6.0 mg-O_2_/L, with *Nitrobacter* and *Nitrospira* clusters below the AOB layer (S8 Fig). At 10°C and 0.1 mg-O_2_/L, no growth was observed because AOB population failed to persist in such limiting conditions (S7 Fig).

The distinct distribution patterns of AOB and NOB (mainly *Nitrospira* genus) obtained in our simulations (Fig 5, S7 Fig and S8 Fig) are, in part, the result of the dependency of the maximum growth rates with the temperature (S9 Fig). The pattern observed at 10°C (*columned stratification* of AOB and *Nitrospira*; Fig 5, first column) can be explained by the small difference in maximum growth rate between AOB and *Nitrospira* at this temperature (0.0123 h^−1^ and 0.0082 h^−1^, respectively; S9 Fig), which allows for prediction of coexistence (S1 Video, S4 Video, S7 Video and S10 Video). The difference in maximum growth rate between AOB and *Nitrospira* increases with temperature and at 20°C, the higher growth rate of AOB (0.0329 h^−1^ vs. 0.0155 h^−1^, respectively; S9 Fig) explains its dominance in the outer region of the granule, forcing *Nitrospira* to grow into the granule core (S8 Video and S11 Video). In these conditions, nitrite is accumulated in the system, and *Nitrospira* can eventually grow faster than AOB coexisting at the most external layers of the granule (S11 Video). At 30°C and ≤1.0 mg-O_2_/L, the difference in maximum growth rates between AOB and *Nitrospira* is such (0.0825 h^−1^ and 0.0282 h^−1^, respectively; S9 Fig) that *Nitrospira* is always outcompeted by AOB (Fig 5 and S7 Fig). Increasing the concentration of oxygen up to 3.0 mg/L or more at 30°C, NOB individuals (including *Nitrobacter* and *Nitrotoga*) can survive at the granule core and reach AOB (Fig 5 and S8 Fig).

Like at 20°C, the portion of nitrification and nitritation at 10°C and 30°C correlates with the amount of active NOB in the granule (*τ* = 0.7576, *p* = 0.0002 for nitrification vs relative abundance of NOB; *τ* = –0.4545, *p* = 0.0447 for nitritation vs relative abundance of NOB; S10A Fig). Nitrification was the highest contributor of nitrogen conversion at 10°C and 0.25 – 6.0 mg-O_2_/L, and 20°C and ≥3.0 mg-O_2_/L (>94.0%; Fig 4 and S8 Fig). PN/A portion exceeded 60.0% contribution at low oxygen levels (≤0.25 mg-O_2_/L) and ≥20°C (Fig 4), reaching over 90.0% at 30°C and when O_2_ concentration was reduced to 0.1 mg-O_2_/L (S7 Fig). Because NOB compete with anammox bacteria for nitrite, PN/A portion and presence of active NOB are inversely correlated (*τ* = –0.7273, *p* = 0.0005; S10A Fig).

### Physiological diversity of *Nitrospira* and NOB population shifts

To evaluate the influence of *Nitrospira* taxonomy on the community structure in the granules and nitrogen process partitioning, a sensitive analysis of ammonia tolerance and oxygen affinity of *Nitrospira* was performed.

Under the most favourable temperatures for NOB growth (i.e., 10°C; S9 Fig), the complete suppression of NOB (including *Nitrospira*) was achieved at 0.25 mg-O_2_/L when the lowest ammonia tolerance (0.032 mg-NH_3_/L; Fig 6A) and ammonia affinity (5.00×10^−6^ M; Fig 6B) were considered for *Nitrospira*. Increasing the tolerance of ammonia (0.362 mg-NH_3_/L; Fig 6A) or the oxygen affinity (5.00×10^−6^ M; Fig 6B), *Nitrospira* remained in the granule even at low oxygen levels (≤0.5 mg-O_2_/L) and they coexisted with AOB on the outer side of the granule. At 20°C, NOB was successfully suppressed under O_2_ concentrations lower than 3.0 mg/L with both values of ammonia tolerance and oxygen affinity (S11 Fig). These observations indicate that resilient *Nitrospira* was the main impediment to achieve a total NOB suppression at low oxygen concentrations (0.25 mg-O_2_/L at 10°C, and ≤0.5 mg-O_2_/L at 20°C).

**Fig 6.**
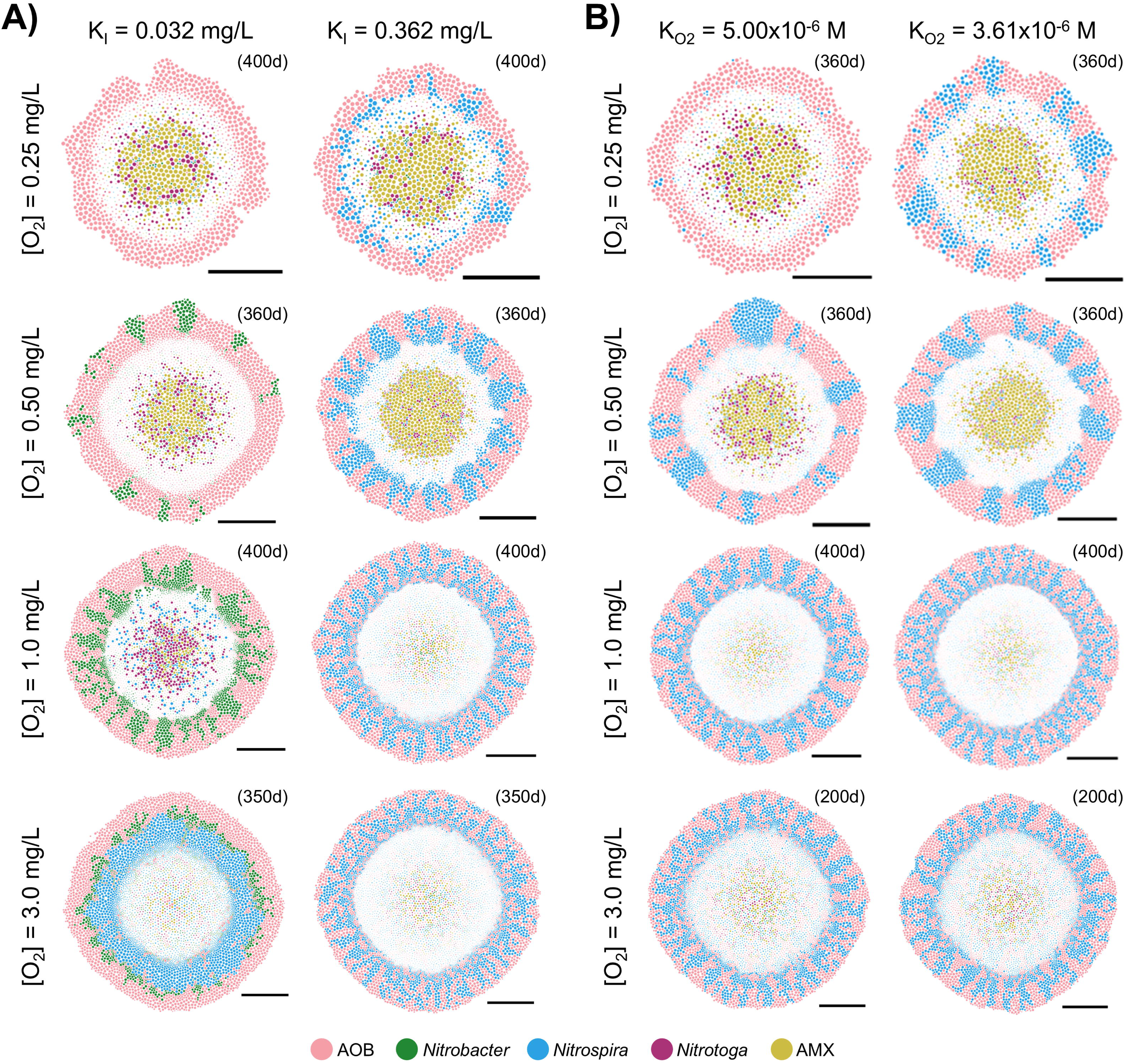
Influence of ammonia tolerance (A) and oxygen affinity (B) of *Nitrospira* on granule community structure at 10°C. The images of granules show steady state of the community. In parenthesis, the time at which this steady state in the liquid bulk was reached. The scale bars represent 40 µm. Replicates are presented in S13 Fig.

When ammonia tolerance for *Nitrospira* was set the lowest (0.032 mg-NH_3_/L), shifts in dominance among the populations of the NOB community were observed (Fig 6A). At 10°C, *Nitrobacter* was the dominant genus at 0.5 mg-O_2_/L, forming a small cluster of individuals on the granule surface, like in Tsuneda *et al.* (2003) [14]; while at 1.0 mg-O_2_/L, *Nitrobacter* was radially distributed next to AOB. *Nitrospira* reappeared again at 3.0 mg-O_2_/L but confined under the AOB layer and coexisting with *Nitrobacter*. At 20°C, the population shift of NOB was observed only at 3.0 mg-O_2_/L (S11 Fig), where *Nitrobacter* dominated (K_I_ = 0.032 mg-NH_3_/L) or coexisted with *Nitrospira* (K_I_ = 0.362 mg-NH_3_/L, K_O2_ = 5.00×10^−6^ mol-O_2_/L and K_O2_ = 3.61×10^−6^ mol-O_2_/L). At 30°C, *Nitrotoga* (instead of *Nitrospira*) surrounds the anammox bacteria under 3.0 mg-O_2_/L (K_I_ = 0.032 mg-NH_3_/L; S12A Fig) and K_O2_ = 5.00×10^−6^ mol-O_2_/L (S12B Fig).

The observed trends on nitrogen process partitioning did not change with the ammonia tolerance and oxygen affinity of *Nitrospira* (Fig 7). A significant nitritation portion was achieved at 10°C when ammonia tolerance and oxygen affinity of *Nitrospira* was reduced to the lowest values (>90% at 0.25 mg-O_2_/L and >40% at 0.5 mg-O_2_/L with K_I_ = 0.032 mg-NH_3_/L and K_O2_ = 5.00×10^−6^ mol-O_2_/L; Fig 7), but the physiology of *Nitrospira* did not influence significatively the nitrogen process partitioning at higher oxygen concentrations (≥1.0 mg/L) and higher temperatures (≥20°C, S4 Table and S5 Table), which evidences the importance of *Nitrospira* resilience on the portion of nitritation at low temperatures. The AMX portion was not significantly improved by a reduced *Nitrospira* tolerance for ammonia (Figs 4 and 7), since the suppression of NOB was achieved at low temperatures where AMX activity was minimum.

**Fig 7.**
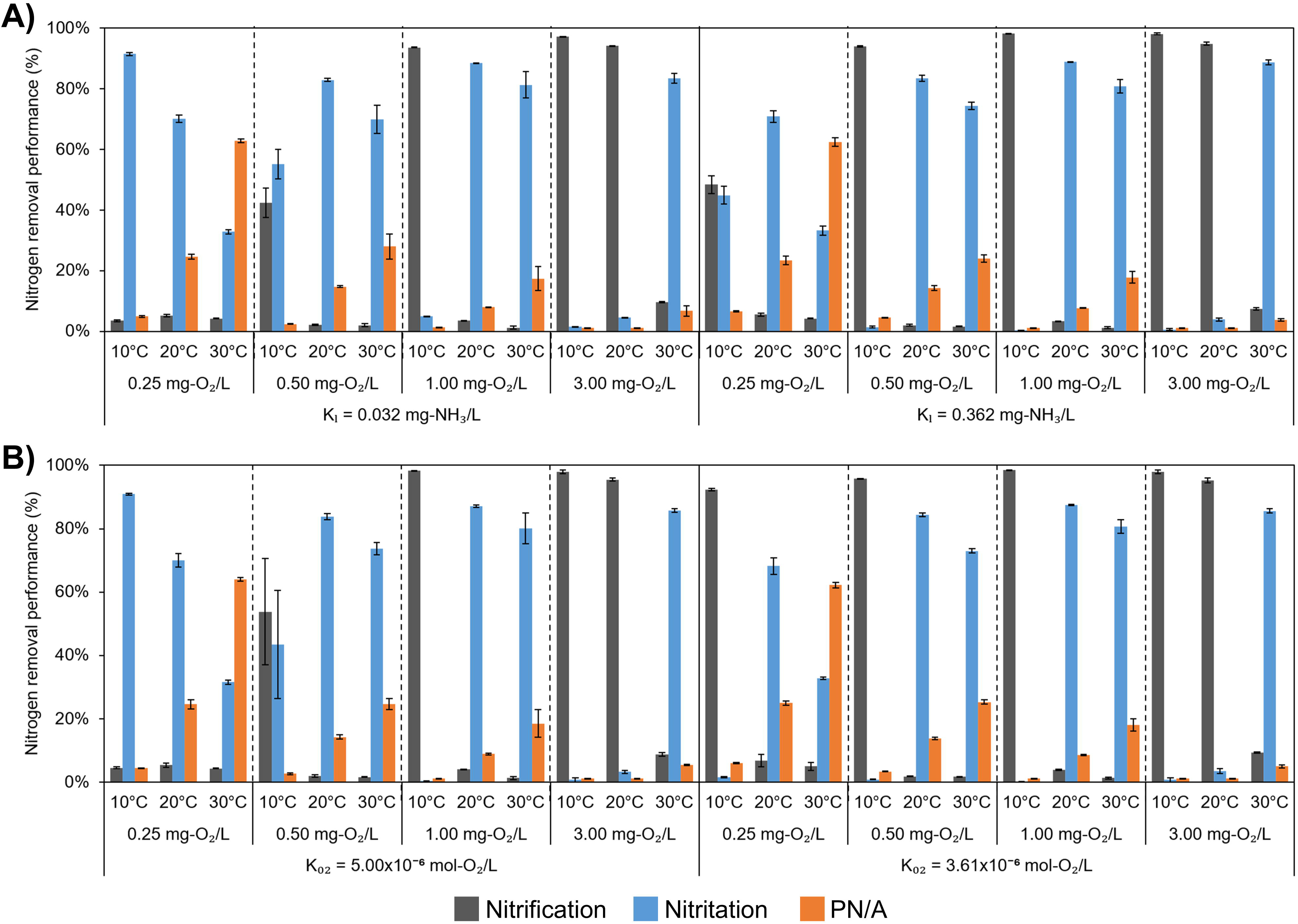
Nitrogen process partitioning at stationary state (average of the last 10 days of operation). **A)** Analysis of the influence of ammonia tolerance of *Nitrospira* (K_I_). Maximum process portion values are included in S6 Table. **B)** Analysis of the influence of oxygen affinity of *Nitrospira* (K_O2_). Maximum process portion values are included in S7 Table. Errors bars show standard deviation of *n* = 3 simulation replicates. The statistical pairwise comparison is shown in S2 Appendix.

The trend in nitrogen conversion with the relative abundance of NOB did not change by reducing the ammonia tolerance or oxygen affinity of *Nitrospira* (*τ* = 0.6897, *p* < 0.0001 for nitrification vs relative abundance of NOB; *τ* = –0.2713, *p* = 0.0065 for nitritation vs relative abundance of NOB; *τ* = –0.7819, *p* < 0.0001 for PN/A vs relative abundance of NOB; S10B Fig). The presence of *Nitrotoga* did not influence the nitritation portion, reaching ≥80% even with 15 wt. % of *Nitrotoga* (triangle symbols, S10B Fig). Although active *Nitrotoga* was present in the granule at certain conditions, it was always found in the core of the granule surrounding anammox bacteria, where the oxygen concentration was low. For this reason, the activity of *Nitrotoga* was not sufficient to reduce the nitrite in the bioreactor. The PN/A portion was influenced by the presence of *Nitrotoga*, not because the nitrite competition (nitrite accumulated in the bioreactor), but rather by the accumulation of oxygen in the granule core, which facilitated the growth of *Nitrotoga* while inhibiting AMX activity. On the other hand, the NOB population shift from *Nitrospira* to *Nitrobacter* at low temperatures (10°C) did not influence the portion of nitrification or nitritation (square and circle symbols, respectively; S10B Fig) because, under these conditions, both populations dominated over AMX activity.

## Conclusions

To maintain an elevated nitrogen removal in one-stage PN/A systems, the presence of an anaerobic core with a persistent AMX activity and the concomitant suppression of NOB populations is required. With the presented model, we have been able to showcase that NOB suppression in one-stage PN/A systems is challenging. Only if oxygen concentration is kept low enough at the bulk liquid (in the simulated conditions lower than 0.5 mg-O_2_/L), ammonia setpoint high (tested conditions fixed at 20.0 mg-NH_3_/L) and temperatures around or higher than 20°C, effective NOB suppression preserving AMX activity is possible. These operational conditions reduce the capacity of NOB to compete for nitrite with anammox bacteria, ensuring the maximization of the ammonia conversion to N_2_ gas. However, if temperatures and/or O_2_ concentrations are too low, the reduced growth rate of nitrifiers will negatively affect the survival of anammox bacteria, reducing the anaerobiosis of the granule core, and maximising total nitrification of the ammonia in the system. Therefore, even under optimal operating conditions, keeping PN/A dominant over nitrification and nitritation within a single granule throughout the year remains challenging.

Contrary to what has been previously hypothesised, we observe that spatial *layered stratification* between AOB and NOB communities positively contributes to the robustness of NOB activity and the overall nitrification activity in the system. In the simulations presented, when this *layered stratification* was observed, anammox bacteria at the core of the granules was in general outcompeted in nitrite consumption by the NOB population.

*Nitrospira*, appears to be the most resilient NOB population in one-stage PN/A systems. The performed sensitivity analysis has shown that if the affinity for O_2_ or ammonia tolerance of *Nitrospira* is reduced, *Nitrobacter* or *Nitrotoga* may colonize the granule, either coexisting or outcompeting *Nitrospira*. However, these population shifts are probably not impacting significantly the observed nitrogen removal, which remained largely similar in all the simulations.

We therefore conclude that NOB physiological diversity, and especially the resilience of *Nitrospira*, is a key design consideration for engineered systems targeting stable PN/A-dominated nitrogen removal (e.g. nitrogen removal in mainstream sewage treatment).

## Methods

### Description of the multiscale model

An Individual-based Model framework was employed to simulate the community assembly of ammonia-oxidizing bacteria (*Nitrosomonas* as the main genus), nitrite-oxidizing bacteria (differentiating *Nitrobacter*, *Nitrospira* and *Nitrotoga* genera) and anammox bacteria (using the physiological properties of *Candidatus* Kuenenia and *Candidatus* Brocadia genera). This community grew in a granular system simulating an ideal continuous stirred tank reactor until the bulk liquid reached steady state. The chemical compounds diffused through the granule, and they were consumed and/or produced by the individuals. Chemical speciation (i.e., the dynamic concentration of various forms of chemical compounds) and the pH dynamics inside the granule are considered in the model (see S3 Appendix – *Chemical speciation and pH*). Chemical speciation is important in this system because ammonia- and nitrite-oxidizing bacteria preferentially consume free ammonia (FA, NH_3_) and free nitrous acid (FNA, HNO_2_) respectively, rather than their ionic forms [29, 30]. The pH of the bulk liquid of the reactor, and throughout the chemical gradient of the granule, is buffered using sodium carbonate (Na_2_CO_3_) and sodium sulphate (Na_2_SO_4_). Individuals would change their growth and decay rates function of the local environment (because of diffusion processes and microbial activity). A detailed description of the mathematical model is included in S3 Appendix.

The model consists of three sub-models: *physical model*, *biological model* and *mechanical model*. The *physical model* comprises a set of diffusion-reaction partial differential equations (2^nd^ Ficks law plus reaction term, Eq 1) coupled with the local spatial position and parametric information of microbial individuals (their mass, stoichiometries and kinetics).

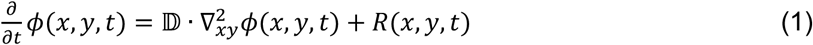

Where *φ*(*x*, *y*, *t*) is the concentration of a soluble component in a specific location of the simulation domain (*x, y*) and in a time step (*t*), D is the effective coefficient of diffusion and *R*(*x*, *y*, *t*) is the biological reaction term. To simplify the resolution of Eq 1, a constant D value is considered for the whole aggregate region [31]. The diffusion-reaction equations were solved through a multigrid method (V-cycle) [32]. For more information, see S3 Appendix – *Solving the diffusion-reaction equation*.

The *biological model* simulates microbial dynamics (growth and decay) following the Monod expression (Eq 2), in which the growth rate of an individual (*μ^n^*) is described function of their maximum growth rate 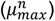, substrate affinities 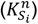, and inhibition constants 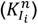, and local concentrations of the limiting substrates (*S_i_*), and inhibitory components (*I_i_*). Inhibition was described as a non-competitive inhibition. Maintenance requirements are considered in Eq 2 as a negative growth rate term (specific maintenance rate, *a^n^*) [33].

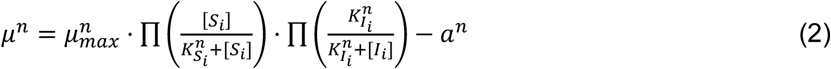

Eq 2 returns a value of *μ^n^* indicating if a microbe *m* is growing (*μ^n^* > 0) or decaying (*μ^n^* < 0). The model assumes that once cells achieve a maximum mass (*M_max_*) they divide. When *μ^n^* < 0 the individual shrinks, reducing its mass. When cells reach a minimum mass (*M_min_*), or a size considered negligible, they become inactive (i.e., they do not grow nor decay) and only can become active again if they increase their mass under more favourable conditions. In this study, the influence of temperature and pH on 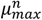 and *a^n^* values were considered. The kinetic parameters and expressions defining the Monod curve for all microbial populations (S2 Table) were chosen based on the guidelines stated in S3 Appendix – *Physiological definition of microbial populations*. The overall metabolic stoichiometry of microbial populations (S8 Table) was determined following the Gibbs Energy Dissipation method and assuming the general biomass formula CH_1.8_O_0.5_H_0.2_ [34].

The bulk concentrations change over time due to the inflow and outflow of reactor, and the metabolic activity in the granule. They are modelled resolving the mass balance of an ideal continuous stirred tank reactor, where the average microbial activity defined at the granule level, is used as a macroscopic reaction term (Eq 3).

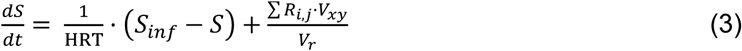

Where HRT is the hydraulic retention time, *S_inf_* is the concentration in the influent, *V_xy_* is the volume of a grid cell, *R_ij_* is the reaction in node (*i,j*), and *V_r_* is the representative volume computed as 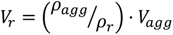 (*ρ_agg_*, density of the aggregate; ρ*_r_*, biomass density of reactor; and 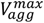, the maximum volume of one granule). In this study, a setpoint of a constant ammonia concentration in the bulk liquid of the reactor is fixed, varying the HRT accordingly.

The *mechanical model* includes the shoving of individuals and the influence of shear forces on the granule morphology (detachment and erosion processes). Due to microbial growth and cell division, individuals might overlap. During shoving, overlapping individuals exert a force, moving them away from each other. An iterative shoving algorithm is executed based on a quadtree-data structure that resolves the overlapping of cells. For more information, see S3 Appendix – *Microbial shoving*. Because the shear forces from bulk liquid are essential for the formation of a microbial granule [35], the model requires an accurate description of detachment and erosion. Although Navier-Stokes equations together with the viscoelastic description of the granule is the most precise methodology, this requires parameters that are not always available and quickly becomes computationally expensive. Instead, the generalized model of detachment proposed by Xavier *et al.* (2005) [36] was used in this study, where the hyperparameter *k_Det_* determines the rate of detachment and sets the size of granule. For more information, see S3 Appendix – *Detachment and Erosion*.

Cells divide in a time scale much slower than the diffusion-reaction process (~1 hour versus ~10^−8^ hours). To solve the system, the model takes advantage of this time scale differentiation to separate the resolution of the diffusion-reaction equation, and the simulation of cell dynamics [37]. A detailed description of the multiscale integration is included in S3 Appendix – *Multiscale integration*.

### Selection of microbial populations

Because the typical temperature and pH of municipal wastewater are 14°C – 22°C and 7.0 – 8.0, respectively [38], this study focuses on mesophilic and neutrophilic microbes. Three main microbial populations were considered: ammonia-oxidizing bacteria (AOB), nitrite-oxidizing bacteria (NOB) and anammox bacteria. In most WWTP, the dominant ammonia and nitrite oxidizers belong to *Nitrosomonas* and *Nitrospira* genera, respectively [39], although *Nitrobacter* and *Nitrotoga* have also been found in some WWTP [40, 41]. Regarding anammox bacteria, *Ca.* Kuenenia, *Ca.* Brocadia, *Candidatus* Anammoxoglobus, *Candidatus* Anammoximicrobium and *Candidatus* Jettenia are the genera most commonly found in WWTP [42].

Based on the most recent literature about nitrifying communities (AOB and NOB) and anammox bacteria in WWTP, the following microbial populations were defined for this study:

- AOB group: a single microbial population was identified, using the physiological properties of *Nitrosomonas* genera.
- NOB group: three microbial populations were identified: *Nitrobacter* (Nb), *Nitrospira* (Ns) and *Nitrotoga* (Ng). Although *Nitrobacter* is not a dominant population in engineered systems, this NOB has been found in some WWTP [12] and in the final states of the partial nitritation operation after the suppression of *Nitrospira* population [11]. For *Nitrospira*, the most resilient individual was considered: highest ammonia tolerance, K_I_ = 2.21 mg-NH_3_/L; highest oxygen affinity, 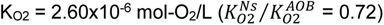.
- Anammox bacteria: a single microbial population was identified, using the physiological properties of *Ca.* Kuenenia and *Ca.* Brocadia.

The physiological definition of microbial populations (i.e., growth kinetic parameters and the influence of temperature and pH on growth rates) is detailed in S3 Appendix – *Physiological definition of microbial populations*.

### Reactor set-up and operational conditions of simulated reactor

All the simulation experiments started with a single granule with the same portion of each microbial population randomly distributed. The simulated reactor operated in continuous mode with a constant temperature, pH and concentration of ammonia and oxygen (at specified setpoint). Different temperatures and oxygen concentrations were applied to explore the role of these parameters on the microbial community assembly and nitrogen process partitioning (nitrification, nitritation and PN/A). During the simulation experiments, the concentration of ammonia and pH in bulk liquid was fixed to 20.0 mg/L and 7.7, respectively. The base case was selected based on:

- Temperature of 20°C, at which the maximum specific growth rate of AOB and *Nitrobacter* intersects (S9 Fig).
- Setpoint of oxygen concentration at 1.0 mg/L, to ensure limiting-oxygen conditions inside the granule and the competition for it between AOB and NOB.
- Setpoint of ammonia concentration at 20.0 mg/L, to ensure the negative impact of free ammonia to NOB community.
- A pH value of 7.7, corresponding to the average optimal pH of nitrifying community (see Appendix S3 – Fig SM8)

Six oxygen concentrations (0.1 mg/L, 0.25 mg/L, 0.5 mg/L, 1.0 mg/L, 3.0 mg/L, 6.0 mg/L) and three temperatures (10°C, 20°C, 30°C) were evaluated. Conditions of all simulation experiments are included in S1 Table. For each condition, the simulations were run in triplicates using different inoculum (randomly generated).

### Evaluation of physiological diversity of *Nitrospira*

Physiological diversity is present within the NOB genera themselves, as shown in S3 Appendix – *Summary of the selection of kinetic parameters* (SM10 Fig). Clearly, it is not reasonable to check all the possible combination of kinetic parameters available in the database from Martinez-Rabert et al. (2022) [20]. Because *Nitrospira* is the most common NOB population in WWTP [39] and due to the relevance of residual ammonia concentration on partial nitritation [11], the simulation experiments (S1 Table) were repeated modifying the ammonia tolerance of *Nitrospira* (set by *K_I_*_,*FA*_) based on the literature values: 0.032 mg-NH_3_/L [43] and 0.362 mg-NH_3_/L [17].

On the other hand, the competition for oxygen between AOB and NOB populations is also determined by the physiological diversity of NOB. While *Nitrobacter* tends to have a lower oxygen affinity than AOB, *Nitrospira* has lower, similar or higher oxygen affinity than AOB (Fig 2B), giving *Nitrospira* the capacity to compete closely against AOB for oxygen. Because there is still debate as to which nitrifying population (AOB or NOB) has a higher oxygen affinity [20, 44–47], a second extra set of simulations was planned to change the oxygen affinity of *Nitrospira*: 5.00×10^−6^ 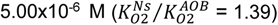 and 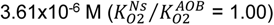.

### Quantification of nitrogen process partitioning

In one-stage PN/A process three main nitrogen conversion processes occur: the aerobic nitrification by AOB and NOB populations, the aerobic nitritation by AOB (supressing NOB activity), and anaerobic ammonia oxidation (anammox process) by anammox bacteria. For nitrification, two partitioning parameters are considered: nitrification portion (ammonia oxidation to nitrate, Eq 4) and nitritation portion (ammonia oxidation to nitrite, Eq 5):

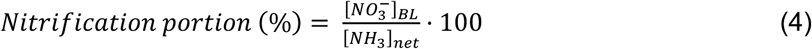

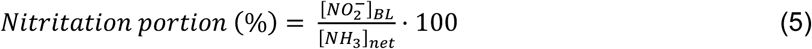

Where 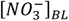 and 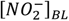 are the concentrations of nitrate and nitrite in the bulk liquid, and [*NH*_3_]*_net_* is the net concentration of ammonia (difference between influent concentration and bulk liquid, [*NH*_3_]*_inf_* − [*NH*_3_]*_BL_*). The rest of ammonia, together with the generated nitrite by AOB, is transformed to N_2_ by anammox bacteria. To calculate the PN/A portion, Eq 6 is employed:

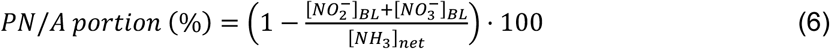

### Statistical analyses

The statistical test to analyse the differences between nitrogen process partitioning (nitrification, nitritation and PN/A) across the different operational conditions, and ammonia tolerance and oxygen affinity of *Nitrospira* was performed using an ANOVA test with Tukey test as post hoc. For comparison tests, the significance was predetermined as *p* < 0.05. To evaluate the correlation between nitrogen process partitioning (nitrification, nitritation and PN/A) and relative abundance of NOB populations (*Nitrobacter*, *Nitrospira*, *Nitrotoga*), Kendall’s rank correlation method (*τ*) was employed [48].

## Acknowledgements

This research did not receive any specific grant from funding agencies in the public, commercial, or not-for-profit sectors.

## Software availability

The source code used to produce the results presented in this study is available in a public GitHub repository at https://github.com/Computational-Platform-IbM/IbM.

## Supporting Information

**S1 Appendix. Supplementary Material 1.** Statistical pairwise comparison of nitrogen process partitioning at stationary state of all simulations (S1 Table) and kinetic parameters shown in S2 Table.

**S2 Appendix. Supplementary Material 2.** Statistical pairwise comparison of nitrogen process partitioning at stationary state of all simulations (S1 Table) plus considering different ammonia tolerance of *Nitrospira* (K_I_; 0.032, 0.362 and 2.21 mg-NH_3_/L) and different oxygen affinity of *Nitrospira* (K_O2_; 2.60×10^−6^, 5.00×10^−6^ and 3.61×10^−6^ M).

**S3 Appendix. Supplementary Methods.** Including explanation of discretization and methodology of diffusion-reaction equation; description of reaction term; details about shoving algorithm; details about detachment and erosion algorithms; the algorithm of the integration process; summary of model boundary, range of application and model assumptions; algorithm to solve chemical speciation and pH; and detailed description of the physiological definition of microbial populations.

**S1 Table. Summary of conditions of simulation experiments.** Symbols: [S]_sp_ – setpoint concentration; [S]_in_ – influent concentration.

**S2 Table. Summary of kinetic parameters of AOB, NOB (*Nitrobacter*, *Nitrospira*, *Nitrotoga*) and AMX.** EST – estimated. For more details, see S3 Appendix – *Physiological definition of microbial populations*.

**S3 Table. Maximum portion values of nitrogen conversion process.** Symbol legend: NF – nitrification, NI – nitritation, PN/A – partial nitritation + anammox. Standard deviations are from three replicates.

**S4 Table. Statistical significance from comparison of nitrogen process partitioning considering different ammonia tolerance of *Nitrospira* (K_I_, mg-NH_3_/L) A)** [O_2_] = 0.25 mg/L; 10°C. **B)** [O_2_] = 0.25 mg/L; 20°C. **C)** [O_2_] = 0.25 mg/L; 30°C. **D)** [O_2_] = 0.5 mg/L; 10°C. **E)** [O_2_] = 0.5 mg/L; 20°C. **F)** [O_2_] = 0.5 mg/L; 30°C. **G)** [O_2_] = 1.0 mg/L; 10°C. **H)** [O_2_] = 1.0 mg/L; 20°C. **I)** [O_2_] = 1.0 mg/L; 30°C. **J)** [O_2_] = 3.0 mg/L; 10°C. **K)** [O_2_] = 3.0 mg/L; 20°C. **L)** [O_2_] = 3.0 mg/L; 30°C. Symbol legend: ns – not significant; * – *p* < 0.05; ** – *p* < 0.01; *** – *p* < 0.001.

**S5 Table. Statistical significance (*p*-value) from comparison of nitrogen process partitioning considering different oxygen affinity of *Nitrospira* (K_I_, mol-O_2_/L) A)** [O_2_] = 0.25 mg/L; 10°C. **B)** [O_2_] = 0.25 mg/L; 20°C. **C)** [O_2_] = 0.25 mg/L; 30°C. **D)** [O_2_] = 0.5 mg/L; 10°C. **E)** [O_2_] = 0.5 mg/L; 20°C. **F)** [O_2_] = 0.5 mg/L; 30°C. **G)** [O_2_] = 1.0 mg/L; 10°C. **H)** [O_2_] = 1.0 mg/L; 20°C. **I)** [O_2_] = 1.0 mg/L; 30°C. **J)** [O_2_] = 3.0 mg/L; 10°C. **K)** [O_2_] = 3.0 mg/L; 20°C. **L)** [O_2_] = 3.0 mg/L; 30°C. Symbol legend: ns – not significant; * – *p* < 0.05; ** – *p* < 0.01; *** – *p* < 0.001.

**S6 Table. Influence of ammonia tolerance of *Nitrospira* (*K_I_*) on maximum portion values of nitrogen conversion processes.** Symbol legend: NF – nitrification, NI – nitritation, PN/A – partial nitritation + anammox. Standard deviation from three replicates.

**S7 Table. Influence of oxygen affinity of *Nitrospira* (*K_O2_*) on maximum portion values of nitrogen conversion processes.** Symbol legend: NF – nitrification, NI – nitritation, PN/A – partial nitritation + anammox. Standard deviation from three replicates

**S8 Table. Metabolic stoichiometries of AOB, NOB (Nitrobacter, Nitrospira and Nitrotoga) and AMX.** Cat and Anab refer to catabolism and anabolism, respectively.

**S9 Table. List of general parameters used in all simulation experiments.** EST – estimated.

**S10 Table. Expressions of growth-temperature influence on growth kinetics.**

**S11 Table. Arrhenius coefficients (*θ*) of *Nitrobacter, Nitrospira* and *Nitrotoga*.**

**S12 Table. Fitting parameters for each model ant the resulted ARD and s_e_.**

**S1 Fig. Evolution of bulk concentrations and granule at 20°C, 0.1 mg-O_2_/L and 20.0 mg-NH_3_/L in bulk liquid.** Arrows indicate the time at which the images of granules were taken (15d, 30d, 60d, 120d, 240d). Errors bars of substrate concentrations show standard deviation of *n* = 3 simulation replicates. Granule images do not follow any scale. Granule replicates included in S7 Fig.

**S2 Fig. Evolution of bulk concentrations and granule at 20°C, 6.0 mg-O_2_/L and 20.0 mg-NH_3_/L in bulk liquid.** Arrows indicate the time at which the images of granules were taken 6d, 15d, 30d, 60d and 120d). Errors bars of substrate concentrations show standard deviation of *n* = 3 simulation replicates. Granule images do not follow any scale. Granule replicates included in S8 Fig.

**S3 Fig. Influence of dissolved oxygen ([O_2_]) and temperature with ammonia concentration of 20 mg/L in bulk liquid (Replicate 2).** Images from steady state in bulk liquid: Images from steady state in bulk liquid: 400 days for 0.25 mg-O_2_/L, and 10°C and 30°C; 360 days for 0.25 mg-O_2_/L and 20°C, and 0.5 mg-O_2_/L and 10°C – 30°C; 385 days for 10°C and 1.0 mg-O_2_/L; 120 days for 20°C and 1.0, 3.0 mg-O_2_/L; 85 days for 30°C and 1.0 mg-O_2_/L; 300 days for 10°C and 3.0 mg-O_2_/L; and 315 days for 30°C and 3.0 mg-O_2_/L. Scale bars represent 40 µm.

**S4 Fig. Influence of dissolved oxygen ([O_2_]) and temperature with ammonia concentration of 20 mg/L in bulk liquid (Replicate 3).** Images from steady state in bulk liquid: Images from steady state in bulk liquid: 400 days for 0.25 mg-O_2_/L, and 10°C and 30°C; 360 days for 0.25 mg-O_2_/L and 20°C, and 0.5 mg-O_2_/L and 10°C – 30°C; 385 days for 10°C and 1.0 mg-O_2_/L; 120 days for 20°C and 1.0, 3.0 mg-O_2_/L; 85 days for 30°C and 1.0 mg-O_2_/L; 300 days for 10°C and 3.0 mg-O_2_/L; and 315 days for 30°C and 3.0 mg-O_2_/L. Scale bars represent 40 µm.

**S5 Fig. Active relative abundances of AOB and NOB (distinguishing *Nitrobacter*, *Nitrospira* and *Nitrotoga*) in simulations at 20°C and oxygen concentrations of 3.0 mg/L (panel A) and 6.0 mg/L (panel B).**

**S6 Fig. Evolution of oxygen profile on the transverse plane of granules at 20°C and bulk liquid concentrations of ammonia and oxygen 20.0 mg/L and 1.0 mg/L, respectively.**

**S7 Fig. Influence of temperature (T) with ammonia and oxygen concentrations in bulk liquid of 20 mg/L and 0.1 mg/L, respectively. A)** Images from steady state in bulk liquid: 360 days for 20°C and 30°C. Scale bars represent 40 µm. **B)** Nitrogen process partitioning at stationary state (average of the last 10d of operation). Symbol legend: NF – nitrification, NI – nitritation, PN/A – partial nitritation + anammox. The process that contributes most to nitrogen conversion is shown in bold. Standard deviations are from three replicates. Maximum portion values of nitrogen conversion processes are included in S3 Table.

**S8 Fig. Influence of temperature (T) with ammonia and oxygen concentrations in bulk liquid of 20 mg/L and 6.0 mg/L, respectively. A)** Images from steady state in bulk liquid: 300 days for 10°C, 120 days for 20°C and 350 days for 30°C. Scale bars represent 40 µm. **B)** Nitrogen process partitioning at stationary state (average of the last 10d of operation). Symbol legend: NF – nitrification, NI – nitritation, PN/A – partial nitritation + anammox. The process that contributes most to nitrogen conversion is shown in bold. Standard deviations are from three replicates. Maximum portion values of nitrogen conversion processes are included in S3 Table.

**S9 Fig. Influence of temperature on the maximum specific growth rate (µ_max_) of AOB, *Nitrobacter*, *Nitrospira*, *Nitrotoga* and AMX at pH 7.7.** Equations to calculate µ_max_ are included in S2 Table.

**S10 Fig. Correlation between nitrogen process partitioning (nitrification, nitritation, PN/A) and active relative abundance of NOB. A)** Simulations considering the most resilient *Nitrospira* (highest ammonia tolerance, K_I_ = 2.21 mg-NH_3_/L; highest oxygen affinity, K_O2_ = 2.60×10^−6^ mol-O_2_/L). **B)** Simulations reducing the ammonia tolerance and oxygen affinity of *Nitrospira*. Smaller plots show a zoom of the region of low relative abundance of NOB (≤15.0 wt. %). Symbol legend: square symbols, *Nitrospira* dominating NOB population; circle symbols, *Nitrobacter* dominating NOB population; triangle symbols, *Nitrotoga* dominating NOB population.

**S11 Fig. Influence of ammonia tolerance (A) and oxygen affinity (B) of *Nitrospira* on granule community assembly at 20°C.** Images of granules taken when steady state in bulk liquid was reached, indicated in parenthesis. Scale bars represent 40 µm. Images of granules assuming the most resilient *Nitrospira* (highest ammonia tolerance and oxygen affinity (K_I_ = 2.21 mg/L, K_O2_ = 2.60×10^−6^ M) are shown in Fig 4. Replicates included in S14 Fig.

**S12 Fig. Influence of ammonia tolerance (A) and oxygen affinity (B) of *Nitrospira* on granule community assembly at 30°C.** Images of granules taken when steady state in bulk liquid was reached, indicated in parenthesis. Scale bars represent 40 µm. Images of granules assuming the most resilient *Nitrospira* (highest ammonia tolerance and oxygen affinity (K_I_ = 2.21 mg/L, K_O2_ = 2.60×10^−6^ M) are shown in Fig 4. Replicates included in S15 Fig.

**S13 Fig. Influence of ammonia tolerance (A) and oxygen affinity (B) of *Nitrospira* on granule community assembly at 10°C (Replicates 2 and 3).** Images of granules taken when steady state in bulk liquid was reached, indicated in parenthesis. Scale bars represent 40 µm.

**S14 Fig. Influence of ammonia tolerance (A) and oxygen affinity (B) of *Nitrospira* on granule community assembly at 20°C (Replicates 2 and 3).** Images of granules taken when steady state in bulk liquid was reached, indicated in parenthesis. Scale bars represent 40 µm.

**S15 Fig. Influence of ammonia tolerance (A) and oxygen affinity (B) of *Nitrospira* on granule community assembly at 30°C (Replicates 2 and 3).** Images of granules taken when steady state in bulk liquid was reached, indicated in parenthesis. Scale bars represent 40 µm.

**S1 Video. Progress of granule at 10°C and 0.25 mg-O_2_/L.** Colour legend – salmon, AOB; green, *Nitrobacter*; light blue, *Nitrospira*; purple, *Nitrotoga*; khaki, AMX. Inactive microbes are shown in a lighter colour.

**S2 Video. Progress of granule at 20°C and 0.25 mg-O_2_/L.** Colour legend – salmon, AOB; green, *Nitrobacter*; light blue, *Nitrospira*; purple, *Nitrotoga*; khaki, AMX. Inactive microbes are shown in a lighter colour.

**S3 Video. Progress of granule at 30°C and 0.25 mg-O_2_/L.** Colour legend – salmon, AOB; green, *Nitrobacter*; light blue, *Nitrospira*; purple, *Nitrotoga*; khaki, AMX. Inactive microbes are shown in a lighter colour.

**S4 Video. Progress of granule at 10°C and 0.5 mg-O_2_/L.** Colour legend – salmon, AOB; green, *Nitrobacter*; light blue, *Nitrospira*; purple, *Nitrotoga*; khaki, AMX. Inactive microbes are shown in a lighter colour.

**S5 Video. Progress of granule at 20°C and 0.5 mg-O_2_/L.** Colour legend – salmon, AOB; green, *Nitrobacter*; light blue, *Nitrospira*; purple, *Nitrotoga*; khaki, AMX. Inactive microbes are shown in a lighter colour.

**S6 Video. Progress of granule at 30°C and 0.5 mg-O_2_/L.** Colour legend – salmon, AOB; green, *Nitrobacter*; light blue, *Nitrospira*; purple, *Nitrotoga*; khaki, AMX. Inactive microbes are shown in a lighter colour.

**S7 Video. Progress of granule at 10°C and 1.0 mg-O_2_/L.** Colour legend – salmon, AOB; green, *Nitrobacter*; light blue, *Nitrospira*; purple, *Nitrotoga*; khaki, AMX. Inactive microbes are shown in a lighter colour.

**S8 Video. Progress of granule at 20°C and 1.0 mg-O_2_/L.** Colour legend – salmon, AOB; green, *Nitrobacter*; light blue, *Nitrospira*; purple, *Nitrotoga*; khaki, AMX. Inactive microbes are shown in a lighter colour.

**S9 Video. Progress of granule at 30°C and 1.0 mg-O_2_/L.** Colour legend – salmon, AOB; green, *Nitrobacter*; light blue, *Nitrospira*; purple, *Nitrotoga*; khaki, AMX. Inactive microbes are shown in a lighter colour.

**S10 Video. Progress of granule at 10°C and 3.0 mg-O_2_/L.** Colour legend – salmon, AOB; green, *Nitrobacter*; light blue, *Nitrospira*; purple, *Nitrotoga*; khaki, AMX. Inactive microbes are shown in a lighter colour.

**S11 Video. Progress of granule at 20°C and 3.0 mg-O_2_/L.** Colour legend – salmon, AOB; green, *Nitrobacter*; light blue, *Nitrospira*; purple, *Nitrotoga*; khaki, AMX. Inactive microbes are shown in a lighter colour.

**S12 Video. Progress of granule at 30°C and 3.0 mg-O_2_/L.** Colour legend – salmon, AOB; green, *Nitrobacter*; light blue, *Nitrospira*; purple, *Nitrotoga*; khaki, AMX. Inactive microbes are shown in a lighter colour.

