## Supplementary material for "Physiological diversity and the assembly of nitrifying communities in aerobic granular sludge": S3 Appendix

### Contents

Simulation domain 2

Solving the diffusion-reaction equation 2

Boundary conditions 4

Domain definition – diffusion and no-diffusion regions 4

Reaction term 6

Microbial shoving 8

Detachment and Erosion 10

Multiscale integration 11

Pseudo-steady state steps 12

Microbial activity steps 12

Summary of model boundary, range of application and model assumptions 14

Chemical speciation and pH 14

Physiological definition of microbial populations 15

Optimal temperature and pH 15

Influence of temperature on growth and specific maintenance rates 16

Influence of pH on growth rate 17

Selection of substrate affinities 18

Selection of inhibition constants 18

Summary of the selection of kinetic parameters 19

References 20

### Simulation domain

The simulation domain is a two-dimensional and micro-scale space. The domain is divided in three different zones: the aggregate, the boundary layer and the bulk liquid (Fig SM1A).

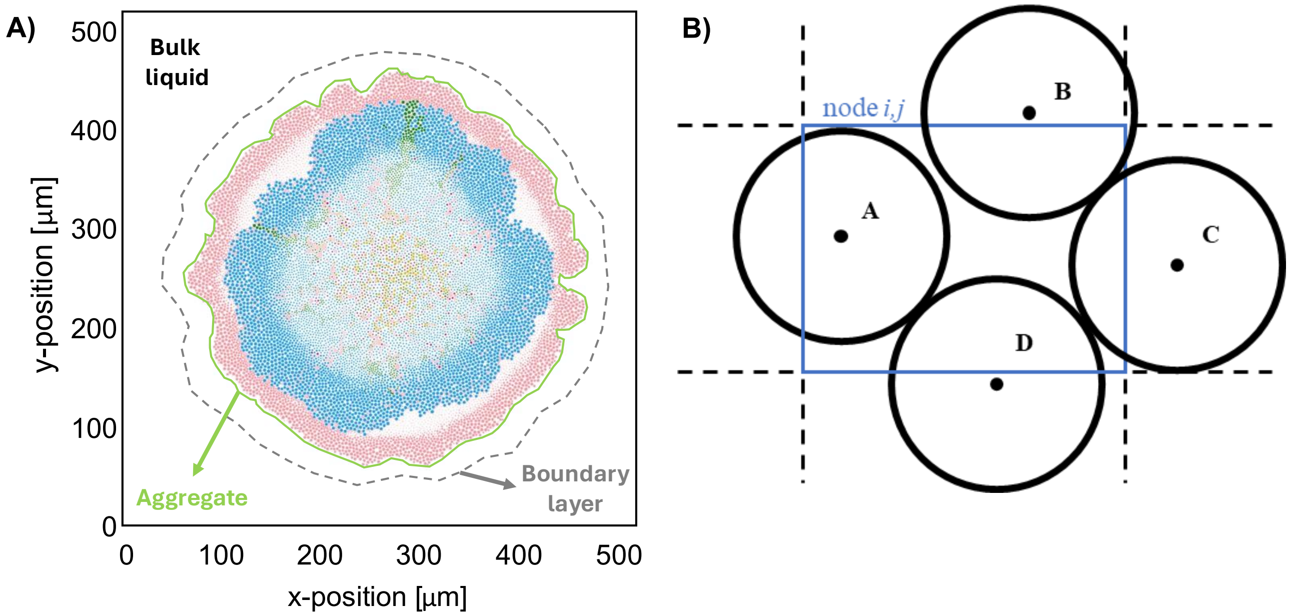


**Fig SM1. A) Representation of simulation domain.** Each circle represents a microbe. Colour legend – salmon, ammonia-oxidizing bacteria; green, *Nitrobacter*; light blue, *Nitrospira*; purple, *Nitrotoga*; khaki, anammox bacteria. Inactive microbes are shown in a lighter colour. **B) Hypothetical position of microbes in a node.** Only "microbe A" belongs to node $(i, j)$ because its centre is inside the node.

When microbes grow and divide, they push each other, thereby increasing the size of the overall aggregate. Diffusion of soluble components occurs throughout the aggregate where they are consumed or produced by the microbes. The boundary layer is the surrounding space of the aggregate defined to simulate the gradient of concentrations between the bulk liquid and the surface of the microbial aggregate. Only diffusion of the soluble components is resolved in this space. At the outside of the boundary layer (considered bulk liquid) the gradient of concentration of all soluble components is considered negligible, assuming a well-mixed homogenous reactor.

The space is discretized in two coordinates (*x* and *y*). This discretization is needed to solve the diffusion-reaction equation in the aggregate and boundary layer, and to set the position of each individual. The position of a microbe, and subsequent assignment of its local conditions, is set by the location of its centre to the grid (Fig SM1B).

### Solving the diffusion-reaction equation

To solve the diffusion-reaction equation (Eq S1), the implicit Crank-Nicolson method, which is unconditionally stable [1], is used to discretize in time the diffusion term. For the reaction term, an explicit forward Euler formula is used (Eq S2). This can be done because the reaction process has a slower time scale than the diffusion [2, 3].

$$\frac{\partial}{\partial t}\phi(x,y,t) = \mathbb{D}\cdot\nabla^2_{xy}\phi(x,y,t) + R(x,y,t) \tag{S1}$$

$$\frac{\phi^{n+1}_{i,j} - \phi^{n}_{i,j}}{h_t} = \mathbb{D}\cdot\frac{1}{2}\left[\nabla^2\phi^{n+1}_{i,j} + \nabla^2\phi^{n}_{i,j}\right] + R(\phi^{n}_{i,j}) \quad n \in 1\ldots N_t \tag{S2}$$

Where $h_t$ refers to the time step, $N_t$ to the total number of time steps and $\phi^n_{i,j}$ to substrate concentration in *node* $i,j$ and time *n*. The Laplacian of Eq S2 ($\nabla^2$) is discretised in a two-dimensional space ($x$, $y$) using the central finite-difference method (Eq S3), where $h$ is the grid size ($h = \Delta x = \Delta y$).

$$\nabla^2\phi^n_{i,j} = \frac{\phi^n_{i-1,j}+\phi^n_{i+1,j}+\phi^n_{i,j-1}+\phi^n_{i,j+1}-4\phi^n_{i,j}}{h^2} \tag{S3}$$

Expression shown in Eq S3 (also known as discrete Laplacian) is given as the following kernel (Eq S4), and the Laplacian approximation is re-written in a matrix from (Eq S5), where the convolution of $[L]$ and $\phi^n_{i,j}$ is denoted using the symbol *.

$$[L] = \begin{pmatrix} 0 & 1 & 0 \\ 1 & -4 & 1 \\ 0 & 1 & 0 \end{pmatrix} \tag{S4}$$

$$\nabla^2\phi^n_{i,j} = \frac{1}{h^2}([L] * [\phi^n]) \tag{S5}$$

Where $[\phi^n]$ is the concentration of soluble compounds ($\phi^n_{i,j}$) can also be defined in matrix form (Eq S6).

$$[\phi^n] = \begin{pmatrix} \phi^n_{1,1} & \phi^n_{1,2} & \cdots & \phi^n_{1,N_y} \\ \phi^n_{2,1} & \phi^n_{2,2} & & \vdots \\ \vdots & & \ddots & \vdots \\ \phi^n_{Nx,1} & \cdots & \cdots & \phi^n_{N_x,N_y} \end{pmatrix} \in M_{N_x \times N_y} \tag{S6}$$

For discrete and 2-dimensional variables (A and B), Eq S7 defines the convolution of A and B (i.e., $A * B$). Convolution satisfies the distributive property, multiplicative identity, and associative property with scalar multiplication. These properties are essential to rearrange the diffusion-reaction equation properly.

$$A[x] * B[x] = \sum_{k=-\infty}^{+\infty} A[x]\cdot B[x-k] \tag{S7}$$

Then the diffusion-reaction equation (Eq S1) can be re-written as a system of matrixes (Eq S8), where $\psi$ is a constant defined for each soluble component (Eq S9) and $[I_k]$ is the so-called *identity kernel* or *do-nothing convolution kernel* (Eq S10).

$$([I_k] - \psi\cdot[L]) * [\phi^{n+1}] = ([I_k] + \psi\cdot[L]) * [\phi^n] + R([\phi^n])\cdot h_t \tag{S8}$$

$$\psi = \frac{\mathbb{D}\cdot h_t}{2\cdot h^2} \tag{S9}$$

$$[I_k] = \begin{pmatrix} 0 & 0 & 0 \\ 0 & 1 & 0 \\ 0 & 0 & 0 \end{pmatrix} \tag{S10}$$

### Boundary conditions

To solve Eq S8 in all the simulation domain, it is necessary to define the boundary conditions of the problem. In this case, a single boundary condition type is defined – the concentrations at outside of the aggregate are defined by the conditions of the bulk liquid. This is implemented by the Dirichlet boundary condition (Eq S11).

$$\phi_{-1,j}^{n} = \phi_{N_x+1,j}^{n} = \phi_{i,-1}^{n} = \phi_{i,N_y+1}^{n} = \gamma \quad \text{(S11)}$$

To implement Dirichlet boundary condition in this system, first the region of the simulation domain is determined. Then, which nodes belong to the diffusion region (comprising boundary layer and granular region) are determined (explained below). Like this, the concentration matrix $[\phi^n]$ is modified accordingly (Eq S12): those nodes in which diffusion region is considered have the corresponding concentration value ($[\phi^n]$), and that ones in which no-diffusion region is considered have the boundary value (i.e., concentration of bulk liquid, $\gamma$).

$$[\phi^n]^{\gamma} = diffR \circ [\phi^n] + \mathrm{inv}(diffR) \cdot \gamma \quad \text{(S12)}$$

Where $diffR$ is a logical matrix with 1 (*true*) in those nodes of diffusion region, and 0 (false) in those nodes of no-diffusion region; $\mathrm{inv}(diffR)$ is the inverse of $diffR$. Then, the diffusion-reaction equation including the boundary conditions imposed over the simulation domain is written in Eq S13. For each soluble component and time iteration, Eq. S24 is solved calculating $\phi^{n+1}$.

$$([I_k] - \psi \cdot [L]) * [\phi^{n+1}] = ([I_k] + \psi \cdot [L]) * [\phi^n]^{\gamma} + R([\phi^n]) \cdot h_t \quad \text{(S13)}$$

### Domain definition – diffusion and no-diffusion regions

The diffusion region, where the diffusion-reaction equation is solved (Eq S13), comprises the aggregate region and boundary layer. The aggregate region corresponds to cells in which microbes are growing. Boundary layer is the region of grid cells in the immediate vicinity of boundary layer.

The distinction of the diffusion region starts with the detection of which nodes are potentially in this region, creating a preliminary diffusion region. For that, the minimum and maximum position of bacteria on aggregate in coordinate *x* and *y*, and boundary layer thickness are considered. The computational cost is reduced by using this preliminary diffusion region instead of the entire simulation domain.

Then, grid cells with bacteria (at least one individual) are determined and included in the diffusion region defining the aggregate region. To recognise which grid cells are belonging to the boundary layer region (and including them in diffusion region), the boundary grid cells with bacteria (i.e., the outermost grid cells of aggregate region) must be sought. An efficient way to find them is through the convolution of aggregate region matrix and *edge detection kernel* (Eq S14).

$$[K]_{ED} = \begin{pmatrix} -{}^{1}/_{8} & -{}^{1}/_{8} & -{}^{1}/_{8} \\ -{}^{1}/_{8} & 1 & -{}^{1}/_{8} \\ -{}^{1}/_{8} & -{}^{1}/_{8} & -{}^{1}/_{8} \end{pmatrix} \quad \text{(S14)}$$

Once boundary grid cells with bacteria are found, it is time to check whether the neighbouring grid cells ($g_{i,j}$) belong to the boundary layer region and, thus, the diffusion region. By dividing the boundary layer thickness by grid size (*h*), the extent of the neighbouring grid cells with possibility to belong to the boundary layer region is obtained (Fig SM3).

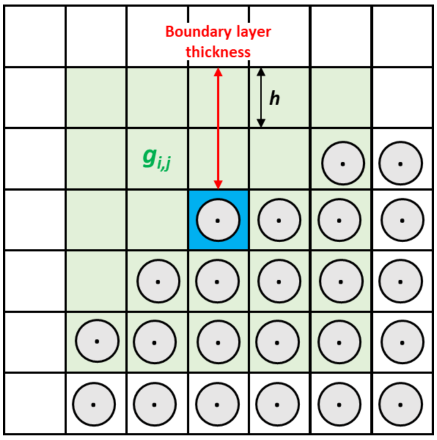


**Fig SM3. Detection of diffusion region.** Neighbouring grid cells (green squares, $g_{i,j}$) of a specific boundary grid cell with bacteria (blue square) with potential to be included in diffusion region.

Finally, the focus region is defined, establishing the region of the simulation domain where diffusion-reaction equation is solved, and the pseudo-steady state is checked. The flowchart of the diffusion region determination is presented in Fig SM4.

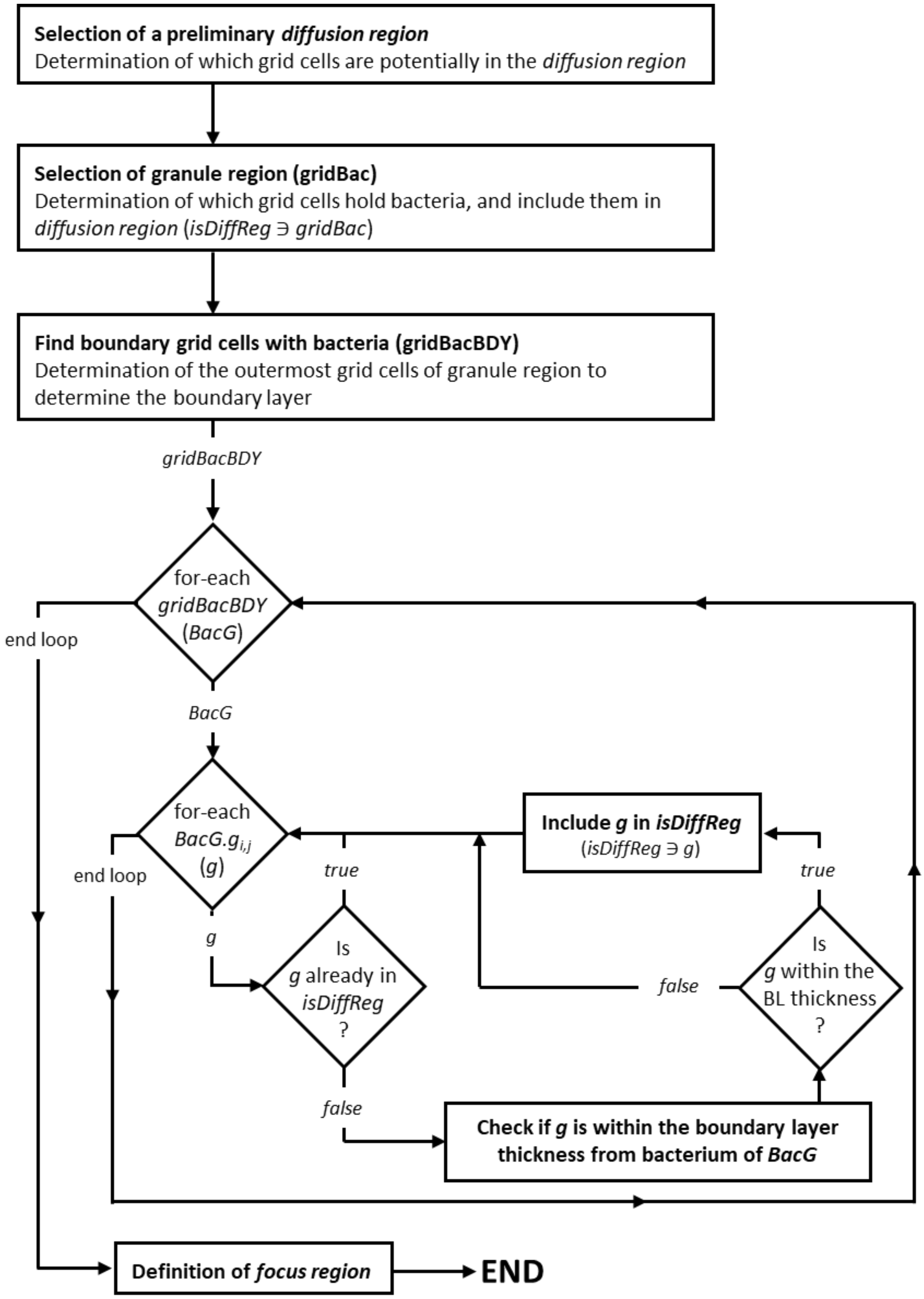


**Fig SM4. Algorithm of diffusion region determination.**

#### Reaction term

The matrix of reaction components $R([\phi^n])$ must be calculated and added to Eq S13. The synthesis and consumption of soluble components are assumed to be the result of microbial activity in the aggregate.

The kinetics of the microbial activity are calculated function of the local conditions of the node where the cell is located. The stoichiometry of microbial species is also necessary to compute the synthesis and consumption of soluble components. Eq S15 – Eq S17 show examples of stoichiometries for anabolism (*Ana*), catabolism (*Cat*) and decay (*Dec*), where $C_s$ and $N_s$ refer to carbon and nitrogen sources, $eD$ and $eA$ to electron donor and acceptor and *X* to biomass considering an average formula of $C_1H_{1.8}O_{0.5}N_{0.2}$ [4, 5].

$$Ana \ = \ 1 \cdot C_s + 0.2 \cdot N_s \ + \cdots \rightarrow \cdots + \ 1 \cdot \ X \tag{S15}$$

$$Cat \ = \ \alpha \cdot \ eD \ + \ \beta \cdot \ eA \ \rightarrow \cdots \tag{S16}$$

$$Dec \ = \ 1 \cdot \ X + \cdots \rightarrow \ 1 \cdot C_s + 0.2 \cdot N_s \tag{S17}$$

The stoichiometry of the overall metabolism (*Met*) is calculated function of the catabolic and anabolic stoichiometries, and the growth yield ($Y_{XS}$, Eq S18).

$$Met \ = \frac{1}{Y_{XS}} \cdot Cat + Ana \tag{S18}$$

With the metabolic stoichiometry and the kinetic parameters ($\mu_{max}$, $K_S$, $K_I$ and *a*) for each of the microbial species present in the reactor, the growth rate is calculated ($\mu$, Eq 2; main text). Then, through the stoichiometry and the bacterial mass, the growth/decay of the cell, substrate uptakes and product generations are calculated (Eq S19 and Eq S20, respectively). Eq S19 is the derivative of the mass of a microbe which is integrated in time using a forward Euler scheme; and Eq S20 computes the reaction term for a specific soluble component *s* due to the activity of the cell *m*.

$$\frac{dX_m}{dt} \ = \ \mu_m^n \cdot \ X_m^n \tag{S19}$$

$$R_m^n \ = \ \frac{\mu_m^n \cdot \delta_{s,m}}{V_{xy}} \tag{S20}$$

Where $X_m$ refers to the mass in moles of the cell *m*, $\delta_{s,m}$ to the stoichiometric coefficient of the substrate S for the cell *m* and $V_{xy}$ to the volume of one node of the simulation domain. The calculation of the reaction term for the soluble components is function of the position in the simulation domain (one reaction term per node for each soluble component). Eq S21 is evaluated considering all the microorganisms that are contributing to the reaction term of a specific soluble component in the *node* $i, j$.

$$R_{i,j}^n = \sum_{m\,=\,1}^{m\,=\,M_{i,j}} R_m^n \tag{S21}$$

Where $M_{i,j}$ refers to all microorganism that are in the *node* $i, j$. The reaction terms $R_{i,j}^n$ calculated by Eq S21 for all nodes of the simulation domain ($R([\phi^n])$), are then used in Eq S13 to calculate the concentration of each of the soluble components in the simulation domain.

### Microbial shoving

To compute the shoving of the cells in the aggregate after cell growth and division, first the overlap between microorganisms is checked by Eq S22 and Eq S23.

$$|\vec{v}| = \sqrt{(x_m - x_{m+1})^2 + (y_m - y_{m+1})^2} \quad \text{(S22)}$$

$$overlap = kDist \cdot (r_m + r_{m+1}) \; - |\vec{v}| \quad \text{(S23)}$$

Where $|\vec{v}|$ is the norm of the vector that links the centres of both cells *m* and *m+1* and $kDist$ is just a multiplier that allows adjustment of the minimal spacing between bacteria. In this case, the *quadtree algorithm* is applied to detect the *overlapping* between bacteria. Subsequently, the shoving of overlapping microbes is computed [6]. If the *overlap* value is bigger than the distance allowed by the user, then microorganisms are pushing each other function their mass and distance (Eq S24 – Eq S29).

$$\vec{p} = \frac{kDist \cdot (r_m + r_{m+1}) \; - |\vec{v}|}{|\vec{v}|} \quad \text{(S24)}$$

$$a_m \; = 1 - \frac{X_m}{X_m + X_{m+1}} \; ; \; a_{m+1} = 1 - \frac{X_{m+1}}{X_m + X_{m+1}} \quad \text{(S25)}$$

$$x_{new,m} = \; x_{old} - (x_{m+1} - x_m) \cdot a_m \; \cdot \vec{p} \quad \text{(S26)}$$

$$y_{new,m} = \; y_{old} - (y_{m+1} - y_m) \cdot a_m \; \cdot \vec{p} \quad \text{(S27)}$$

$$x_{new,m+1} = \; x_{old} + (x_{m+1} - x_m) \cdot a_{m+1} \cdot \vec{p} \quad \text{(S28)}$$

$$y_{new,m+1} = \; y_{old} + (y_{m+1} - y_m) \cdot a_{m+1} \cdot \vec{p} \quad \text{(S29)}$$

Overlap assessment and shoving estimation (if needed) are only performed over the neighbour individuals. Because bacteria are not motile and therefore will not move very far from the original position, it is assumed that chosen neighbouring bacteria stay constant. An overview of the shoving algorithm is presented in Fig SM5.

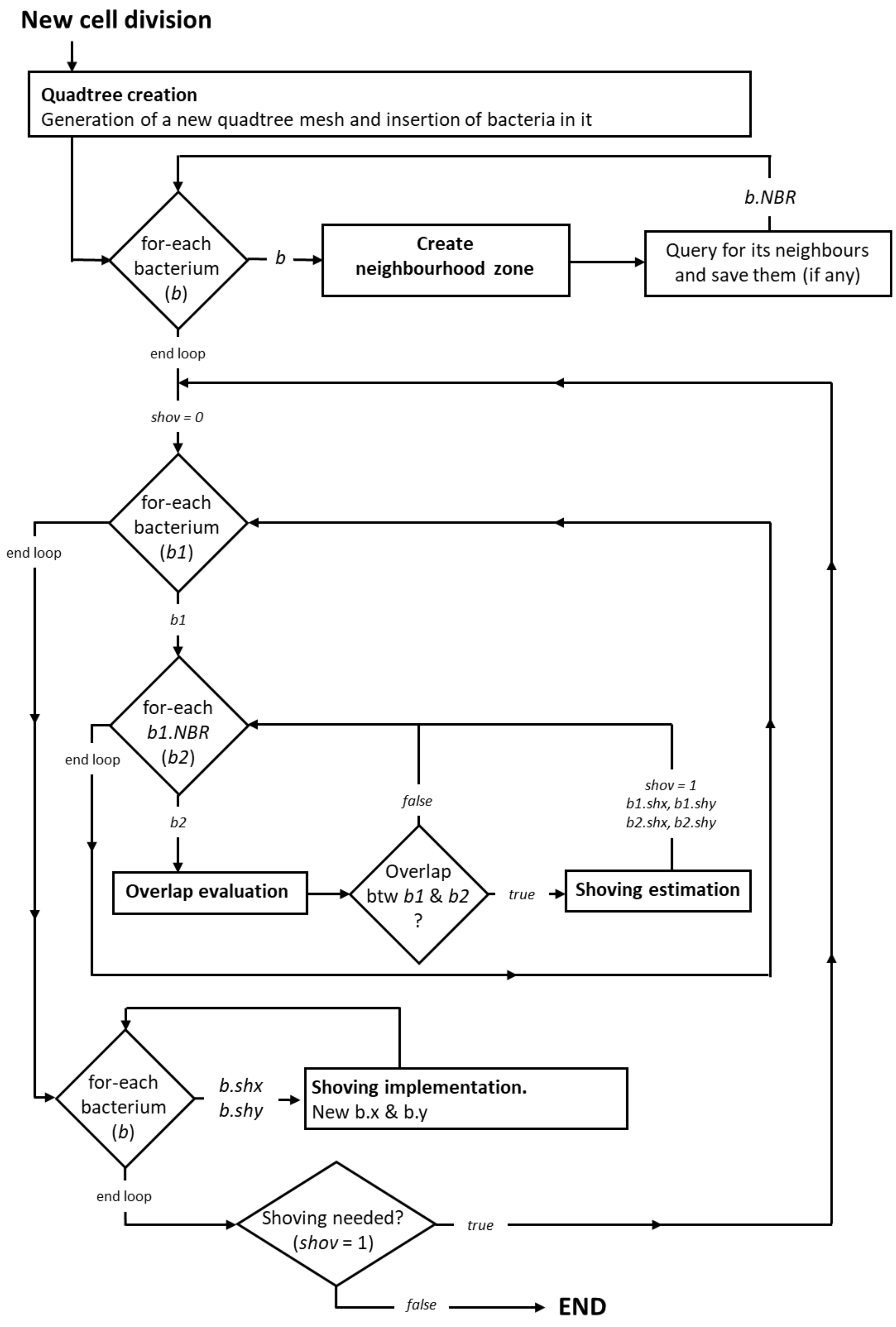


**Fig SM5. Scheme of shoving algorithm.** Legend: $b.shx$ and $b.shy$ – displacement of bacterium *b* due to shoving of the others; $b.NBR$ – neighbours of bacterium *b*.

### Detachment and Erosion

Detachment and erosion descriptions developed by Xavier et al. are based on a global detachment rate ($k_{Det}$) [7]. It is assumed that detachment is a moving of the edge of the granule (i.e., the 'front', $\Gamma$). The edge of the granule is established by the local detachment speed ($F_{det}$), defined as the perpendicular speed to the front of the microbe at which it is removed. In this study, this speed is proportional with the squared distance from the centre of the aggregate (Eq S30).

$$F_{det}(x,y) = d_c(x,y)^2 \cdot k_{Det} \tag{S30}$$

Where $d_c$ is the distance from the centre and $k_{Det}$ is the global detachment rate. As biomass can never be accumulated due to the detachment, this speed is always positive. Considering this, detachment can be described with the Eq S31.

$$F_{det}|\nabla T(x,y)| = 1, \qquad T = 0 \; on \, \Gamma \tag{S31}$$

Where $T(x,y)$ is the time that it takes before the front $\Gamma$ reaches the position $(x,y)$. Although shear forces are applied continuously, detachment and erosion can be discretized in time by using the local time of detachment across the granule at each detachment calculation. The local time of detachment are calculated using the fast-marching method [8, 9]. In order to calculate the time of detachment at each grid cell, Eq S31 is first discretized and approximated using an explicit function, giving Eq S32.

$$\left(\Delta_{i,j}^{x}T\right)^2 + \left(\Delta_{i,j}^{y}T\right)^2 = -\frac{1}{F_{i,j}^2} \tag{S32}$$

Where $\Delta_{i,j}^{x}T$ is given by $\left[T_{i,j} - \min\left(T_{i+1,j} - T_{i-1,j}\right)\right]$ and $\Delta_{i,j}^{y}T$ by $\left[T_{i,j} - \min\left(T_{i,j+1} - T_{i,j-1}\right)\right]$. In the fast-marching algorithm, the current position of the front is kept tracking constantly. The front is then continuously advanced by moving towards the grid cell with the next lowest time of detachment. After moving the front, the time it takes to reach the grid cells next to the front is determined by solving Eq S32. For accuracy of the fast-marching method, it is important to only use the time of detachment values of grid cells that are located on the front in Eq S32 [9]. If in one direction there is no cells located on the front, this term $\left(\Delta_{i,j}^{x}T \text{ or } \Delta_{i,j}^{y}T\right)$ is omitted. Once the front has traversed the entire aggregate, the time of detachment in each grid cells is determined (Fig SM6). The time of detachment values are compared to the timestep of detachment calculation ($\Delta t_{det}$), which yields the ratio $r$ (Eq S33)

$$r = {\Delta t_{det}}/{T_{i,j}} \tag{S33}$$

Individuals whose centre is located in grid cells with a ratio greater than 1, meaning that the time of detachment is smaller than the timestep taken, are removed. All other individuals have a fraction $r$ eroded from their biomass.

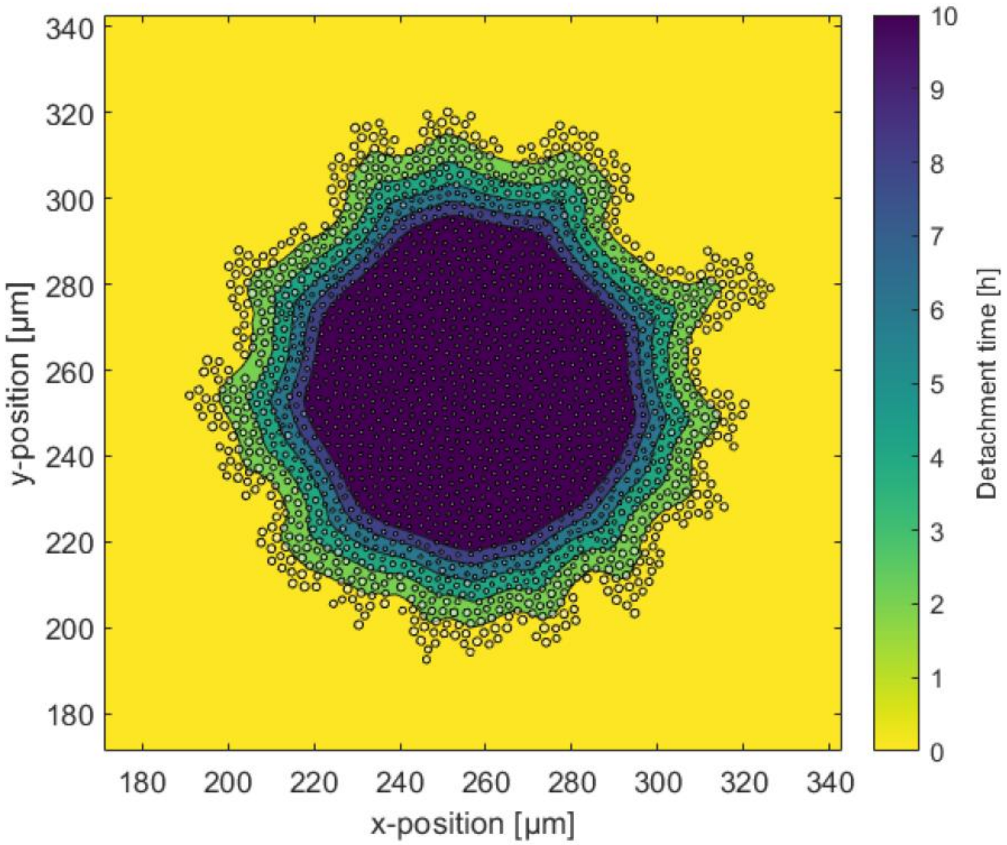


**Fig SM6. Representation of detachment based on Xavier et al. model [7].** Determined time of detachment calculated with an artificially high detachment rate of $kDet = 1000\ \mathrm{m^{-1}h^{-1}}$, where the detachment speed is proportional to the squared distance from the centre of the granule. Circles represent individuals of the granule. Isolines show the detachment front after $2n$ hours.

### Multiscale integration

The integration of domains with different spatial and temporal scales (multiscale system) is performed as follows. First, the diffusion-reaction equation is integrated (with a time step *dt*) until it reaches a pseudo-steady state, allowing a deviation of up to 1% from the real steady state (Eq S34). To check whether pseudo-steady state is reached or not, only diffusion region is considered, as the Dirichlet boundary condition and the bulk liquid concentrations are constants in this specific time span, changing only when the microbial community change significantly (due to a cell division, cell inactivation or a substantial variation in microbial mass).

$$RES = \mathbb{D} \cdot [L] * [\phi]^{\gamma} + R([\phi]) \tag{S34}$$

When the pseudo-steady state is reached, the biomass growth is integrated in a bigger time step ($dt_{bac}$). Once the mass of all individuals is updated, the mass balances of the overall reactor are integrated with the time step $dt_{bac}$ considering the average microbial activity of the aggregate. After that, the Dirichlet boundary condition and the reaction term are updated. At the end of this bigger step, the diffusion-reaction equation needs to be integrated again to reach a new pseudo-steady state.

In modelling, there is always a trade-off between accuracy and performance. To speed up the model, maintaining accuracy, an adaptative time step is used both for reaching pseudo-steady state and for the main iterations. The main concept is to integrate with larger steps when changes are small and integrate with shorter steps if there are large changes.

**Pseudo-steady state steps**

The initial step for reaching pseudo-steady state (*dt*) is based on the Von Neumann stability coefficient [10]. Clamped between a minimum and maximum value (0.01 and 0.5, respectively), *dt* is reduced under three conditions:

- At any point if there is an increase in $\max|RES_{i,j}|$ values detected.
- The solution is not converging any more. Empirically this means that the solution is stuck in an oscillation, where it breaks when *dt* is reduced.
- When any solute concentration in bulk liquid or aggregate region becomes negative.

The *dt* is only increased if multiple pseudo-steady states have been reached with more diffusion iteration than a threshold, while there has not been any change to the *dt*.

**Microbial activity steps**

The time step for microbial activity ($dt_{bac}$) is dependent on the bulk concentrations. Also clamped between a minimum and maximum (0.05 h and 0.5 h, respectively), $dt_{bac}$ is decreased to ensure a maximum relative difference for any bulk concentration of 2%. If multiple iterations have been performed without decreasing $dt_{bac}$, this is incrementally increased over multiple iterations until the relative difference in any bulk concentration or maximum threshold is reached.

An overall scheme of model integration, and summary of parameters are presented in Fig SM7 and S9 Table, respectively.

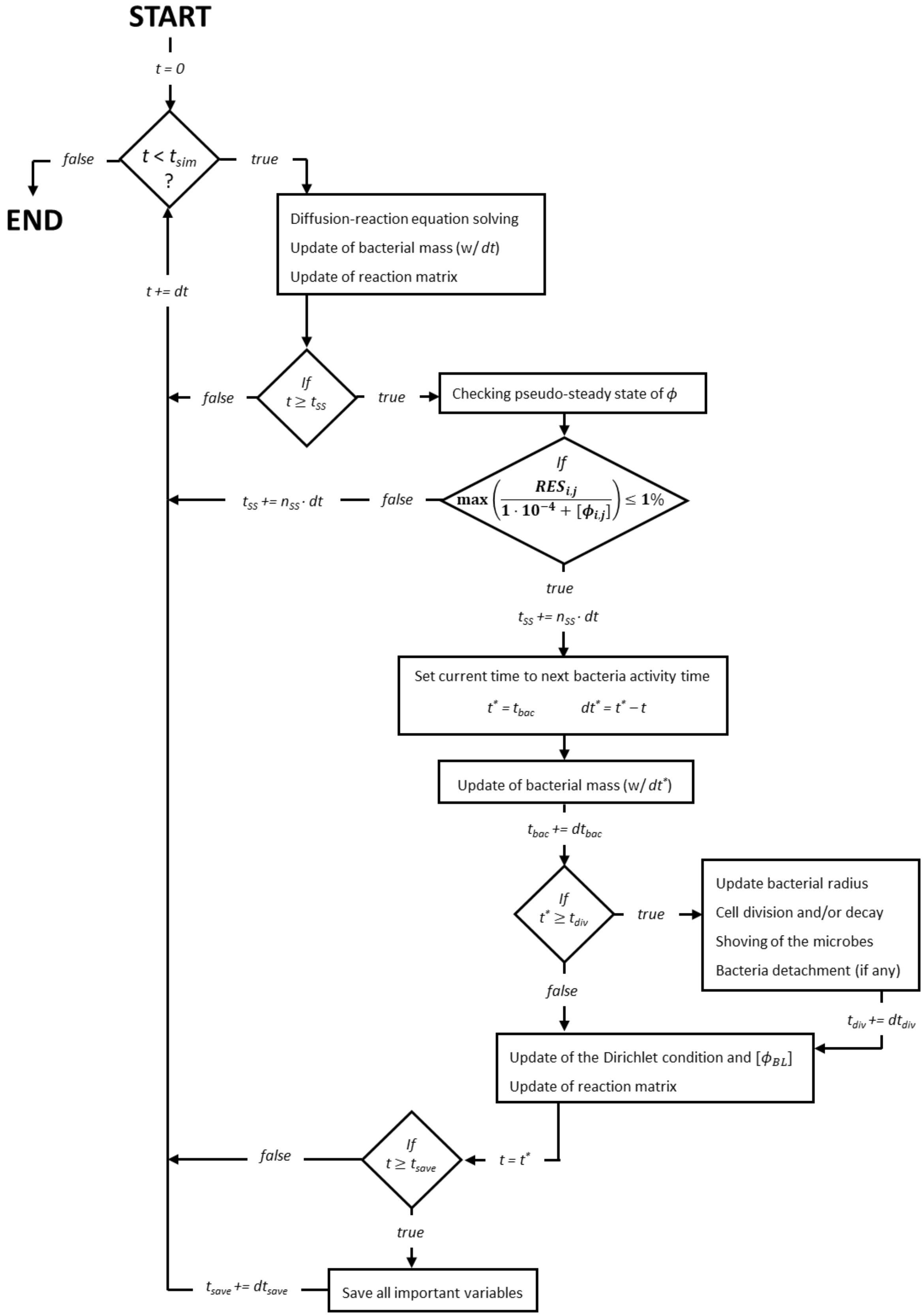


**Fig SM7. Algorithm scheme of the integration process.**

### Summary of model boundary, range of application and model assumptions

· **Model boundary and range of application.** The aim of this study is to understand the implications of the diversity of nitrite-oxidizing bacteria on the nitrogen process partitioning and conversion stability between nitrification ($NH_3 \rightarrow NO_3^-$), nitritation ($NH_3 \rightarrow NO_2^-$) and one-stage partial nitritation/anammox process (PN/AMX, $NH_3 + NO_2^- \rightarrow N_2$).

This model is capable to simulate any other microbial community in self-aggregated systems: dens granules (this study), or suspended flocs [11]. For more information, see the public GitHub repository at Computational-Platform-IbM/IbM (github.com).

· **Model assumptions.** Assumptions are the set of explicitly stated (or implicit premised) conventions and choices that fulfil the holes in our interpretation of reality. These establish the limits of our model and simplify the problem. Main assumptions applied in this model are listed below:

- Cell division and inactivation occur at a certain microbe's size, independently of the state of other cells. Cell growth and decay is established by the actual specific growth rate ($\mu$): $\mu > 0$ – microbial growth; $\mu < 0$ – microbial decay.
- No lysis cellular is considered. When a cell reaches a minimum size, this becomes inactive.
- Only diffusive transport of substrate and products throughout boundary layer and aggregate regions is considered. Convection transport of substances is not considered here.
- The growth of individuals is only affected by local conditions, which in turn are influenced by the activity of surrounding microbes.
- The system was simulated as an ideal continuous stirred tank reactor.
- Simulated aggregates are considered a representation of the activity of the whole reactor.
- Bulk liquid dynamics is only influenced by the microbial activity in aggregates except oxygen, which is assumed to be constant in bulk liquid. Planktonic cells are not considered in this study.
- Static metabolism for all microbial populations is assumed.
- Active movement (cell motility) is not considered. Microbes can move passively due to shoving forces exerted by neighbouring cells as they grow and divide.

### Chemical speciation and pH

Chemical (or ion) speciation refers to the changing concentration of varying forms of chemical species as the pH of the solution changes [12]. The concentration of all chemical species is calculated in a generalized format, considering the chain of equilibrium reactions shown in Eq S35.

$$H_2O + A \xleftrightarrow{K_d} B \xleftrightarrow{K_{a1}} H^+ + B^- \xleftrightarrow{K_{a2}} 2 \cdot H^+ + B^{2-} \quad \text{(S35)}$$

Where the equilibrium constants and total compound concentration ($C_t$) are defined as:

$$K_d = \frac{[A]}{[B]}, \qquad K_{a1} = \frac{[H^+][B^-]}{[B]}, \qquad K_{a2} = \frac{[H^+][B^{2-}]}{[B^-]} \quad \text{(S36)}$$

$$C_T = [A] + [B] + [B^-] + [B^{2-}] \quad \text{(S37)}$$

By combining and reorganizing Eq S36 and Eq S37, all chemical species concentrations can be expressed as a function of $[H^+]$ ($[H^+] = 10^{-pH}$), the total concentration and equilibrium constants (Eq S38), where $\theta = (1 + K_d) \cdot [H^+]^2 + K_{a1} \cdot [H^+] + K_{a1} \cdot K_{a2}$. The acid equilibrium constants ($K_i$) are estimated using the Nernst relation at equilibrium state (Eq S39).

$$[A] = K_d \cdot \frac{C_T\,[H^+]^2}{\theta},\ [B] = \frac{C_T\,[H^+]^2}{\theta},\ [B^-] = K_{a1} \cdot \frac{C_T\,[H^+]}{\theta},\ [B^{2-}] = K_{a1} \cdot K_{a2} \cdot \frac{C_T}{\theta} \quad \text{(S38)}$$

$$K_{ai} = \exp\left(\frac{\Delta G_r^0}{-R \cdot T}\right) \quad \text{(S39)}$$

Where $\Delta G_r^0$ is the standard Gibbs free energy of chemical speciation [kJ/mol] ($\Delta G_r^0 = \sum v_i \cdot \Delta G_f^0$, where $v_i$ is the stoichiometry of reaction) and $R$ is the universal gas constant [0.008314 kJ/mol/K]. The $\Delta G_f^0$ values were collected from literature [13-16]. When a compound does not have all equilibrium steps shown in Eq S35, this generalized formalization of chemical speciation is reduced into the correct and proper form when the non-required equilibrium constants are set to 0.

Microbial activity is also affected by pH. For this reason, acid-based equilibrium is included in the model. The change of pH in the reactor and throughout granule can be estimated by the charge balance. Due to equilibrium and acid-base reactions have fast kinetics (in comparison to diffusion-reaction process and microbial dynamics), these can be considered instantaneous [17]. The dynamic pH (expressed as $[H^+]$) can be determined by solving Eq S40.

$$[H^+] + \sum z_k \cdot [C_k] = 0 \quad \text{(S40)}$$

Where $z_k$ is the charge of chemical species $k$ and $[C_k]$ is the concentration of chemical species $k$. To find the root of Eq S40 (i.e., the value of $[H^+]$), the Newton-Raphson method is used [18]. Then, the value of pH is calculated as $pH = -\log([H^+])$.

### Physiological definition of microbial populations

This section details the guidelines followed for the selection of kinetic parameters and expressions that define microbial growth and activity. The necessary kinetic parameters for Monod expression are the maximum specific growth rate ($\mu_{max}$), substrate affinities ($K_S$), inhibition constants ($K_I$) and specific maintenance rate ($a$). For the estimation of $\mu_{max}$ and $a$ values, the influence of temperature (all microbial populations) and pH (all microbial populations except anammox bacteria) was considered. Additionally, analyses of optimal temperature ($T_{opt}$) and optimal pH ($pH_{opt}$) have been included. Kinetic database of AOB and NOB comes from a previous study [19]. For AMX, the following studies were considered: [20-31].

#### Optimal temperature and pH

Selected AOB, *Nitrobacter* (NOB), *Nitrospira* (NOB) and AMX have similar optimum temperature ranges. Only *Nitrotoga* has a slightly lower optimal temperature range than the other microbial populations, although still within the range of temperatures typical in municipal wastewater (Fig SM8A). On the other hand, all microbial populations (AOB, NOB and AMX) have similar pH optimum range (Fig SM8B).

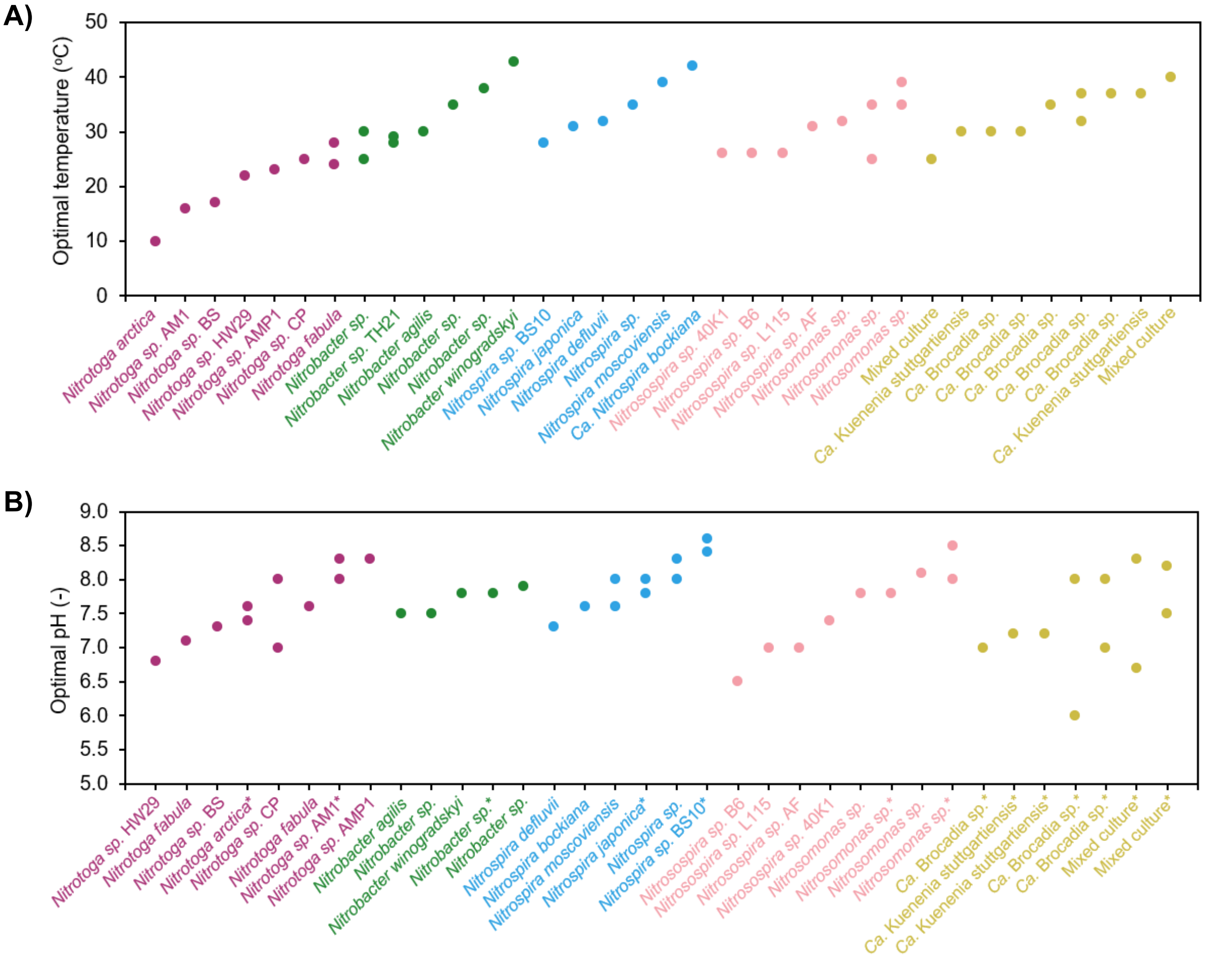


**Fig SM8. A) Optimum temperature for known NOB (*Nitrobacter*, *Nitrospira*, *Nitrotoga*), AOB and AMX. B) Optimum pH for known NOB (*Nitrobacter*, *Nitrospira*, *Nitrotoga*), AOB and AMX.** Asterisks on names indicate those pH values in which refer to pH cultivation rather than pH optimum. Colour labels – purple: *Nitrotoga*; green: *Nitrobacter*; light blue: *Nitrospira*; salmon: AOB; khaki: AMX. References of optimal temperature: [23, 26, 27, 30-52]. References of optimal pH: [23, 26, 27, 29, 30, 32-34, 36-38, 40-43, 45-54].

#### Influence of temperature on growth and specific maintenance rates

Considering operational temperatures of simulations (10°C, 20°C and 30°C) and optimum temperatures of AOB, NOB (*Nitrobacter, Nitrotoga* and *Nitrospira*) and AMX (Fig SM8A), Arrhenius equation (S10 Table) was selected to simulate the influence of temperature on maximum specific growth rate ($\mu_{max}$) and specific maintenance rate ($a$) of AOB, *Nitrobacter*, and AMX ($T_{opt} \geq 30^{o}C$). The Arrhenius equation for AOB and *Nitrobacter* (NOB) comes from Jubany et al. (2008) [53]. In this project, no *Nitrospira* was detected, only *Nitrobacter* [55]. Because the temperature influence on growth rate of *Nitrobacter* and *Nitrospira* are not significantly different (S11 Table, *p* > 0.9), the same activation energy ($E_a$) value was assumed for *Nitrobacter* and *Nitrospira*. Based on the $\mu_{max}^{20^{o}C}$ values gathered from literature (0.0138 ± 0.0091 $h^{-1}$ for Nitrobacter [31, 33-38]; 0.0087 ± 0.0027 $h^{-1}$ for *Nitrospira* [35, 39, 40]), the Arrhenius frequency factor ($A$)

value of *Nitrospira* was the 63% of the *Nitrobacter* value. For AMX, the Arrhenius equation comes from Puyol et al. (2014) study [56].

For *Nitrotoga*, Hinshelwood model, modified Ratkowsky and Yan & Hunt model (S10 Table) were tested ($T_{opt} < 30^{o}\mathrm{C}$). Experimental data was obtained from *Candidatus* Nitrotoga fabula, the first *Nitrotoga* specie isolated in activated sludge [57]. The fitting results with experimental data is shown in Fig SM9, and equation parameters together with the resulted average relative deviations (ARD) and standard error of estimate ($s_e$) are gathered in S12 Table. Comparing ARD and $s_e$ from the tested models, the Yan & Hunt model was chosen for *Nitrotoga*.

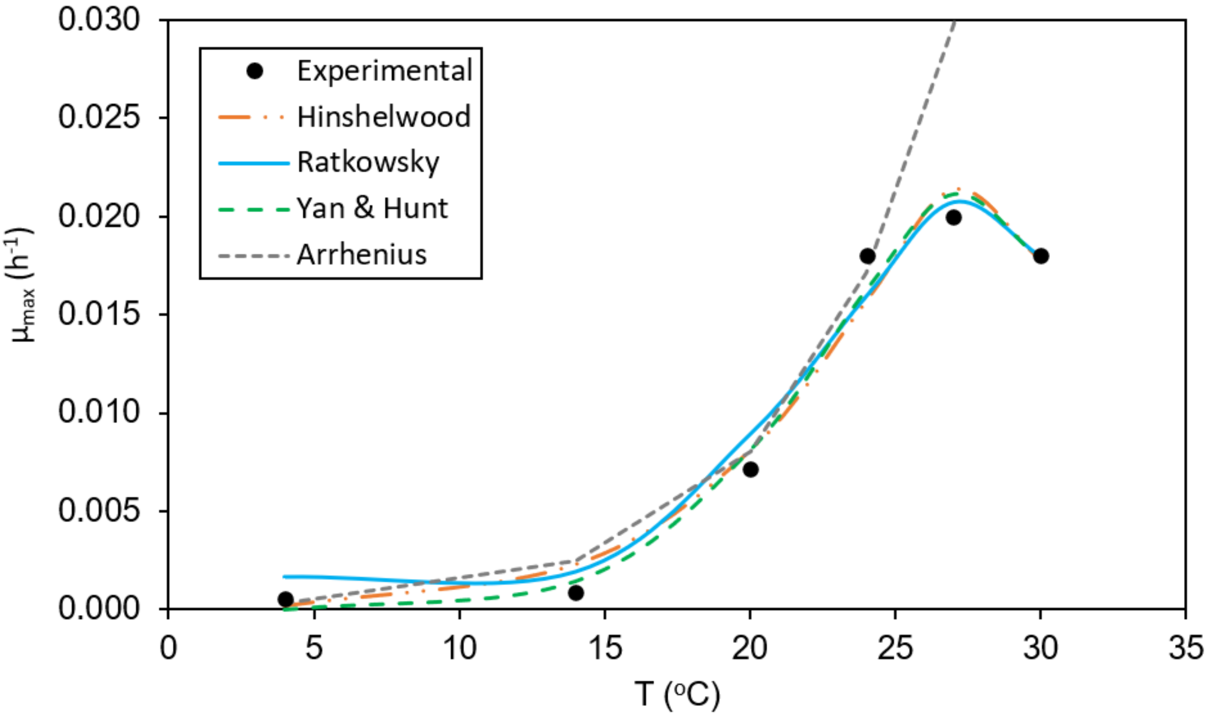


**Fig SM9. Maximum specific growth rate ($\mu_{max}$) as a function of temperature of *Nitrotoga*.** The fitting results to experimental data [57] using Hinshelwood model, modified Ratkowsky, Yan & Hunt equation and Arrhenius equation (S10 Table) are shown in coloured and different style of line.

**Influence of pH on growth rate**

To simulate the effect of pH on maximum specific growth rate ($\mu_{max}$), a bell curve expression was used (Eq S41) [37, 46, 53, 58-60].

$$\mu_{max}(pH) = \frac{\mu_{max}(pH_{op})}{1+10^{-pK1+pH}+10^{-pH+pK2}} \qquad \text{(S41)}$$

Where $pK1$ and $pK2$ refer to the pH in which $\mu_{max}$ is half of the value at optimum pH. Because the neutrophil AOB, *Nitrobacter* (NOB), *Nitrospira* (NOB) and *Nitrotoga* (NOB) have similar optimum pH range (Fig SM8B), the $pK1$ and $pK2$ values are from Jubany et al. (2008) study [53]. Due to the lack of information about optimum pH for AMX bacteria and the wide range of culture pH employed in AMX cultivation (Fig SM8B), it is assumed that there is no effect of pH on the growth rate of AMX.

**Selection of substrate affinities**

· **Oxygen affinity ($K_{O_2}$).** To select oxygen affinities of AOB and NOB, an exhausted analysis of $K_{O_2}$ values from literature was carried out. First, the ratio of $K_{O_2}$ values between NOB (*Nitrobacter* and *Nitrospira*) and AOB is calculated ($\mathrm{K_{O2}^{NOB}/K_{O2}^{AOB}}$) and represented on a histogram to see their frequencies (Fig 2B). When the ratio $\mathrm{K_{O2}^{NOB}/K_{O2}^{AOB}}$ is higher than 1, it means that AOB group has a higher oxygen affinity than NOB and vice versa.

In general, *Nitrobacter* tends to have a lower oxygen affinity than AOB (91% of calculated $\mathrm{K_{O2}^{Nb}/K_{O2}^{AOB}}$ is higher than 1.0). In contrast, is more frequent to find that *Nitrospira* have a higher oxygen affinity than AOB (56% of calculated $\mathrm{K_{O2}^{Ns}/K_{O2}^{AOB}}$ is lower than 1.0, with a maximum frequency of 14% in $\mathrm{K_{O2}^{Ns}/K_{O2}^{AOB}}$ = 0.2 – 0.3). It is important to consider that less $K_{O_2}$ values of *Nitrospira* were available in literature in comparison to AOB and *Nitrobacter* (*n* = 5 for *Nitrospira* (2007-2019), *n* = 19 for AOB (1993-2008), *n* = 17 for *Nitrobacter* (1989-2007)). This could influence the distribution of $\mathrm{K_{O2}^{NOB}/K_{O2}^{AOB}}$ ratios.

For this study, the $K_{O_2}$ values for AOB, *Nitrobacter* and *Nitrospira* have been chosen in function of the weighted averaged of $\mathrm{K_{O2}^{NOB}/K_{O2}^{AOB}}$ ratios shown in Fig 2B: 2.2 for *Nitrobacter*/AOB ratio ($\mathrm{K_{O2}^{Nb}/K_{O2}^{AOB}}$) and 0.72 for *Nitrospira*/AOB ratio ($\mathrm{K_{O2}^{Ns}/K_{O2}^{AOB}}$). To ensure that all $K_{O_2}$ values used are inside the range found in literature, the smallest value of *Nitrospira* is taken and then, the values for *Nitrobacter* and AOB are calculated with the mentioned ratios. Because no measured value of $K_{O_2}$ was found for *Nitrotoga*, the selection of the oxygen affinity of *Nitrotoga* was based on their biochemistry. The genome of *Ca.* N. fabula revealed that *Nitrotoga* encodes the high-affinity, proton-pumping heme-copper cytochrome *c* oxidase *cbb*$_3$-type [57]. Although, this terminal oxidase type have a higher intrinsic oxygen affinity than the terminal oxidase of *Nitrospira* (*bd*-like type) [19], the same value of $K_{O2}$ for *Nitrospira* and *Nitrotoga* was chosen.

· **Free nitrous acid affinity ($K_{FNA}$).** Nitrous acid affinities of *Nitrobacter* (Nb), *Nitrospira* (Ns) and *Nitrotoga* (Ng) are significantly different (Nb/Ns – $p < 0.001$; Nb/Ng – $p < 0.001$; Ns/Ng – $p < 0.02$). To set the $K_{FNA}$ values for NOB populations, it was first chosen the value for *Nitrobacter* from Keen and Prosser (1987) study [61]. Then, the $K_{FNA}$ value for *Nitrospira* was calculated based on the average ratio between *Nitrobacter* and *Nitrospira* ($\mathrm{K_{FNA}^{Ns}/K_{FNA}^{Nb}}$ = 0.02). For *Nitrotoga* and anammox bacteria, the $K_{FNA}$ values are from Ishii et al. (2017) [33] and van der Star et al. (2008) [24], respectively, the most favourable values (i.e., the highest reported affinity). This was done because the small number of measured values of $K_{FNA}$ for these two populations.

**Selection of inhibition constants**

· **Free ammonia inhibition of NOB ($K_{I,FA}$).** NOB is inhibited by free ammonia (FA, $NH_3$) [46]. Like nitrous acid affinities ($K_{FNA}$), FA tolerances of *Nitrobacter*, *Nitrospira* and *Nitrotoga* are significantly different. In general, *Nitrobacter* has a higher FA tolerance than *Nitrospira* and *Nitrotoga* [46, 62]. For *Nitrobacter* and *Nitrotoga*, the $K_{I,FA}$ values are from Kim et al. (2008) [63] and Li et al. (2020) [62], respectively. For

*Nitrospira*, different FA tolerance has been reported: 0.04 – 0.08 mg-$NH_3$/L (*Nitrospira sp.*) [59], 0.85 mg-$NH_3$/L (*Nitrospira sp.* strain ND1) [64] and 4.3 mg-$NH_3$/L (*Nitrospira japonica*) [64]. To obtain the non-competitive inhibition constant for *Nitrospira* ($K_{I,FA}$), the $\mathrm{K}_{I,NH3}^{\mathrm{Ns}}/\mathrm{K}_{I,NH3}^{\mathrm{Nb}}$ ratios were calculated using the FA tolerance of *Nitrobacter* from Blackburne et al. (2007) study [46]: $\mathrm{K}_{I,NH3}^{\mathrm{Ns}}/\mathrm{K}_{I,NH3}^{\mathrm{Nb}}$ = 0.0015 [46]; $\mathrm{K}_{I,NH3}^{\mathrm{Ns}}/\mathrm{K}_{I,NH3}^{\mathrm{Nb}}$ = 0.017 [64]; $\mathrm{K}_{I,NH3}^{\mathrm{Ns}}/\mathrm{K}_{I,NH3}^{\mathrm{Nb}}$ = 0.104 [64]. Then, the non-competitive inhibition constant of *Nitrobacter* from Kim et al. (2008) study was used to calculate the $K_{I,FA}$ values: 0.032 mg-$NH_3$/L (*Nitrospira sp.*), 0.362 mg-$NH_3$/L (*Nitrospira sp.* strain ND1) and 2.210 mg-$NH_3$/L (*N. japonica*). For the main set of simulations, the most favourable value (i.e., the highest $K_{I,FA}$ value) was chosen (2.210 mg-$NH_3$/L). The effect of free ammonia inhibition of *Nitrospira* will be evaluated, modifying the $K_{I,FA}^{Ns}$ value.

**Summary of the selection of kinetic parameters**

Here below is shown a recap of the kinetic selection (Fig SM10). All selected kinetic parameters and expressions are summarized in S2 Table.

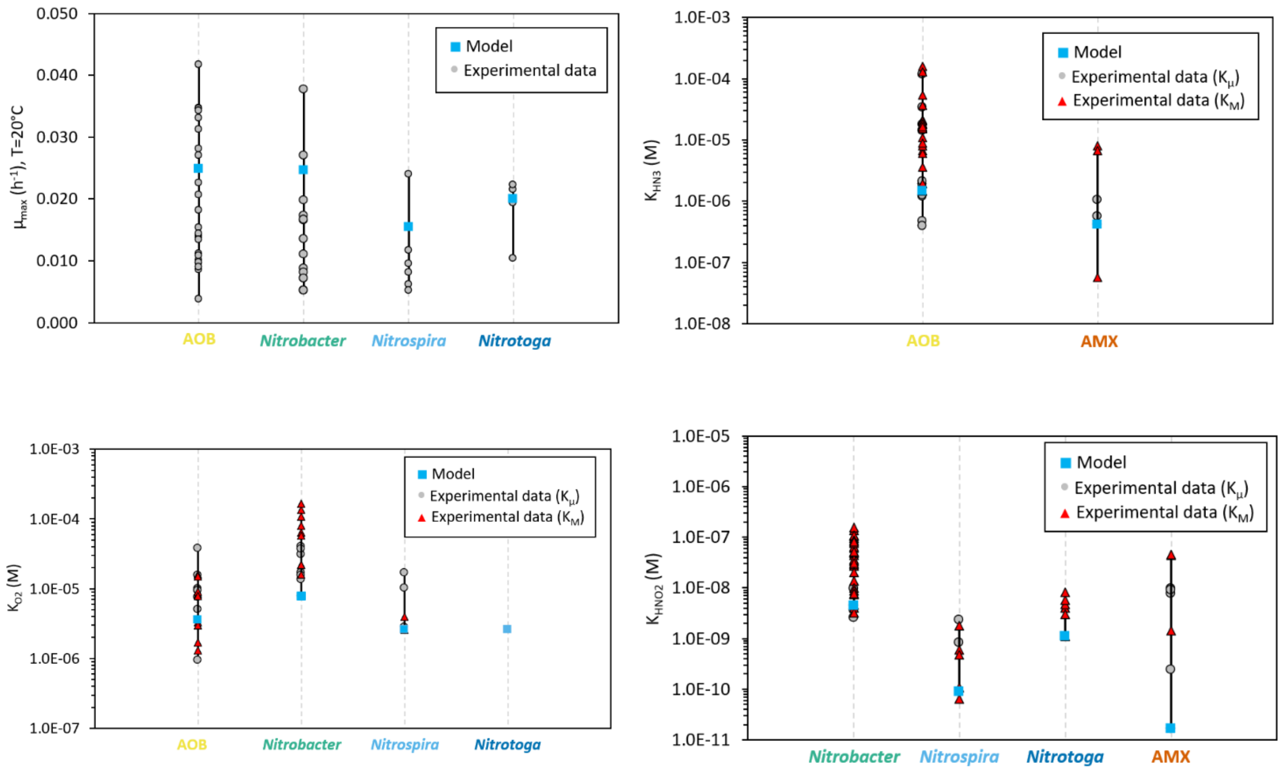


**Fig SM10. Summary of kinetic parameters of AOB, NOB (*Nitrobacter*, *Nitrospira* and *Nitrotoga*) and AMX considered in this study.** For all substrate affinities, Monod constant (growth rate; $K_\mu$) and Michaelis-Menten constant (uptake rate; $K_M$) were differentiated.
