## Supplementary figures and images for "Physiological diversity and the assembly of nitrifying communities in aerobic granular sludge"

### S1 Figure

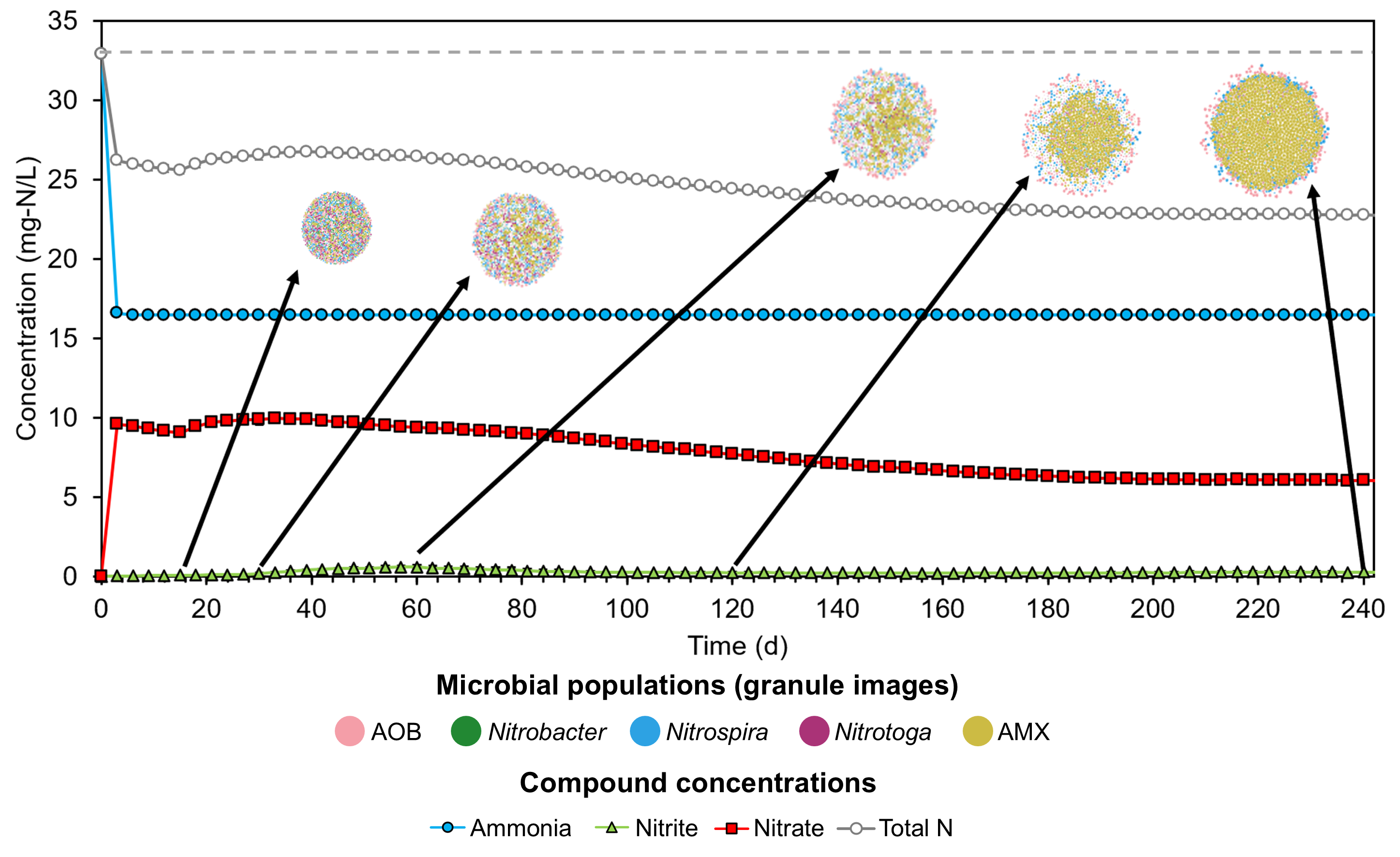

### S2 Figure

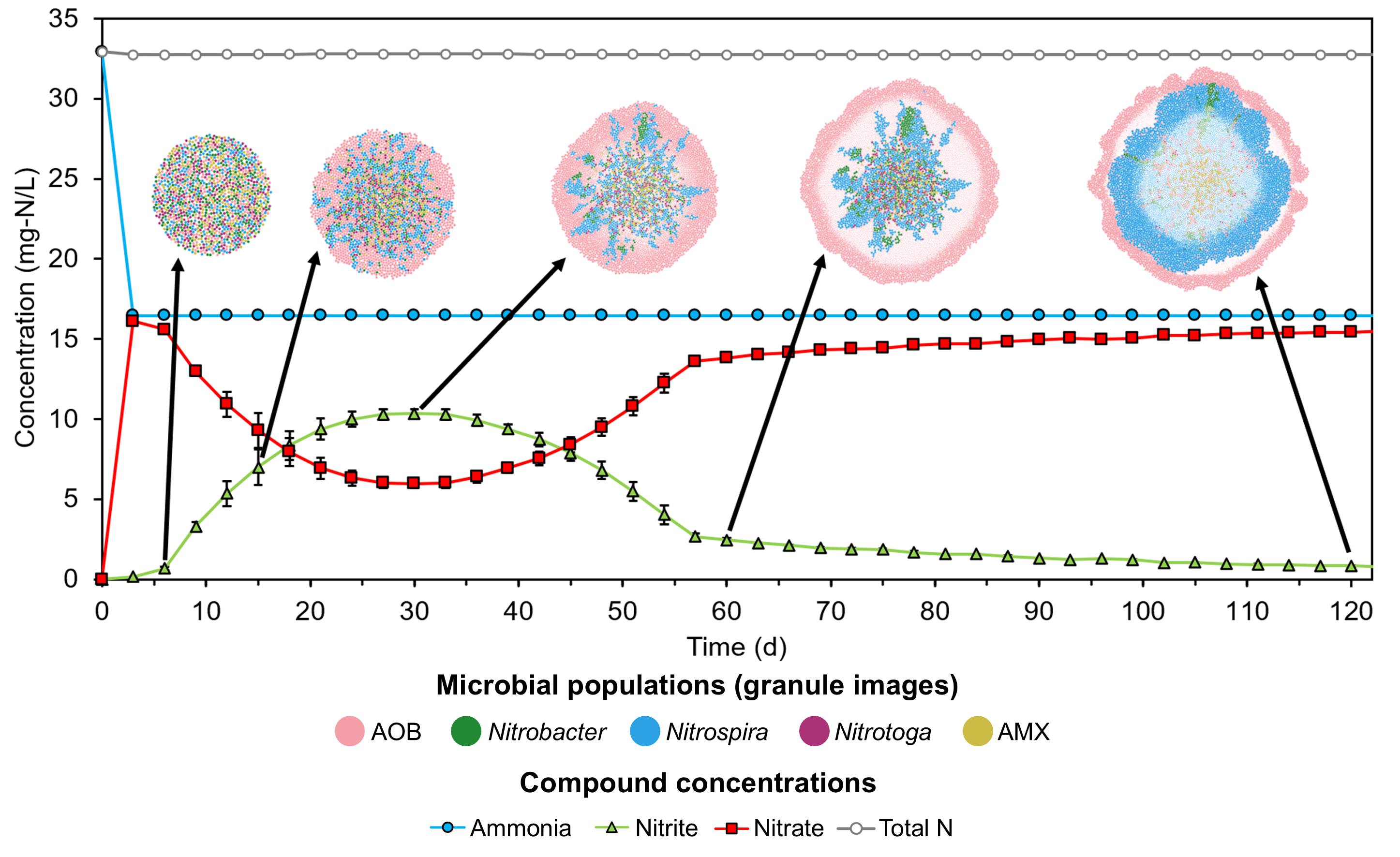

### S3 Figure

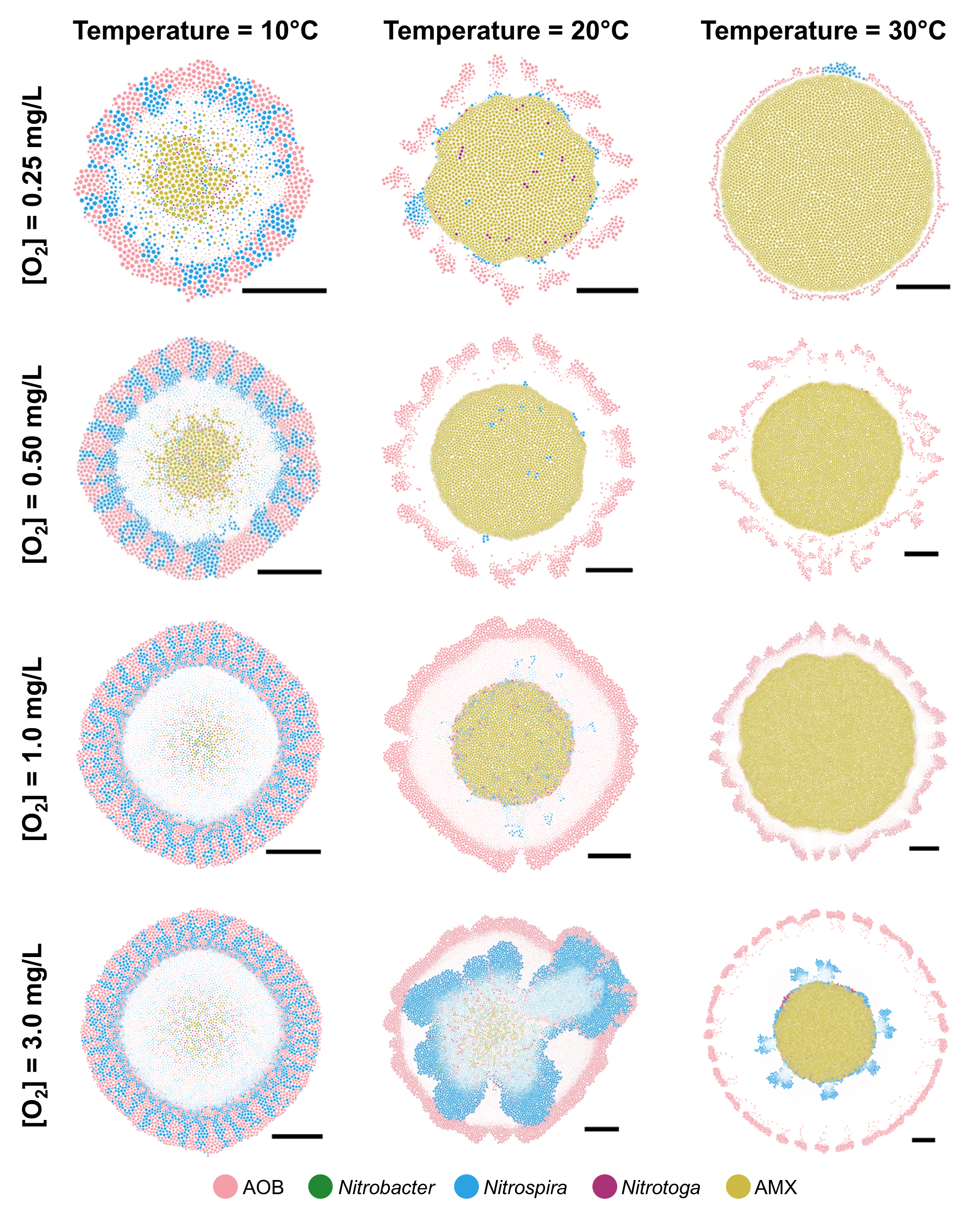

### S4 Figure

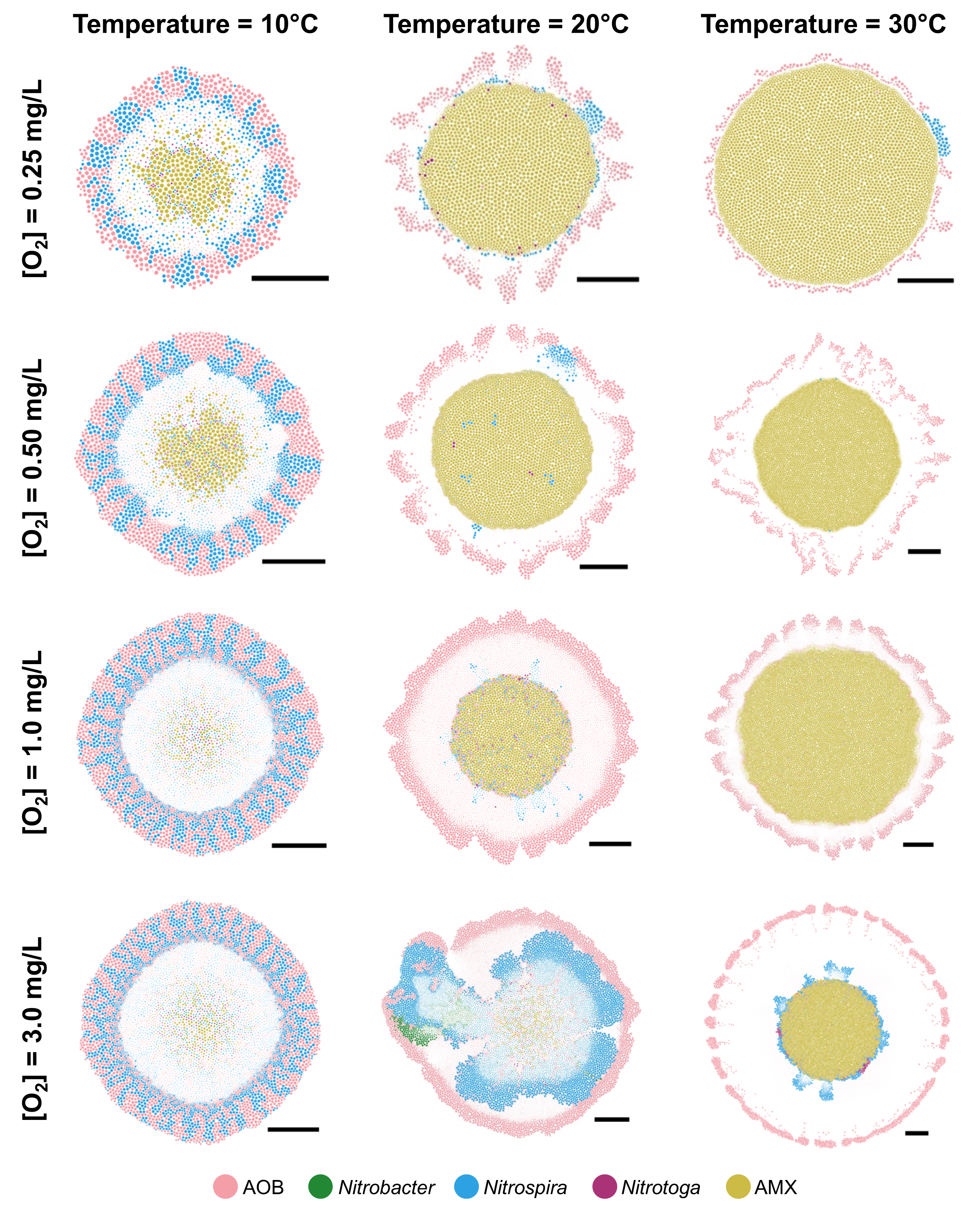

### S5 Figure

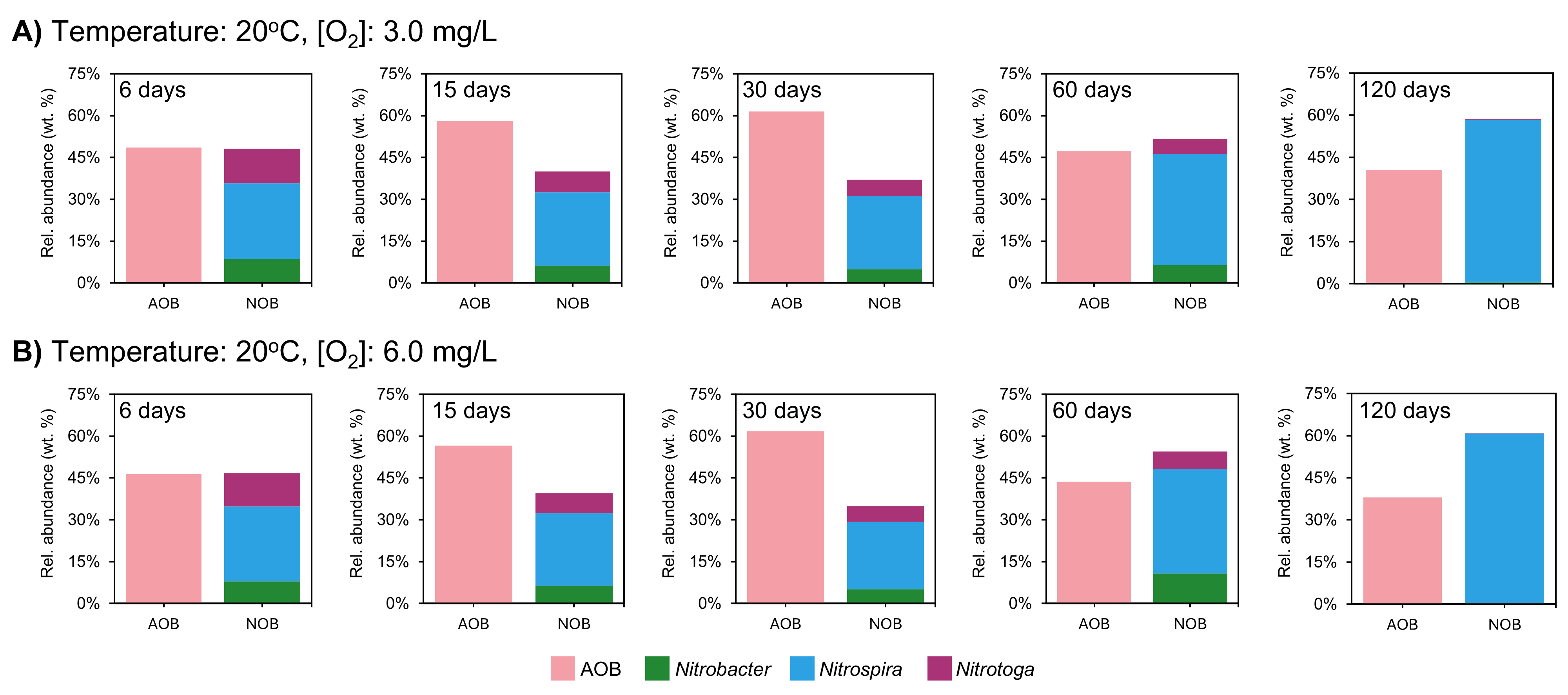

### S6 Figure

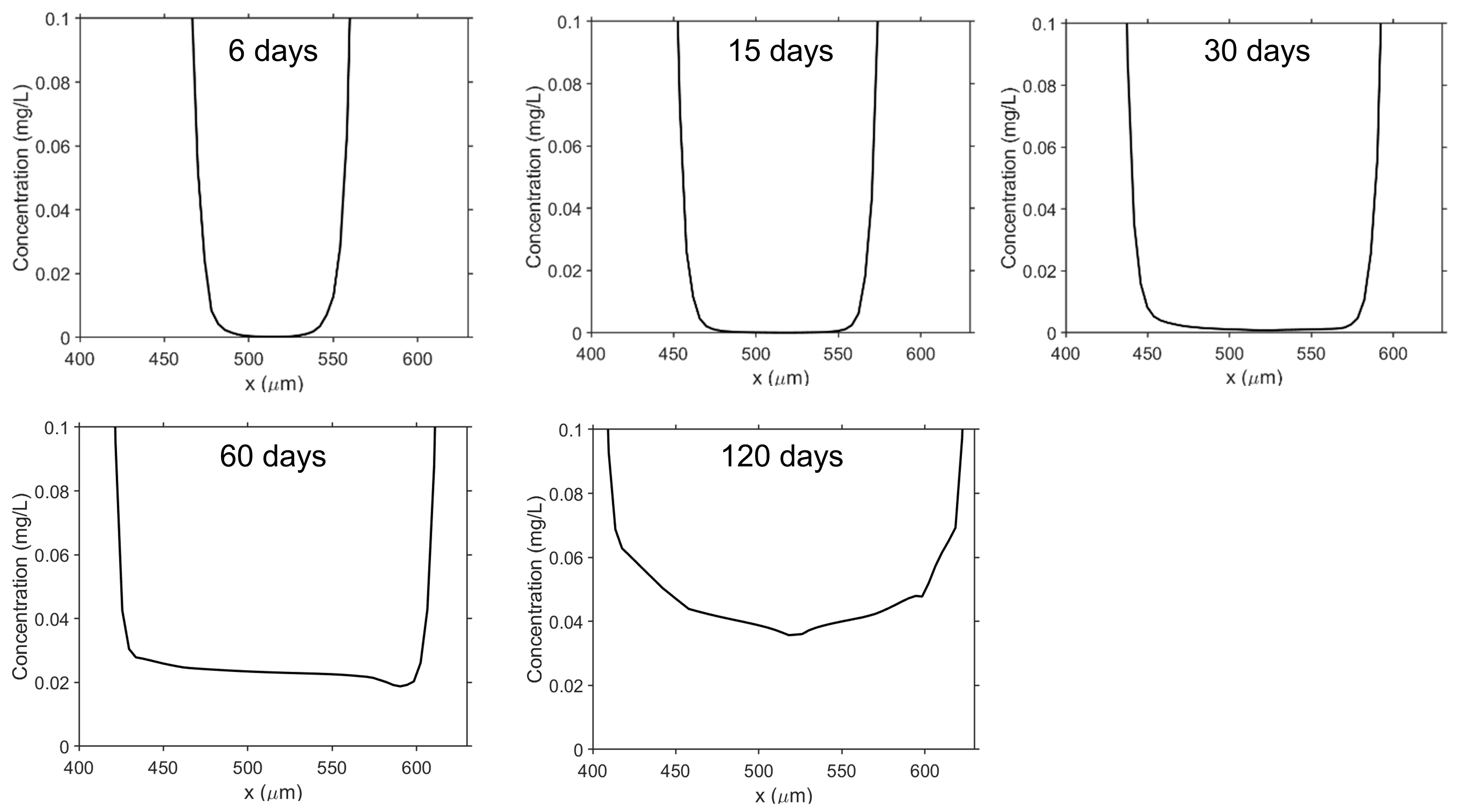

### S7 Figure

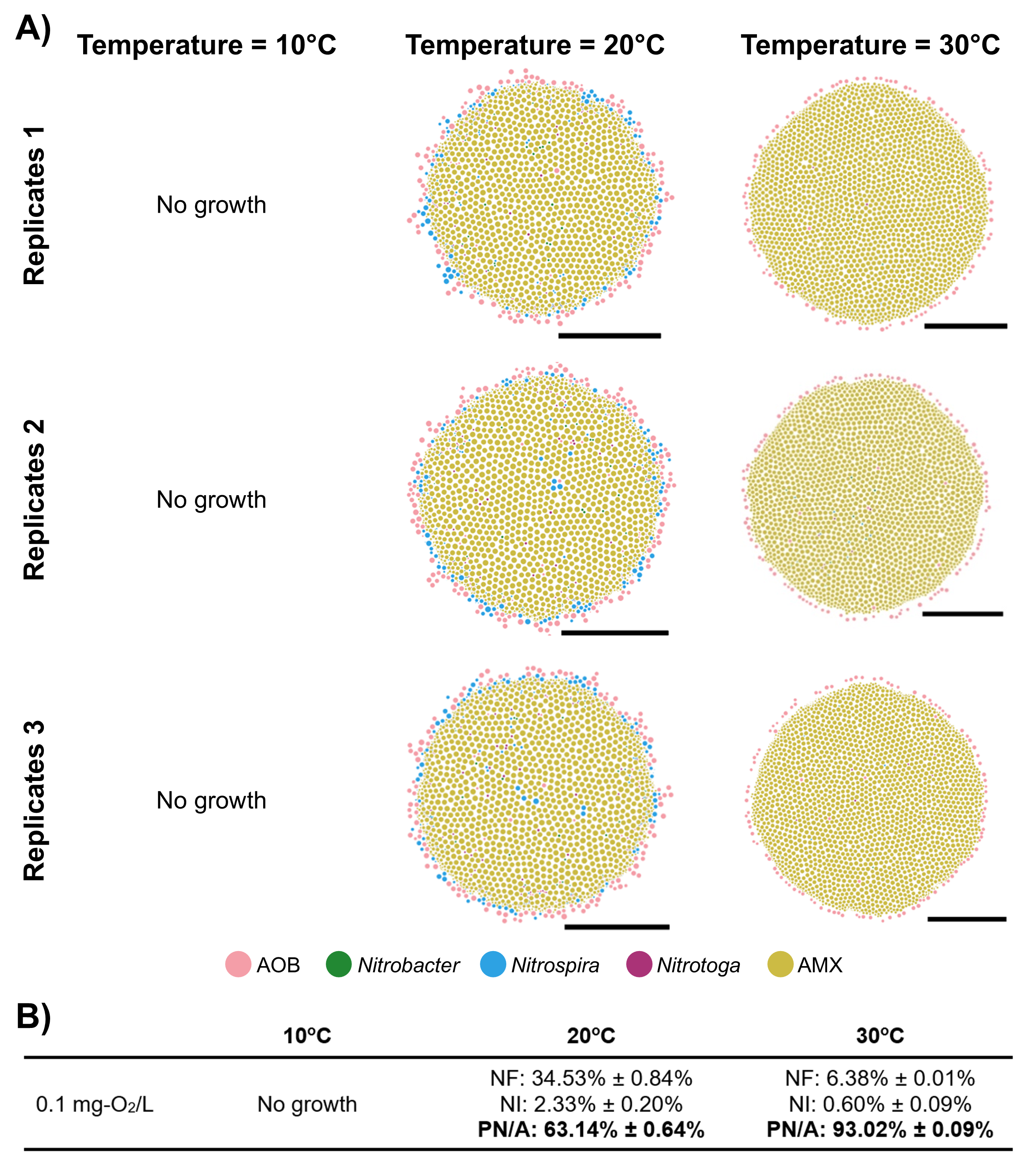

### S8 Figure

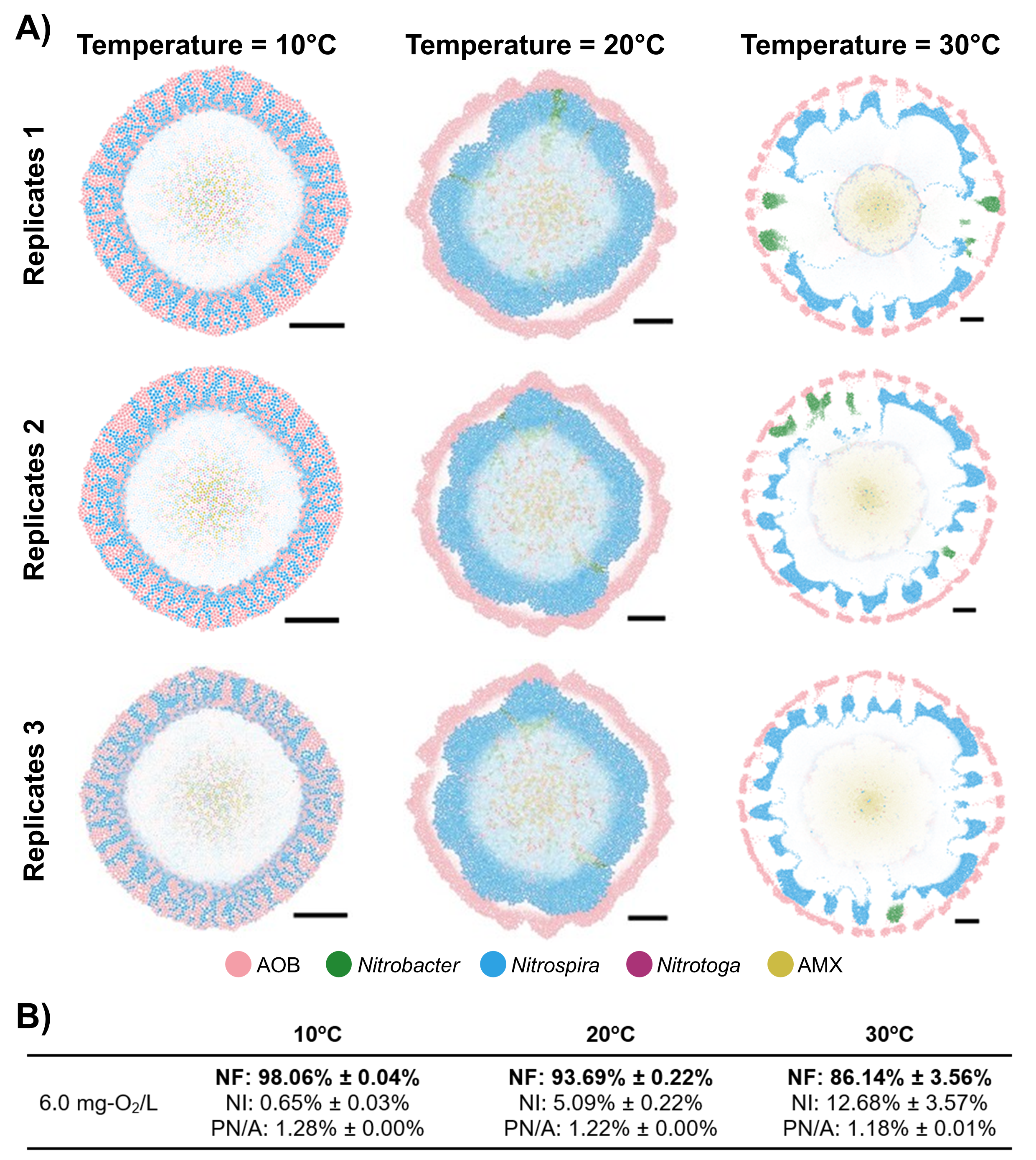

### S9 Figure

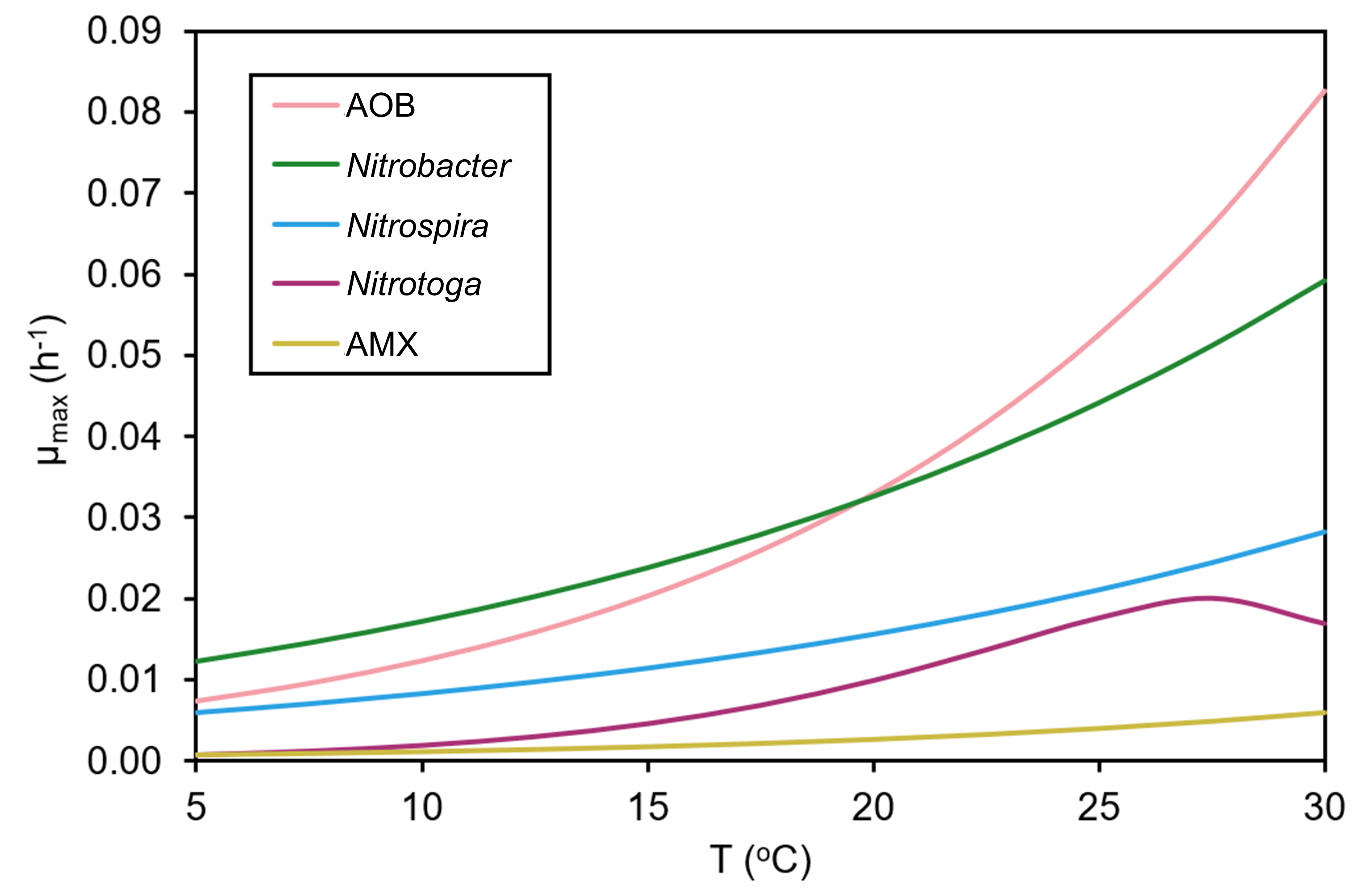

### S10 Figure

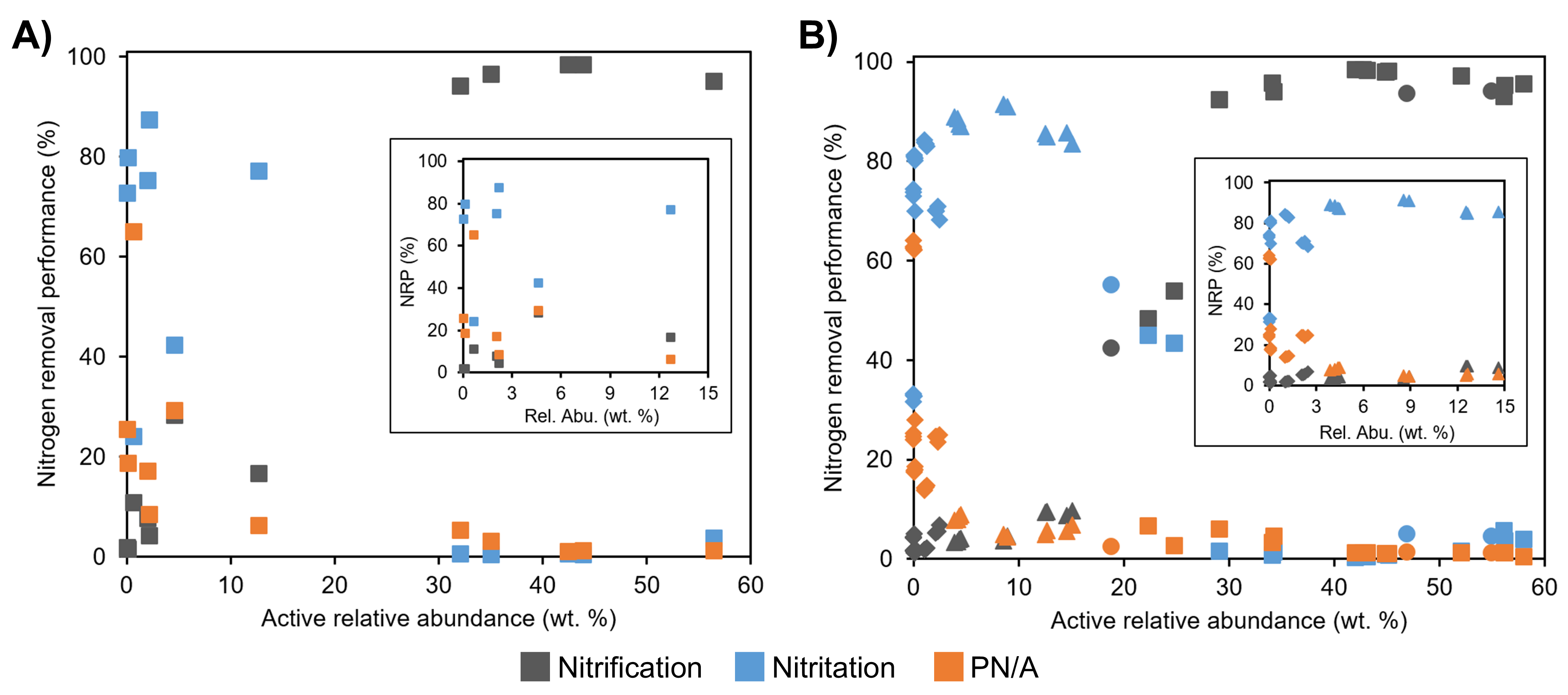

### S11 Figure

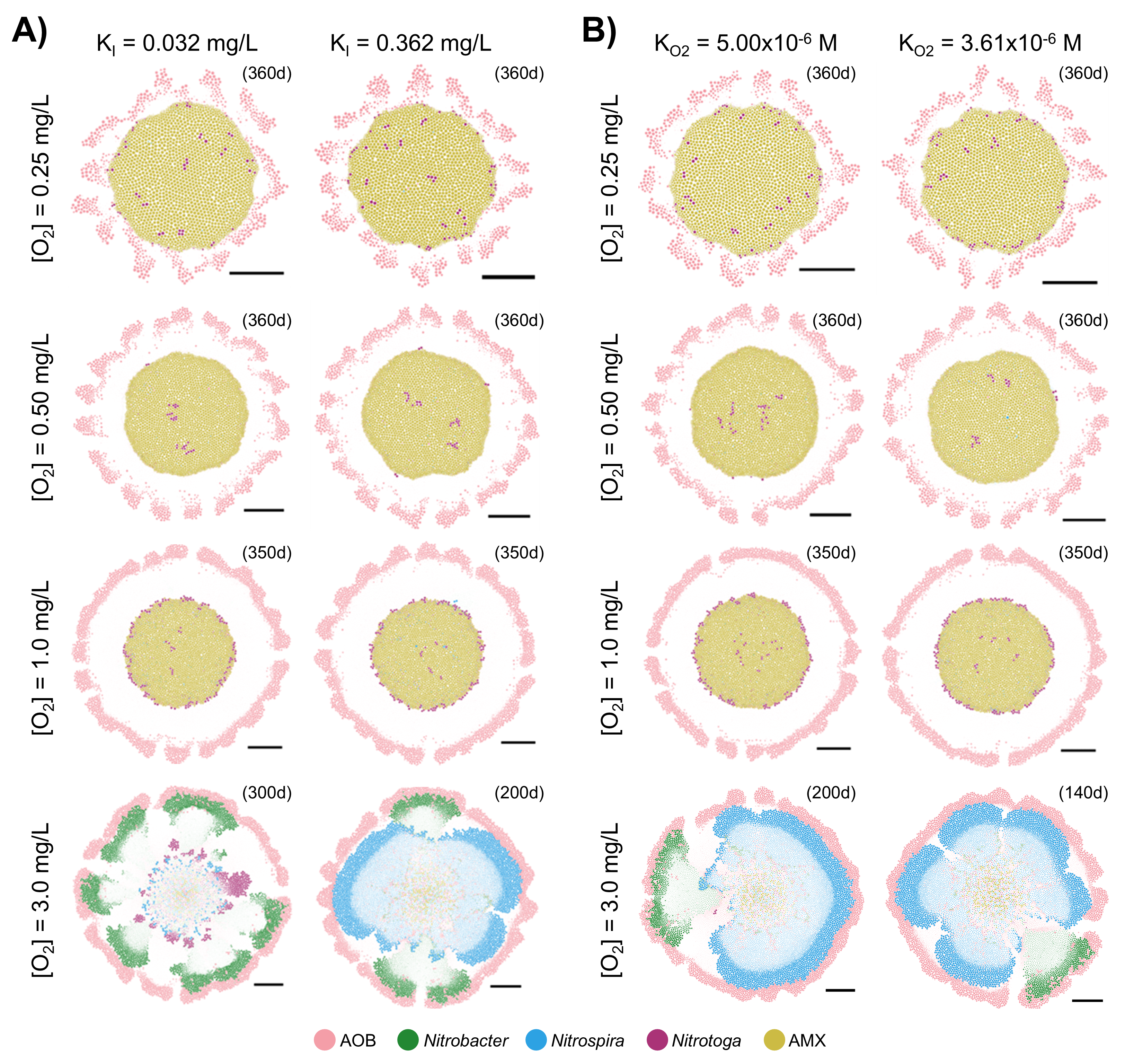

### S12 Figure

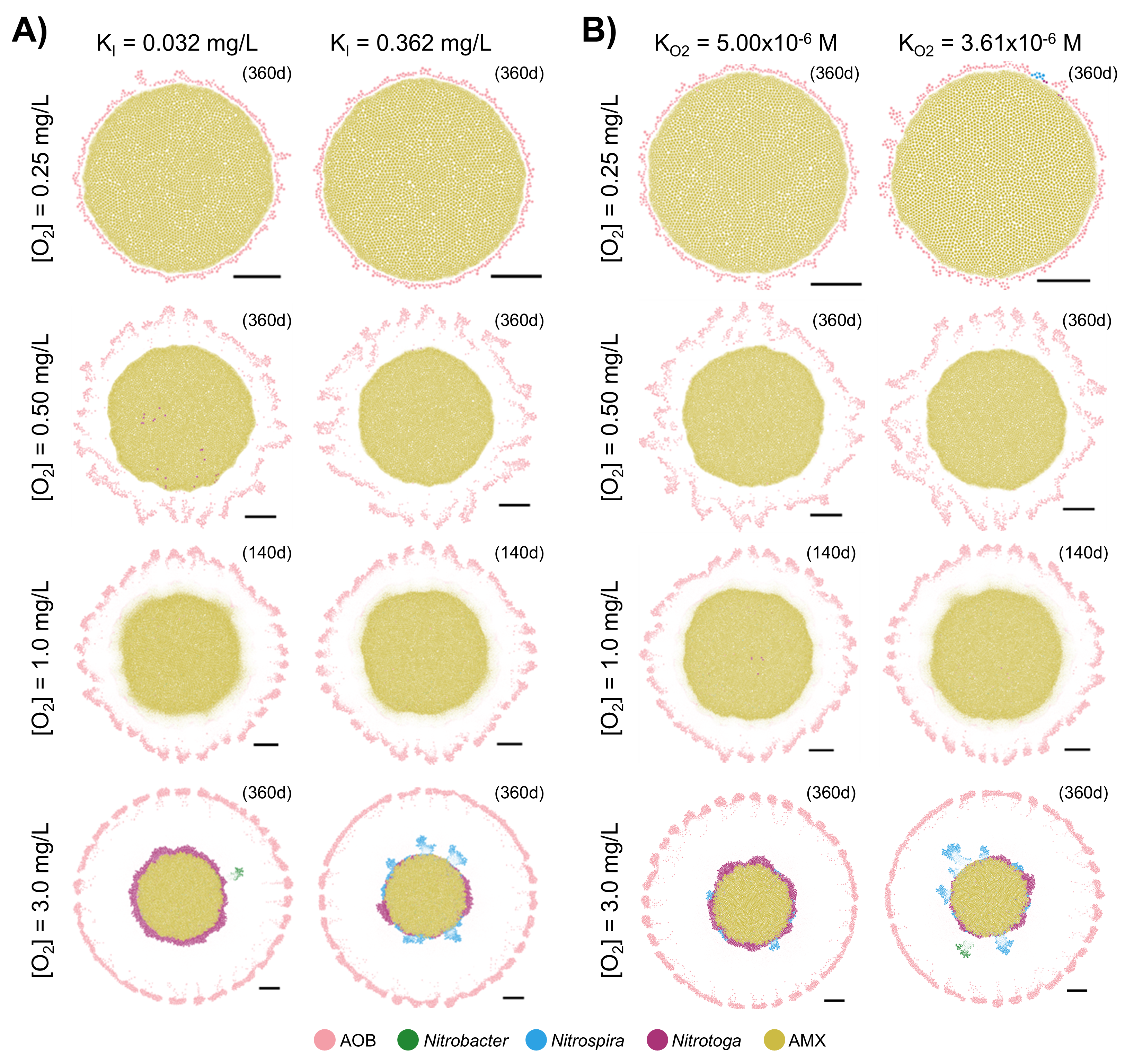

### S13 Figure

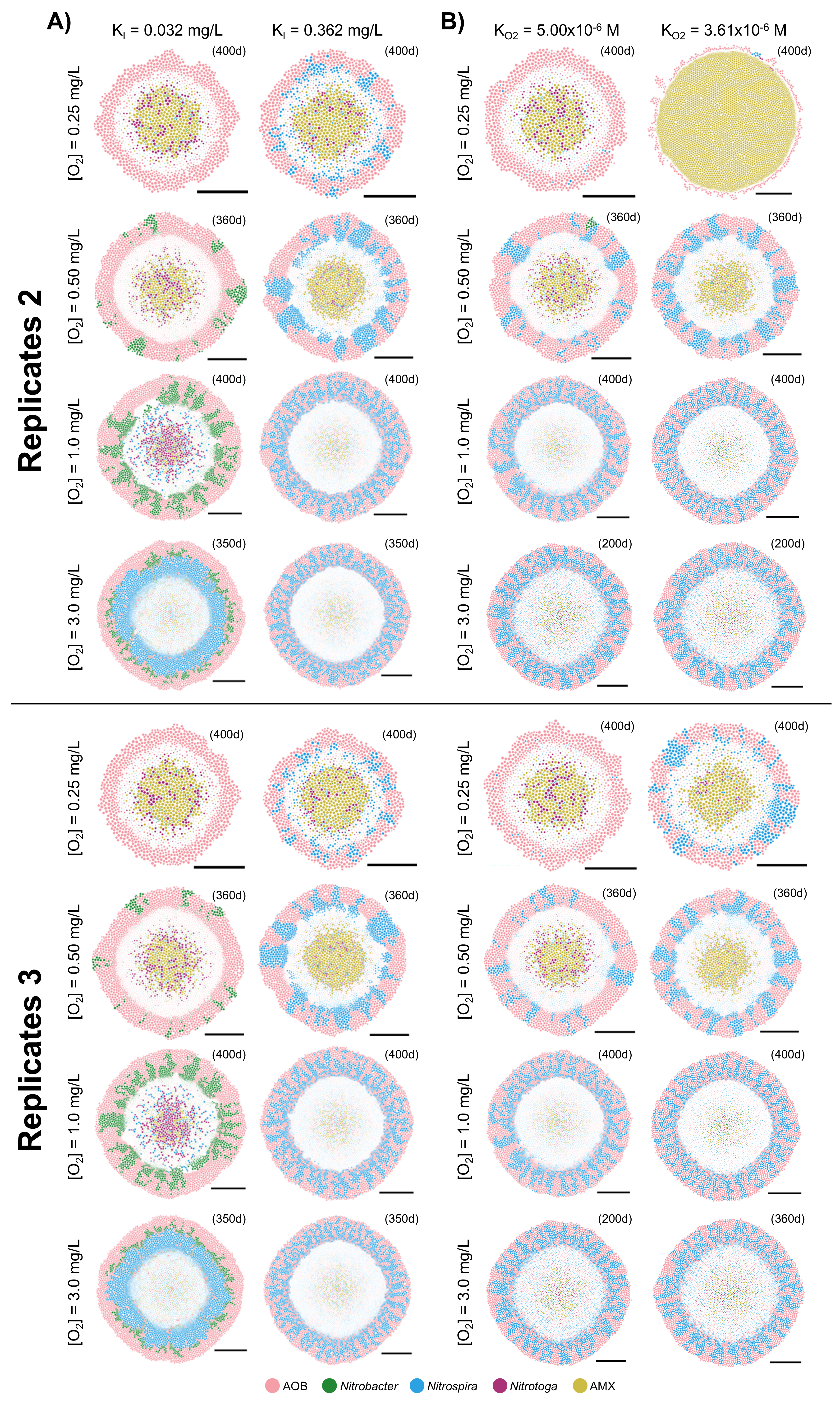

### S14 Figure

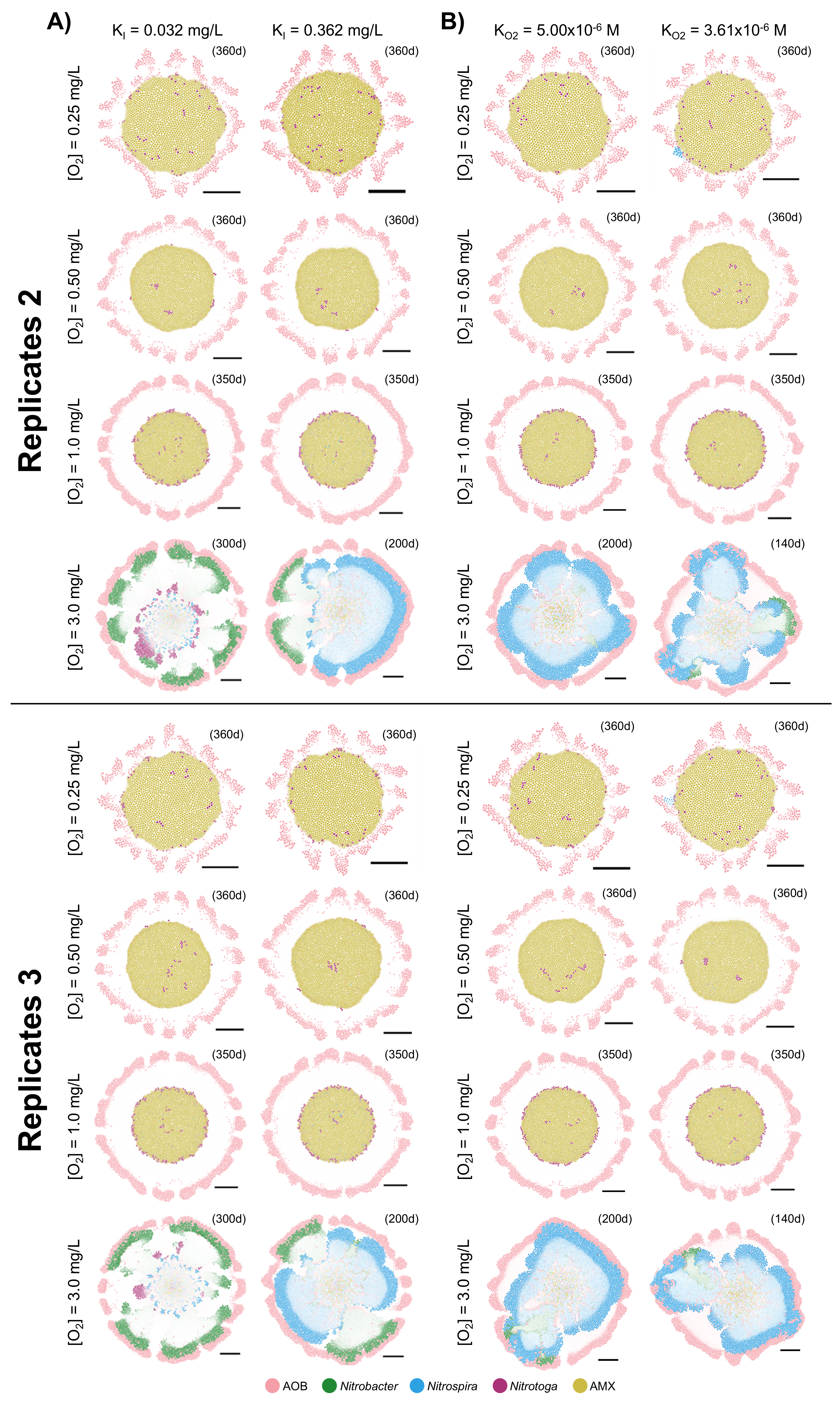

### S15 Figure

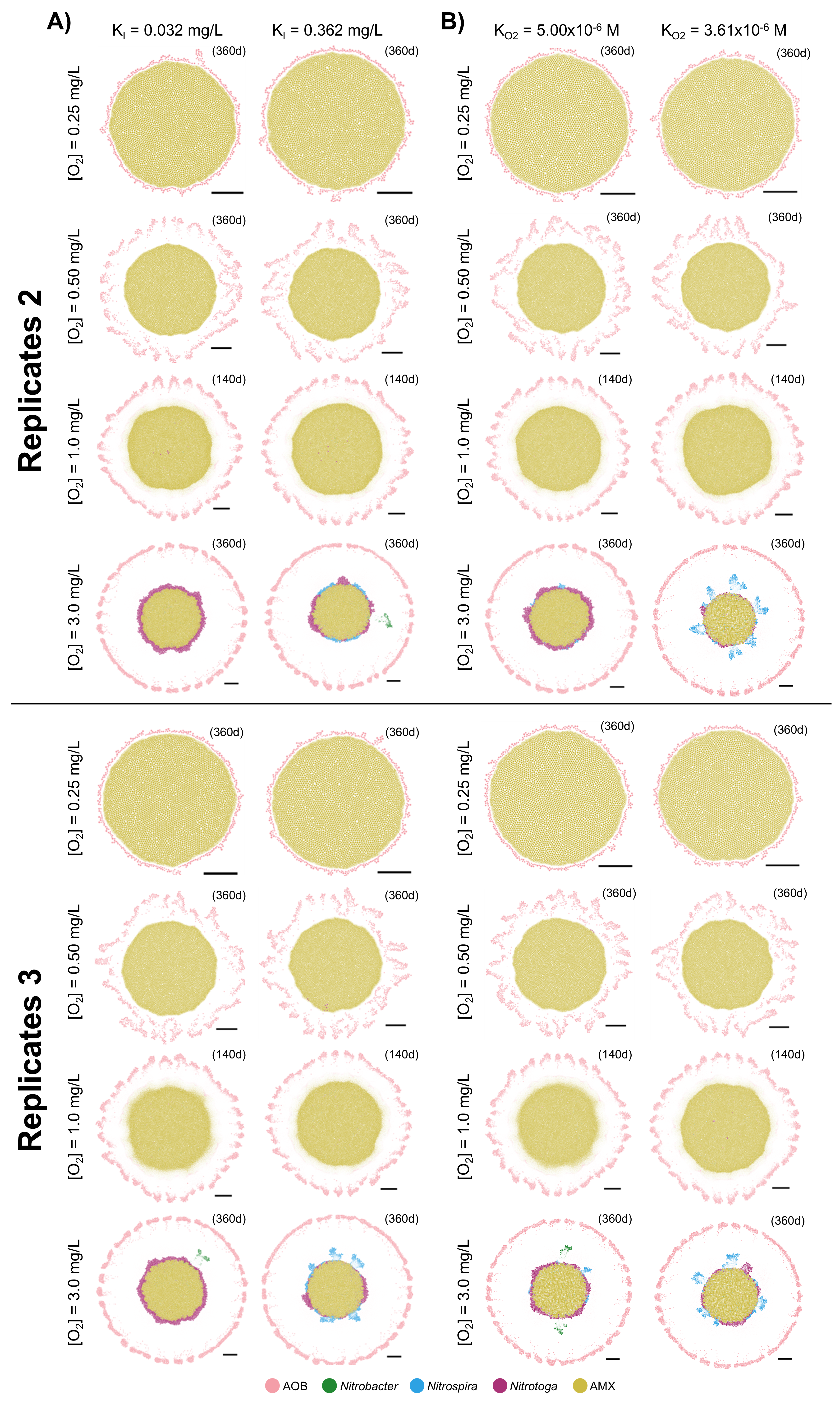
