## Supplementary material for "Physiological diversity and the assembly of nitrifying communities in aerobic granular sludge": S1 Table

**S1 Table. Summary of conditions of simulation experiments.** Symbols: $[S]_{sp}$ – setpoint concentration; $[S]_{in}$ – influent concentration.

| | T (ºC) | $[O_2]_{sp}$ (mg/L) | $[NH_3]_{sp}$ (mg/L) | $[NH_3]_{in}$ (mg/L) | pH (–) |
|---|---|---|---|---|---|
| **Base case** | | | | | |
| Poot et al. (2016) | 20 | 1.0 | 20 | 40 | 7.7 |
| **Evaluation of $[O_2]$ influence on base case** | | | | | |
| $[O_2]$ setpoint nº1 | 20 | 0.1 | 20 | 40 | 7.7 |
| $[O_2]$ setpoint nº2 | 20 | 0.25 | 20 | 40 | 7.7 |
| $[O_2]$ setpoint nº3 | 20 | 0.5 | 20 | 40 | 7.7 |
| $[O_2]$ setpoint nº4 | 20 | 3.0 | 20 | 40 | 7.7 |
| $[O_2]$ setpoint nº5 | 20 | 6.0 | 20 | 40 | 7.7 |
| **Evaluation of temperature influence under $[O_2]_{sp}$ = 0.1 mg/L** | | | | | |
| Temperature nº1 | 10 | 0.1 | 20 | 40 | 7.7 |
| Temperature nº2 | 30 | 0.1 | 20 | 40 | 7.7 |
| **Evaluation of temperature influence under $[O_2]_{sp}$ = 0.25 mg/L** | | | | | |
| Temperature nº1 | 10 | 0.25 | 20 | 40 | 7.7 |
| Temperature nº2 | 30 | 0.25 | 20 | 40 | 7.7 |
| **Evaluation of temperature influence under $[O_2]_{sp}$ = 0.5 mg/L** | | | | | |
| Temperature nº1 | 10 | 0.5 | 20 | 40 | 7.7 |
| Temperature nº2 | 30 | 0.5 | 20 | 40 | 7.7 |
| **Evaluation of temperature influence under $[O_2]_{sp}$ = 1.0 mg/L** | | | | | |
| Temperature nº1 | 10 | 1.0 | 20 | 40 | 7.7 |
| Temperature nº2 | 30 | 1.0 | 20 | 40 | 7.7 |
| **Evaluation of temperature influence under $[O_2]_{sp}$ = 3.0 mg/L** | | | | | |
| Temperature nº1 | 10 | 3.0 | 20 | 40 | 7.7 |
| Temperature nº2 | 30 | 3.0 | 20 | 40 | 7.7 |
| **Evaluation of temperature influence under $[O_2]_{sp}$ = 6.0 mg/L** | | | | | |
| Temperature nº1 | 10 | 6.0 | 20 | 40 | 7.7 |
| Temperature nº2 | 30 | 6.0 | 20 | 40 | 7.7 |
