## Supplementary material for "Physiological diversity and the assembly of nitrifying communities in aerobic granular sludge": S2 Table

**S2 Table. Summary of kinetic parameters of AOB, NOB (*Nitrobacter, Nitrospira*, *Nitrotoga*) and AMX.** EST – estimated. For more details, see S3 Appendix – *Physiological definition of microbial populations*.

| Parameter | Symbol | Value/Equation | Units | REF |
|---|---|---|---|---|
| **Ammonia-oxidizing bacteria (AOB)** | | | | |
| Max. specific growth rate | $\mu_{max}^{AOB}$ | $\frac{5.333 \times 10^{10} * \exp\left(-\frac{8183}{T}\right)}{1 + 10^{-8.688+pH} + 10^{-pH+6.780}}$ | $h^{-1}$ | [1] |
| Specific maintenance rate | $a^{AOB}$ | $6.879 \times 10^{9} * \exp\left(-\frac{8183}{T}\right)$ | $h^{-1}$ | [1] |
| Growth yield | $Y_{XS}^{AOB}$ | $0.0409$ | $mol_{Cx}/mol_{NH3}$ | [2] |
| Ammonia affinity (FA) | $K_{S,FA}^{AOB}$ | $1.453 \times 10^{-6}$ | mol $L^{-1}$ | [3] |
| Oxygen affinity | $K_{S,O_2}^{AOB}$ | $3.606 \times 10^{-6}$ | mol $L^{-1}$ | [4] |
| Nitrous acid inhibition (FNA) | $K_{I,FNA}^{AOB}$ | $1.170 \times 10^{-5}$ | mol $L^{-1}$ | [1] |
| **Nitrite-oxidizing bacteria, *Nitrobacter* (Nb)** | | | | |
| Max. specific growth rate | $\mu_{max}^{Nb}$ | $\frac{2.788 \times 10^{6} * \exp\left(-\frac{5295}{T}\right)}{1 + 10^{-8.688+pH} + 10^{-pH+6.780}}$ | $h^{-1}$ | [1] |
| Specific maintenance rate | $a^{Nb}$ | $3.594 \times 10^{5} * \exp\left(-\frac{5295}{T}\right)$ | $h^{-1}$ | [1] |
| Growth yield | $Y_{XS}^{Nb}$ | $0.0242$ | $mol_{Cx}/mol_{NO2}$ | [2] |
| Nitrous acid affinity (FNA) | $K_{S,FNA}^{Nb}$ | $4.447 \times 10^{-9}$ | mol $L^{-1}$ | [5] |
| Oxygen affinity | $K_{S,O_2}^{Nb}$ | $7.813 \times 10^{-6}$ | mol $L^{-1}$ | [6] |
| Ammonia inhibition (FA) | $K_{I,FA}^{Nb}$ | $1.521 \times 10^{-3}$ | mol $L^{-1}$ | [7] |
| **Nitrite-oxidizing bacteria, *Nitrospira* (Ns)** | | | | |
| Max. specific growth rate | $\mu_{max}^{Ns}$ | $\frac{1.756 \times 10^{6} * \exp\left(-\frac{5295}{T}\right)}{1 + 10^{-8.688+pH} + 10^{-pH+6.780}}$ | $h^{-1}$ | [1] |
| Specific maintenance rate | $a^{Ns}$ | $2.264 \times 10^{5} * \exp\left(-\frac{5295}{T}\right)$ | $h^{-1}$ | [1] |
| Growth yield | $Y_{XS}^{Ns}$ | $0.0242$ | $mol_{Cx}/mol_{NO2}$ | [2] |
| Nitrous acid affinity (FNA) | $K_{S,FNA}^{Ns}$ | $8.894 \times 10^{-11}$ | mol $L^{-1}$ | EST |
| Oxygen affinity | $K_{S,O_2}^{Ns}$ | $2.594 \times 10^{-6}$ | mol $L^{-1}$ | [8] |
| Ammonia inhibition (FA) | $K_{I,FA}^{Ns}$ | $1.300 \times 10^{-4}$ | mol $L^{-1}$ | [7, 8] |

**Table S1.** ***Continuation…***

| Parameter | Symbol | Value/Equation | Units | REF |
|---|---|---|---|---|
| **Nitrite-oxidizing bacteria, *Nitrotoga* (Ng)** | | | | |
| Max. specific growth rate | $\mu_{max}^{Ng}$ | $\frac{0.025 \cdot \left(\frac{305.6 - T}{4.67}\right)\left(\frac{T}{300.93}\right)^{64.44}}{1 + 10^{-8.688+pH} + 10^{-pH+6.780}}$ | $h^{-1}$ | [9] |
| Specific maintenance rate | $a^{Ng}$ | $3.225 \times 10^{-3}\left(\frac{305.6 - T}{4.67}\right)\left(\frac{T}{300.93}\right)^{64.44}$ | $h^{-1}$ | EST |
| Growth yield | $Y_{XS}^{Ng}$ | $0.0242$ | $mol_{Cx}/mol_{NO2}$ | [2] |
| Nitrous acid affinity (FNA) | $K_{S,FNA}^{Ng}$ | $1.119 \times 10^{-9}$ | mol $L^{-1}$ | [10] |
| Oxygen affinity | $K_{S,O_2}^{Ng}$ | $2.594 \times 10^{-6}$ | mol $L^{-1}$ | EST |
| Ammonia inhibition (FA) | $K_{I,FA}^{Ng}$ | $8.643 \times 10^{-4}$ | mol $L^{-1}$ | [11] |
| **Anammox bacteria (AMX)** | | | | |
| Max. specific growth rate | $\mu_{max}^{AMX}$ | $1.89 \times 10^{8} * \exp\left(-\frac{7330}{T}\right)$ | $h^{-1}$ | [12, 13] |
| Specific maintenance rate | $a_{max}^{AMX}$ | $5\% * \mu_{max}^{AMX}$ | $h^{-1}$ | [14] |
| Growth yield | $Y_{XS}^{AMX}$ | $0.066$ | $mol_{Cx}/mol_{NH3}$ | [2] |
| Ammonia affinity (FA) | $K_{S,FA}^{AMX}$ | $4.118 \times 10^{-7}$ | mol $L^{-1}$ | [15] |
| Nitrous acid affinity (FNA) | $K_{S,FNA}^{AMX}$ | $1.632 \times 10^{-11}$ | mol $L^{-1}$ | [16] |
| Oxygen inhibition | $K_{I,O_2}^{AMX}$ | $3.125 \times 10^{-7}$ | mol $L^{-1}$ | [17] |
