## Supplementary material for "Physiological diversity and the assembly of nitrifying communities in aerobic granular sludge": S3 Table

**S3 Table. Maximum portion values of nitrogen conversion process.** Symbol legend: NF – nitrification, NI – nitritation, PN/A – partial nitritation + anammox. Standard deviations are from three replicates.

| | 10°C | 20°C | 30°C |
|---|---|---|---|
| 0.1 mg-$O_2$/L | No growth | NF: 60.52% ± 1.59%<br>NI: 3.63% ± 0.65%<br>PN/A: 63.75% ± 0.37% | NF: 38.84% ± 1.25%<br>NI: 4.40% ± 0.22%<br>PN/A: 93.32% ± 0.03% |
| 0.25 mg-$O_2$/L | NF: 95.01% ± 0.08%<br>NI: 0.58% ± 0.02%<br>PN/A: 5.67% ± 0.08% | NF: 80.97% ± 1.65%<br>NI: 44.78% ± 18.74%<br>PN/A: 32.35% ± 2.40% | NF: 55.33% ± 0.40%<br>NI: 24.35% ± 6.63%<br>PN/A: 73.20% ± 0.56% |
| 0.5 mg-$O_2$/L | NF: 97.63% ± 0.05%<br>NI: 0.68% ± 0.03%<br>PN/A: 5.12% ± 0.40% | NF: 89.46% ± 0.83%<br>NI: 76.87% ± 6.36%<br>PN/A: 24.94% ± 1.05% | NF: 56.66% ± 2.44%<br>NI: 72.83% ± 1.55%<br>PN/A: 60.94% ± 1.55% |
| 1.0 mg-$O_2$/L | NF: 98.74% ± 0.01%<br>NI: 0.79% ± 0.08%<br>PN/A: 1.44% ± 0.00% | NF: 93.14% ± 1.08%<br>NI: 93.77 ± 0.23%<br>PN/A: 11.56% ± 0.12% | NF: 38.69% ± 8.54%<br>NI: 82.02% ± 1.89%<br>PN/A: 55.34% ± 1.27% |
| 3.0 mg-$O_2$/L | NF: 98.73% ± 0.01%<br>NI: 0.99% ± 0.30%<br>PN/A: 1.29% ± 0.00% | NF: 97.59% ± 0.19%<br>NI: 56.69% ± 5.43%<br>PN/A: 1.67% ± 0.32% | NF: 45.36% ± 0.74%<br>NI: 97.11% ± 0.02%<br>PN/A: 9.12% ± 0.30% |
| 6.0 mg-$O_2$/L | NF: 98.71% ± 0.01%<br>NI: 1.40% ± 0.04%<br>PN/A: 1.29% ± 0.00% | NF: 97.80% ± 0.11%<br>NI: 62.84% ± 1.48%<br>PN/A: 1.25% ± 0.00% | NF: 88.25% ± 0.56%<br>NI: 96.71% ± 0.11%<br>PN/A: 12.67% ± 6.78% |
