## Supplementary material for "Physiological diversity and the assembly of nitrifying communities in aerobic granular sludge": S4 Table

**S4 Table. Statistical significance from comparison of nitrogen process partitioning considering different ammonia tolerance of *Nitrospira* ($K_I$, mg-$NH_3$/L) A)** [$O_2$] = 0.25 mg/L; 10°C. **B)** [$O_2$] = 0.25 mg/L; 20°C. **C)** [$O_2$] = 0.25 mg/L; 30°C. **D)** [$O_2$] = 0.5 mg/L; 10°C. **E)** [$O_2$] = 0.5 mg/L; 20°C. **F)** [$O_2$] = 0.5 mg/L; 30°C. **G)** [$O_2$] = 1.0 mg/L; 10°C. **H)** [$O_2$] = 1.0 mg/L; 20°C. **I)** [$O_2$] = 1.0 mg/L; 30°C. **J)** [$O_2$] = 3.0 mg/L; 10°C. **K)** [$O_2$] = 3.0 mg/L; 20°C. **L)** [$O_2$] = 3.0 mg/L; 30°C. Symbol legend: ns – not significant; * – $p < 0.05$; ** – $p < 0.01$; *** – $p < 0.001$.

| | **Ammonia tolerance of *Nitrospira*** | **Nitrification** | **Nitritation** | **PN/A** |
|---|---|---|---|---|
| **A)** | 0.032 mg-$NH_3$/L \| 0.362 mg-$NH_3$/L | ** | *** | ** |
| | 0.032 mg-$NH_3$/L \| 2.21 mg-$NH_3$/L | *** | *** | ns |
| | 0.362 mg-$NH_3$/L \| 2.21 mg-$NH_3$/L | ** | ** | ** |
| **B)** | **Ammonia tolerance of *Nitrospira*** | **Nitrification** | **Nitritation** | **PN/A** |
| | 0.032 mg-$NH_3$/L \| 0.362 mg-$NH_3$/L | ns | ns | ns |
| | 0.032 mg-$NH_3$/L \| 2.21 mg-$NH_3$/L | ns | ns | ns |
| | 0.362 mg-$NH_3$/L \| 2.21 mg-$NH_3$/L | ns | ns | * |
| **C)** | **Ammonia tolerance of *Nitrospira*** | **Nitrification** | **Nitritation** | **PN/A** |
| | 0.032 mg-$NH_3$/L \| 0.362 mg-$NH_3$/L | ns | ns | ns |
| | 0.032 mg-$NH_3$/L \| 2.21 mg-$NH_3$/L | ns | ns | ns |
| | 0.362 mg-$NH_3$/L \| 2.21 mg-$NH_3$/L | ns | ns | ns |
| **D)** | **Ammonia tolerance of *Nitrospira*** | **Nitrification** | **Nitritation** | **PN/A** |
| | 0.032 mg-$NH_3$/L \| 0.362 mg-$NH_3$/L | ** | ** | *** |
| | 0.032 mg-$NH_3$/L \| 2.21 mg-$NH_3$/L | ** | ** | * |
| | 0.362 mg-$NH_3$/L \| 2.21 mg-$NH_3$/L | *** | * | *** |
| **E)** | **Ammonia tolerance of *Nitrospira*** | **Nitrification** | **Nitritation** | **PN/A** |
| | 0.032 mg-$NH_3$/L \| 0.362 mg-$NH_3$/L | ns | ns | ns |
| | 0.032 mg-$NH_3$/L \| 2.21 mg-$NH_3$/L | ns | ns | ns |
| | 0.362 mg-$NH_3$/L \| 2.21 mg-$NH_3$/L | ns | ns | * |
| **F)** | **Ammonia tolerance of *Nitrospira*** | **Nitrification** | **Nitritation** | **PN/A** |
| | 0.032 mg-$NH_3$/L \| 0.362 mg-$NH_3$/L | ns | ns | ns |
| | 0.032 mg-$NH_3$/L \| 2.21 mg-$NH_3$/L | ns | ns | ns |
| | 0.362 mg-$NH_3$/L \| 2.21 mg-$NH_3$/L | * | ns | ns |
| **G)** | **Ammonia tolerance of *Nitrospira*** | **Nitrification** | **Nitritation** | **PN/A** |
| | 0.032 mg-$NH_3$/L \| 0.362 mg-$NH_3$/L | *** | *** | *** |
| | 0.032 mg-$NH_3$/L \| 2.21 mg-$NH_3$/L | *** | *** | ** |
| | 0.362 mg-$NH_3$/L \| 2.21 mg-$NH_3$/L | ns | ns | ns |
| **H)** | **Ammonia tolerance of *Nitrospira*** | **Nitrification** | **Nitritation** | **PN/A** |
| | 0.032 mg-$NH_3$/L \| 0.362 mg-$NH_3$/L | ns | * | * |
| | 0.032 mg-$NH_3$/L \| 2.21 mg-$NH_3$/L | ns | ns | ns |
| | 0.362 mg-$NH_3$/L \| 2.21 mg-$NH_3$/L | ns | ns | ns |
| **I)** | **Ammonia tolerance of *Nitrospira*** | **Nitrification** | **Nitritation** | **PN/A** |
| | 0.032 mg-$NH_3$/L \| 0.362 mg-$NH_3$/L | ns | ns | ns |
| | 0.032 mg-$NH_3$/L \| 2.21 mg-$NH_3$/L | ns | ns | ns |
| | 0.362 mg-$NH_3$/L \| 2.21 mg-$NH_3$/L | ns | ns | ns |
| **J)** | **Ammonia tolerance of *Nitrospira*** | **Nitrification** | **Nitritation** | **PN/A** |
| | 0.032 mg-$NH_3$/L \| 0.362 mg-$NH_3$/L | * | * | ** |
| | 0.032 mg-$NH_3$/L \| 2.21 mg-$NH_3$/L | * | * | ** |
| | 0.362 mg-$NH_3$/L \| 2.21 mg-$NH_3$/L | ns | ns | ns |
| **K)** | **Ammonia tolerance of *Nitrospira*** | **Nitrification** | **Nitritation** | **PN/A** |
| | 0.032 mg-$NH_3$/L \| 0.362 mg-$NH_3$/L | ns | ns | * |
| | 0.032 mg-$NH_3$/L \| 2.21 mg-$NH_3$/L | * | * | ** |
| | 0.362 mg-$NH_3$/L \| 2.21 mg-$NH_3$/L | ns | ns | ns |
| **L)** | **Ammonia tolerance of *Nitrospira*** | **Nitrification** | **Nitritation** | **PN/A** |
| | 0.032 mg-$NH_3$/L \| 0.362 mg-$NH_3$/L | ** | * | ns |
| | 0.032 mg-$NH_3$/L \| 2.21 mg-$NH_3$/L | * | * | ns |
| | 0.362 mg-$NH_3$/L \| 2.21 mg-$NH_3$/L | ** | ** | ns |
