## Supplementary material for "Physiological diversity and the assembly of nitrifying communities in aerobic granular sludge": S5 Table

**S5 Table. Statistical significance (*p*-value) from comparison of nitrogen process partitioning considering different oxygen affinity of *Nitrospira* ($K_I$, mol-$O_2$/L) A)** [$O_2$] = 0.25 mg/L; 10°C. **B)** [$O_2$] = 0.25 mg/L; 20°C. **C)** [$O_2$] = 0.25 mg/L; 30°C. **D)** [$O_2$] = 0.5 mg/L; 10°C. **E)** [$O_2$] = 0.5 mg/L; 20°C. **F)** [$O_2$] = 0.5 mg/L; 30°C. **G)** [$O_2$] = 1.0 mg/L; 10°C. **H)** [$O_2$] = 1.0 mg/L; 20°C. **I)** [$O_2$] = 1.0 mg/L; 30°C. **J)** [$O_2$] = 3.0 mg/L; 10°C. **K)** [$O_2$] = 3.0 mg/L; 20°C. **L)** [$O_2$] = 3.0 mg/L; 30°C. Symbol legend: ns – not significant; * – $p < 0.05$; ** – $p < 0.01$; *** – $p < 0.001$.

| | Oxygen affinity of *Nitrospira* | Nitrification | Nitritation | PN/A |
|---|---|---|---|---|
| A) | $5.00 \times 10^{-6}$ mol-$O_2$/L \| $3.61 \times 10^{-6}$ mol-$O_2$/L | *** | *** | *** |
| | $5.00 \times 10^{-6}$ mol-$O_2$/L \| $2.59 \times 10^{-6}$ mol-$O_2$/L | *** | *** | * |
| | $3.61 \times 10^{-6}$ mol-$O_2$/L \| $2.59 \times 10^{-6}$ mol-$O_2$/L | ** | * | * |
| B) | **Oxygen affinity of *Nitrospira*** | **Nitrification** | **Nitritation** | **PN/A** |
| | $5.00 \times 10^{-6}$ mol-$O_2$/L \| $3.61 \times 10^{-6}$ mol-$O_2$/L | ns | ns | ns |
| | $5.00 \times 10^{-6}$ mol-$O_2$/L \| $2.59 \times 10^{-6}$ mol-$O_2$/L | ns | ns | ns |
| | $3.61 \times 10^{-6}$ mol-$O_2$/L \| $2.59 \times 10^{-6}$ mol-$O_2$/L | ns | ns | ns |
| C) | **Oxygen affinity of *Nitrospira*** | **Nitrification** | **Nitritation** | **PN/A** |
| | $5.00 \times 10^{-6}$ mol-$O_2$/L \| $3.61 \times 10^{-6}$ mol-$O_2$/L | ns | ns | ns |
| | $5.00 \times 10^{-6}$ mol-$O_2$/L \| $2.59 \times 10^{-6}$ mol-$O_2$/L | ns | ns | ns |
| | $3.61 \times 10^{-6}$ mol-$O_2$/L \| $2.59 \times 10^{-6}$ mol-$O_2$/L | ns | ns | ns |
| D) | **Oxygen affinity of *Nitrospira*** | **Nitrification** | **Nitritation** | **PN/A** |
| | $5.00 \times 10^{-6}$ mol-$O_2$/L \| $3.61 \times 10^{-6}$ mol-$O_2$/L | * | * | ns |
| | $5.00 \times 10^{-6}$ mol-$O_2$/L \| $2.59 \times 10^{-6}$ mol-$O_2$/L | * | * | ns |
| | $3.61 \times 10^{-6}$ mol-$O_2$/L \| $2.59 \times 10^{-6}$ mol-$O_2$/L | * | * | ns |
| E) | **Oxygen affinity of *Nitrospira*** | **Nitrification** | **Nitritation** | **PN/A** |
| | $5.00 \times 10^{-6}$ mol-$O_2$/L \| $3.61 \times 10^{-6}$ mol-$O_2$/L | ns | ns | ns |
| | $5.00 \times 10^{-6}$ mol-$O_2$/L \| $2.59 \times 10^{-6}$ mol-$O_2$/L | ns | ns | * |
| | $3.61 \times 10^{-6}$ mol-$O_2$/L \| $2.59 \times 10^{-6}$ mol-$O_2$/L | ns | ns | * |
| F) | **Oxygen affinity of *Nitrospira*** | **Nitrification** | **Nitritation** | **PN/A** |
| | $5.00 \times 10^{-6}$ mol-$O_2$/L \| $3.61 \times 10^{-6}$ mol-$O_2$/L | ns | ns | ns |
| | $5.00 \times 10^{-6}$ mol-$O_2$/L \| $2.59 \times 10^{-6}$ mol-$O_2$/L | ns | ns | ns |
| | $3.61 \times 10^{-6}$ mol-$O_2$/L \| $2.59 \times 10^{-6}$ mol-$O_2$/L | * | ns | ns |
| G) | **Oxygen affinity of *Nitrospira*** | **Nitrification** | **Nitritation** | **PN/A** |
| | $5.00 \times 10^{-6}$ mol-$O_2$/L \| $3.61 \times 10^{-6}$ mol-$O_2$/L | ** | ** | ns |
| | $5.00 \times 10^{-6}$ mol-$O_2$/L \| $2.59 \times 10^{-6}$ mol-$O_2$/L | ns | ns | ns |
| | $3.61 \times 10^{-6}$ mol-$O_2$/L \| $2.59 \times 10^{-6}$ mol-$O_2$/L | ns | ns | ns |
| H) | **Oxygen affinity of *Nitrospira*** | **Nitrification** | **Nitritation** | **PN/A** |
| | $5.00 \times 10^{-6}$ mol-$O_2$/L \| $3.61 \times 10^{-6}$ mol-$O_2$/L | ns | ns | ns |
| | $5.00 \times 10^{-6}$ mol-$O_2$/L \| $2.59 \times 10^{-6}$ mol-$O_2$/L | ns | ns | ns |
| | $3.61 \times 10^{-6}$ mol-$O_2$/L \| $2.59 \times 10^{-6}$ mol-$O_2$/L | ns | ns | ns |
| I) | **Oxygen affinity of *Nitrospira*** | **Nitrification** | **Nitritation** | **PN/A** |
| | $5.00 \times 10^{-6}$ mol-$O_2$/L \| $3.61 \times 10^{-6}$ mol-$O_2$/L | ns | ns | ns |
| | $5.00 \times 10^{-6}$ mol-$O_2$/L \| $2.59 \times 10^{-6}$ mol-$O_2$/L | ns | ns | ns |
| | $3.61 \times 10^{-6}$ mol-$O_2$/L \| $2.59 \times 10^{-6}$ mol-$O_2$/L | ns | ns | ns |
| J) | **Oxygen affinity of *Nitrospira*** | **Nitrification** | **Nitritation** | **PN/A** |
| | $5.00 \times 10^{-6}$ mol-$O_2$/L \| $3.61 \times 10^{-6}$ mol-$O_2$/L | ns | ns | ns |
| | $5.00 \times 10^{-6}$ mol-$O_2$/L \| $2.59 \times 10^{-6}$ mol-$O_2$/L | ns | ns | ns |
| | $3.61 \times 10^{-6}$ mol-$O_2$/L \| $2.59 \times 10^{-6}$ mol-$O_2$/L | ns | ns | ns |
| K) | **Oxygen affinity of *Nitrospira*** | **Nitrification** | **Nitritation** | **PN/A** |
| | $5.00 \times 10^{-6}$ mol-$O_2$/L \| $3.61 \times 10^{-6}$ mol-$O_2$/L | ns | ns | ns |
| | $5.00 \times 10^{-6}$ mol-$O_2$/L \| $2.59 \times 10^{-6}$ mol-$O_2$/L | ns | ns | ns |
| | $3.61 \times 10^{-6}$ mol-$O_2$/L \| $2.59 \times 10^{-6}$ mol-$O_2$/L | ns | ns | ns |
| L) | **Oxygen affinity of *Nitrospira*** | **Nitrification** | **Nitritation** | **PN/A** |
| | $5.00 \times 10^{-6}$ mol-$O_2$/L \| $3.61 \times 10^{-6}$ mol-$O_2$/L | ns | ns | ns |
| | $5.00 \times 10^{-6}$ mol-$O_2$/L \| $2.59 \times 10^{-6}$ mol-$O_2$/L | * | * | ns |
| | $3.61 \times 10^{-6}$ mol-$O_2$/L \| $2.59 \times 10^{-6}$ mol-$O_2$/L | * | * | ns |
