## Supplementary material for "Physiological diversity and the assembly of nitrifying communities in aerobic granular sludge": S6 Table

**S6 Table. Influence of ammonia tolerance of *Nitrospira* ($K_I$) on maximum portion values of nitrogen conversion processes.** Symbol legend: NF – nitrification, NI – nitritation, PN/A – partial nitritation + anammox. Standard deviation from three replicates.

| | 10°C | 20°C | 30°C |
|---|---|---|---|
| 0.25 mg-$O_2$/L | $K_I$ = 0.032 mg/L<br>NF: 91.61% ± 0.15%<br>NI: 92.64% ± 0.43%<br>PN/A: 6.42% ± 0.17% | $K_I$ = 0.032 mg/L<br>NF: 73.92% ± 0.85%<br>NI: 70.96% ± 0.41%<br>PN/A: 27.10% ± 1.15% | $K_I$ = 0.032 mg/L<br>NF: 42.74% ± 0.04%<br>NI: 33.51% ± 1.37%<br>PN/A: 72.65% ± 0.50% |
| | $K_I$ = 0.362 mg/L<br>NF: 94.20% ± 2.84%<br>NI: 46.84% ± 1.28%<br>PN/A: 6.71% ± 0.22% | $K_I$ = 0.362 mg/L<br>NF: 79.37% ± 2.58%<br>NI: 72.25% ± 1.02%<br>PN/A: 26.54% ± 0.18% | $K_I$ = 0.362 mg/L<br>NF: 48.99% ± 2.40%<br>NI: 36.16% ± 2.75%<br>PN/A: 72.17% ± 1.50% |
| 0.5 mg-$O_2$/L | $K_I$ = 0.032 mg/L<br>NF: 95.68% ± 0.37%<br>NI: 62.68% ± 2.45%<br>PN/A: 3.41% ± 0.08% | $K_I$ = 0.032 mg/L<br>NF: 77.35% ± 2.50%<br>NI: 87.89% ± 0.18%<br>PN/A: 22.11% ± 0.47% | $K_I$ = 0.032 mg/L<br>NF: 38.89% ± 4.83%<br>NI: 71.75% ± 2.10%<br>PN/A: 60.03% ± 1.13% |
| | $K_I$ = 0.362 mg/L<br>NF: 97.53% ± 0.02%<br>NI: 23.51% ± 1.69%<br>PN/A: 4.97% ± 0.19% | $K_I$ = 0.362 mg/L<br>NF: 85.63% ± 1.28%<br>NI: 87.69% ± 0.24%<br>PN/A: 21.06% ± 1.45% | $K_I$ = 0.362 mg/L<br>NF: 42.40% ± 3.16%<br>NI: 74.72% ± 1.06%<br>PN/A: 59.04% ± 0.49% |
| 1.0 mg-$O_2$/L | $K_I$ = 0.032 mg/L<br>NF: 97.33% ± 0.10%<br>NI: 14.01% ± 0.77%<br>PN/A: 1.47% ± 0.01% | $K_I$ = 0.032 mg/L<br>NF: 79.41% ± 2.56%<br>NI: 94.49% ± 0.05%<br>PN/A: 9.90% ± 0.10% | $K_I$ = 0.032 mg/L<br>NF: 26.33 ± 1.55%<br>NI: 84.18% ± 1.14%<br>PN/A: 54.67% ± 0.69% |
| | $K_I$ = 0.362 mg/L<br>NF: 98.62% ± 0.02%<br>NI: 1.05% ± 0.03%<br>PN/A: 1.47% ± 0.01% | $K_I$ = 0.362 mg/L<br>NF: 88.05% ± 2.51%<br>NI: 94.34% ± 0.07%<br>PN/A: 10.29% ± 0.47% | $K_I$ = 0.362 mg/L<br>NF: 30.06% ± 3.46%<br>NI: 82.72% ± 1.45%<br>PN/A: 55.98% ± 1.03% |
| 3.0 mg-$O_2$/L | $K_I$ = 0.032 mg/L<br>NF: 97.72% ± 0.07%<br>NI: 3.04% ± 0.11%<br>PN/A: 1.27% ± 0.00% | $K_I$ = 0.032 mg/L<br>NF: 95.03% ± 0.18%<br>NI: 39.91% ± 6.08%<br>PN/A: 1.27% ± 0.00% | $K_I$ = 0.032 mg/L<br>NF: 41.58% ± 7.26%<br>NI: 98.21% ± 0.12%<br>PN/A: 7.82% ± 1.33% |
| | $K_I$ = 0.362 mg/L<br>NF: 98.73% ± 0.00%<br>NI: 1.40% ± 0.30%<br>PN/A: 1.28% ± 0.00% | $K_I$ = 0.362 mg/L<br>NF: 97.09% ± 0.12%<br>NI: 55.95% ± 1.11%<br>PN/A: 1.34% ± 0.09% | $K_I$ = 0.362 mg/L<br>NF: 46.75% ± 10.07%<br>NI: 97.49% ± 0.11%<br>PN/A: 5.73% ± 0.30% |
