## Supplementary material for "Physiological diversity and the assembly of nitrifying communities in aerobic granular sludge": S7 Table

**S7 Table. Influence of oxygen affinity of *Nitrospira* ($K_{O2}$) on maximum portion values of nitrogen conversion processes.** Symbol legend: NF – nitrification, NI – nitritation, PN/A – partial nitritation + anammox. Standard deviation from three replicates.

| | 10°C | 20°C | 30°C |
|---|---|---|---|
| 0.25 mg-$O_2$/L | $K_{O2}$ = 5.00x$10^{-6}$ M<br>NF: 94.20% ± 0.20%<br>NI: 91.00% ± 0.30%<br>PN/A: 6.13% ± 0.17% | $K_{O2}$ = 5.00x$10^{-6}$ M<br>NF: 78.39% ± 1.44%<br>NI: 71.32% ± 0.98%<br>PN/A: 27.11% ± 0.18% | $K_{O2}$ = 5.00x$10^{-6}$ M<br>NF: 52.17% ± 4.12%<br>NI: 31.67% ± 0.71%<br>PN/A: 72.37% ± 0.63% |
| | $K_{O2}$ = 3.61x$10^{-6}$ M<br>NF: 94.37% ± 0.23%<br>NI: 4.53% ± 0.88%<br>PN/A: 6.20% ± 0.13% | $K_{O2}$ = 3.61x$10^{-6}$ M<br>NF: 81.15% ± 0.59%<br>NI: 68.47% ± 2.56%<br>PN/A: 28.24% ± 1.02% | $K_{O2}$ = 3.61x$10^{-6}$ M<br>NF: 53.69% ± 1.45%<br>NI: 33.00% ± 0.26%<br>PN/A: 72.71% ± 2.07% |
| 0.5 mg-$O_2$/L | $K_{O2}$ = 5.00x$10^{-6}$ M<br>NF: 97.83% ± 0.12%<br>NI: 46.13% ± 13.13%<br>PN/A: 4.41% ± 0.05% | $K_{O2}$ = 5.00x$10^{-6}$ M<br>NF: 88.02% ± 0.37%<br>NI: 87.24% ± 0.43%<br>PN/A: 22.47% ± 0.31% | $K_{O2}$ = 5.00x$10^{-6}$ M<br>NF: 45.28% ± 4.62%<br>NI: 73.75% ± 1.86%<br>PN/A: 61.82% ± 1.45% |
| | $K_{O2}$ = 3.61x$10^{-6}$ M<br>NF: 97.50% ± 0.06%<br>NI: 2.15% ± 0.93%<br>PN/A: 5.14% ± 0.29% | $K_{O2}$ = 3.61x$10^{-6}$ M<br>NF: 88.48% ± 1.00%<br>NI: 87.91% ± 0.76%<br>PN/A: 21.66% ± 1.45% | $K_{O2}$ = 3.61x$10^{-6}$ M<br>NF: 58.71% ± 3.86%<br>NI: 73.09% ± 0.69%<br>PN/A: 61.98% ± 1.85% |
| 1.0 mg-$O_2$/L | $K_{O2}$ = 5.00x$10^{-6}$ M<br>NF: 98.48% ± 0.01%<br>NI: 0.88% ± 0.09%<br>PN/A: 1.47% ± 0.02% | $K_{O2}$ = 5.00x$10^{-6}$ M<br>NF: 91.85% ± 1.54%<br>NI: 94.60% ± 0.08%<br>PN/A: 10.28% ± 0.60% | $K_{O2}$ = 5.00x$10^{-6}$ M<br>NF: 32.22% ± 5.46%<br>NI: 82.72% ± 2.43%<br>PN/A: 54.87% ± 1.37% |
| | $K_{O2}$ = 3.61x$10^{-6}$ M<br>NF: 98.59% ± 0.05%<br>NI: 0.64% ± 0.09%<br>PN/A: 1.44% ± 0.02% | $K_{O2}$ = 3.61x$10^{-6}$ M<br>NF: 91.85% ± 2.02%<br>NI: 94.60% ± 0.08%<br>PN/A: 10.28% ± 0.60% | $K_{O2}$ = 3.61x$10^{-6}$ M<br>NF: 35.95% ± 4.77%<br>NI: 82.65% ± 0.53%<br>PN/A: 55.03% ± 1.96% |
| 3.0 mg-$O_2$/L | $K_{O2}$ = 5.00x$10^{-6}$ M<br>NF: 98.64% ± 0.09%<br>NI: 1.11% ± 0.42%<br>PN/A: 1.28% ± 0.00% | $K_{O2}$ = 5.00x$10^{-6}$ M<br>NF: 98.38% ± 0.60%<br>NI: 64.27% ± 5.66%<br>PN/A: 1.42% ± 0.16% | $K_{O2}$ = 5.00x$10^{-6}$ M<br>NF: 46.04% ± 6.53%<br>NI: 97.82% ± 0.15%<br>PN/A: 5.57% ± 0.21% |
| | $K_{O2}$ = 3.61x$10^{-6}$ M<br>NF: 98.64% ± 0.09%<br>NI: 1.04% ± 0.41%<br>PN/A: 1.29% ± 0.01% | $K_{O2}$ = 3.61x$10^{-6}$ M<br>NF: 97.81% ± 0.66%<br>NI: 54.60% ± 14.51%<br>PN/A: 1.40% ± 0.14% | $K_{O2}$ = 3.61x$10^{-6}$ M<br>NF: 43.85% ± 2.34%<br>NI: 97.47% ± 0.15%<br>PN/A: 5.05% ± 0.56% |
