## Supplementary material for "Physiological diversity and the assembly of nitrifying communities in aerobic granular sludge": S8 Table

**S8 Table. Metabolic stoichiometries of AOB, NOB (*Nitrobacter*, *Nitrospira* and *Nitrotoga*) and AMX.** *Cat* and *Anab* refer to catabolism and anabolism, respectively.

| | AOB | | | NOB | | | AMX | | |
|---|---|---|---|---|---|---|---|---|---|
| | ***Cat*** | ***Anab*** | ***Decay*** | ***Cat*** | ***Anab*** | ***Decay*** | ***Cat*** | ***Anab*** | ***Decay*** |
| $NH_3$ | -1.0 | -0.9 | 0 | 0 | 0 | 0 | -1.0 | -0.2 | 0 |
| $NO_2^-$ | 1.0 | 0.7 | 0 | -1.0 | -2.9 | 0 | -1.0 | -2.1 | 0 |
| $NO_3^-$ | 0 | 0 | 0 | 1.0 | 2.7 | 0 | 0 | 2.1 | 0 |
| $O_2$ | -1.5 | 0 | 0 | -0.5 | 0 | 0 | 0 | 0 | 0 |
| $CO_2$ | 0 | -1.0 | 1.0 | 0 | -1.0 | 1.0 | 0 | -1.0 | 1.0 |
| $N_2$ | 0 | 0 | 0 | 0 | 0 | 0 | 1.0 | 0 | 0 |
| $H_2O$ | 1.0 | 1.1 | 0 | 0 | 0.2 | 0 | 2.0 | 0.4 | 0 |
| $H^+$ | 1.0 | -1.0 | 0 | 0 | -1.0 | 0 | 0 | -1.0 | 0 |
