## Supplementary material for "Physiological diversity and the assembly of nitrifying communities in aerobic granular sludge": S9 Table

**S9 Table. List of general parameters used in all simulation experiments.** EST – estimated.

| Parameter | Symbol | Value | Unit | REF |
|---|---|---|---|---|
| **Simulation domain** | | | | |
| Domain size | $L_x \times L_y$ | $1028 \times 1028$ | μm | – |
| Cartesian grid cells | $N_x \times N_y$ | $257 \times 257$ | grid cells | – |
| Grid cell size | $\Delta x \times \Delta y$ | $4 \times 4$ | μm | – |
| Size of initial aggregates (radius) | $r_{init}$ | $50$ | μm | – |
| Minimum *dt* diffusion | $\min(dt)$ | $1.51 \cdot 10^{-8}$ | h | – |
| Maximum *dt* diffusion | $\max(dt)$ | $1.21 \cdot 10^{-6}$ | h | – |
| Minimum *dt* bacteria | $\min(dt_{bac})$ | $0.05$ | h | – |
| Maximum *dt* bacteria | $\max(dt_{bac})$ | $1.0$ | h | – |
| **Physical parameters: diffusion** | | | | |
| Ammonia ($NH_3$) | $\mathbb{D}_{NH_3}$ | $7.05 \cdot 10^{-6}$ | $m^2\ h^{-1}$ | [1] |
| Nitrite ($NO_2^-$) | $\mathbb{D}_{NO_2^-}$ | $6.88 \cdot 10^{-6}$ | $m^2\ h^{-1}$ | [1] |
| Nitrate ($NO_3^-$) | $\mathbb{D}_{NO_3^-}$ | $6.85 \cdot 10^{-6}$ | $m^2\ h^{-1}$ | [1] |
| Oxygen ($O_2$) | $\mathbb{D}_{O_2}$ | $7.56 \cdot 10^{-6}$ | $m^2\ h^{-1}$ | [1] |
| Carbon dioxide ($CO_2$) | $\mathbb{D}_{CO_2}$ | $6.91 \cdot 10^{-6}$ | $m^2\ h^{-1}$ | [1] |
| Sulfate ($SO_4^{2-}$) | $\mathbb{D}_{SO_4^{2-}}$ | $4.99 \cdot 10^{-6}$ | $m^2\ h^{-1}$ | [1] |
| Ion sodium ($Na^+$) | $\mathbb{D}_{Na^+}$ | $4.80 \cdot 10^{-6}$ | $m^2\ h^{-1}$ | [1] |
| Diffusion correction term | $\sigma_{Diff}$ | $0.7$ | – | [2] |
| **Physical parameters of bacteria** | | | | |
| Density of biomass | $\rho_b$ | $500$ | $kg\ m^{-3}$ | [3, 4] |
| Division radius | $r_{max}$ | $1.0$ | μm | [5] |
| Inactivation radius | $r_{max}$ | $0.464$ | μm | [5] |
| Molecular weight ($CH_{1.8}O_{0.5}N_{0.2}$) | $MW_b$ | $24.6$ | $g\ mol^{-1}$ | [6] |
| Overlap distance coefficient | $kDist$ | $1.5$ | – | – |
| Global detachment rate | $kDet$ | $5.0$ | $m^{-1}\ h^{-1}$ | – |
| Density of aggregate | $\rho_{agg}$ | $100$ | $g\ L^{-1}$ | [7] |
| Density of reactor | $\rho_r$ | $3.0$ | $g\ L^{-1}$ | [8] |
| **Reactor parameters** | | | | |
| Boundary layer thickness | $L_b$ | $5.0$ | μm | [9] |
| Representative volume | $V_r$ | $4.71 \cdot 10^{-10}$ | $m^3$ | EST |
| Bulk concentration pH | $pH$ | $7.7$ | – | [10] |
| Temperature | $T$ | $10, 20, 30$ | °C | – |
| Influent concentration ammonia | $[NH_3]_{inf}$ | 40.0 | $mg\ L^{-1}$ | [10] |
| Influent concentration oxygen | $[O_2]_{inf}$ | 0.1 – 6.0 | $mg\ L^{-1}$ | [10] |
| Influent concentration pH buffer | $[CO_2]_{inf}$ | $1.00 \cdot 10^{-3}$ | $mol\ L^{-1}$ | – |
| Setpoint concentration ammonia | $[NH_3]_{SP}$ | 20.0 | $mg\ L^{-1}$ | [10] |
| Setpoint concentration oxygen | $[O_2]_{SP}$ | 0.1 – 6.0 | $mg\ L^{-1}$ | [10] |
| Hydraulic retention time | $HRT$ | Variable | h | – |
| **Computational parameters** | | | | |
| Steady-state tolerance | $tol_{SS}$ | $0.05$ | $mol\ L^{-1}\ h^{-1}$ | – |
| Diffusion tolerance | $tol_{Diff}$ | $1.00 \cdot 10^{-7}$ | $mol\ L^{-1}\ h^{-1}$ | – |
| Maximum bulk concentration change | $\Delta\phi_{bulk,max}$ | $2.0$ | % | – |
| pH solver tolerance | $tol_k$ | $1.00 \cdot 10^{-15}$ | – | – |
| Concentration tolerance | $tol_{[S]}$ | $1.00 \cdot 10^{-8}$ | $mol\ L^{-1}$ | – |
