## Supplementary material for "Physiological diversity and the assembly of nitrifying communities in aerobic granular sludge": S10 Table

### S10 Table. Expressions of growth-temperature influence on growth kinetics.

| Model | Equation* | REF |
|---|---|---|
| Arrhenius equation | $\mu_{max} = A \cdot \exp\left(-\frac{E_a}{RT}\right)$ | [1] |
| Modified Ratkowsky | $\mu_{max} = [b(T - T_{min})]^2 \cdot \{1 - \exp[c(T - T_{max})]\}$ | [2] |
| Hinshelwood model [(1)] | $\mu_{max} = k_1 \cdot \exp\left(-\frac{E_1}{R}\left(\frac{1}{T} - \frac{1}{T_{opt}}\right)\right) - k_2 \cdot \exp\left(-\frac{E_2}{R}\left(\frac{1}{T} - \frac{1}{T_{opt}}\right)\right)$ | [3] |
| Yan and Hunt model | $\mu_{max} = A_{YH}\left(\frac{T_{max}-T}{T_{max}-T_{opt}}\right)\left(\frac{T}{T_{opt}}\right)^{T_{opt}/(T_{max}-T_{opt})}$ | [4] |

[(1)] Both the enzyme reaction (first term) and the high-temperature denaturation (second term) show an Arrhenius type of temperature dependency and are zero order.

Symbol legend: $A$ – frequency factor ($h^{-1}$); $E_a$ – activation energy (J $mol^{-1}$); $R$ – gas constant (8.314 J $mol^{-1}$ $K^{-1}$); $T$ – temperature in Kelvin (K); $b$ – Ratkowsky parameter ($°C^{-1}$ $h^{-0.5}$); $c$ – Ratkowsky parameter ($°C^{-1}$); $T_{min}$, $T_{max}$ – minimum and maximum temperature at which growth is observed, respectively; $k_1$, $k_2$ – frequency actors ($h^{-1}$); $E_1$, $E_2$ – activation energies of the enzyme reaction and the high-temperature denaturation, respectively (J $mol^{-1}$); $T_{opt}$ – optimal growth temperature; $\mathrm{A_{YH}}$ – Yan and Hunt constant ($h^{-1}$).
