## Supplementary material for "Physiological diversity and the assembly of nitrifying communities in aerobic granular sludge": S11 Table

**S11 Table. Arrhenius coefficients ($\theta$) of *Nitrobacter, Nitrospira* and *Nitrotoga*.**

| Culture | θ | Temperature (°C) | pH | REF |
|---|---|---|---|---|
| ***Nitrobacter* (NOB)** | | | | |
| *Nitrobacter sp.* | 1.108 | 5.0 – 25.0 | 7.8 | [1] |
| *Nitrobacter sp.* | 1.080 | 5.0 – 23.0 | 8.1 – 8.5 | [2] |
| *Nitrobacter sp.* | 1.064 | 10.0 – 30.0 | 7.5 | [3] |
| *Nitrobacter agilis* | 1.104 | 3.0 – 30.0 | 7.8 | [4] |
| *Nitrobacter agilis* | 1.109 | 15.0 – 30.0 | 7.8 | [5] |
| *Nitrobacter winogradskyi* | 1.089 | 15.0 – 45.0 | 7.8 | [6] |
| *Nitrobacter sp.* | 1.085 | 0.0 – 35.0 | 7.5 | [7] |
| *Nitrobacter sp.* | 1.060 | 5.0 – 31.0 | 7.9 | [8] |
| **Average ± St. dev.** | **1.087 ± 0.019** | – | – | – |
| ***Nitrospira* (NOB)** | | | | |
| *Nitrospira sp.* | 1.066 | 12.0 – 35.0 | 7.35 | [9] |
| *Nitrospira defluvii* | 1.110 | 10.0 – 35.0 | 7.4 | [10] |
| *Nitrospira sp.* BS10 | 1.082 | 10.0 – 28.0 | 8.4 – 8.6 | [11] |
| **Average ± St. dev.** | **1.086 ± 0.022** | – | – | – |
| ***Nitrotoga* (NOB)** | | | | |
| *Nitrotoga fabula* | 1.203 | 5.0 – 25.0 | 7.3 | [12] |
| *Nitrotoga sp.* CP | 1.127 | 4.0 – 25.0 | 7.0 | [13] |
| *Nitrotoga sp.* AMP1 | 1.100 | 4.0 – 20.0 | 8.3 | [14] |
| *Nitrotoga sp.* HW29 | 1.095 | 4.0 – 22.0 | 6.8 | [15] |
| *Nitrotoga sp.* AM1 | 1.095 | 5.0 – 15.0 | 8.0 – 8.3 | [16] |
| **Average ± St. dev.** | **1.124 ± 0.046** | – | – | – |
