## Supplementary material for "Physiological diversity and the assembly of nitrifying communities in aerobic granular sludge": S12 Table

**S12 Table. Fitting parameters for each model ant the resulted ARD and $s_e$.**

| **Hinshelwood model** | | | | | | |
|---|---|---|---|---|---|---|
| **$k_1$ [$h^{-1}$]** | **$E_1$ [J/mol]** | **$k_2$ [$h^{-1}$]** | **$E_2$ [J/mol]** | **$T_{opt}$ [K]** | **ARD [(1)] [%]** | **$s_e$ [(2)] [–]** |
| $7.87 \times 10^{-2}$ | $1.87 \times 10^{5}$ | $6.35 \times 10^{-2}$ | $207 \times 10^{5}$ | 296.92 | 45 | $1.605 \times 10^{-3}$ |
| **Modified Ratkowsky model** | | | | | | |
| **b [°$C^{-1}$ $h^{-0.5}$]** | **$T_{min}$ [K]** | **c [°$C^{-1}$]** | **$T_{max}$ [K]** | | **ARD [%]** | **$s_e$ [–]** |
| $8.47 \times 10^{-3}$ | 281.97 | 0.417 | 305.10 | | 59 | $1.599 \times 10^{-3}$ |
| **Yan & Hunt model** | | | | | | |
| **$T_{max}$ [°C]** | **$T_{opt}$ [°C]** | **$\mu_{max}$ [$h^{-1}$]** | | | **ARD [%]** | **$s_e$ [–]** |
| 32.45 | 27.78 | 0.021 | | | 33 | $1.218 \times 10^{-3}$ |
| **Arrhenius equation** | | | | | | |
| **A [$h^{-1}$]** | **$E_A$ [J/mol]** | | | | **ARD [%]** | **$s_e$ [–]** |
| $1.53 \times 10^{22}$ | $1.35 \times 10^{5}$ | | | | 81 | $1.725 \times 10^{-2}$ |

(1) Average relative deviations: $ARD = \frac{1}{N}\sum \left| {\mu_{i,exp} - \mu_{i,calc}}/{\mu_{i,exp}} \right| \times 100$

(2) Standard error estimate: $s_e = \sqrt{\frac{1}{N-2}\sum\left(\mu_{i,exp} - \mu_{i,calc}\right)^2}$

Symbol legend: $N$ is the number of data, $\mu_{i,exp}$ the experimental value of specific growth rate from [1] and $\mu_{i,calc}$ is the calculated value of specific growth rate by mentioned models/equations.
